# Predicting pose distribution of protein domains connected by flexible linkers is an unsolved problem

**DOI:** 10.1101/2025.04.27.650885

**Authors:** Allen C. McBride, Feng Yu, Edward H. Cheng, Aulane Mpouli, Aimee C. Soe, Michal Hammel, Gaetano T. Montelione, Terrence G. Oas, Susan E. Tsutakawa, Bruce R. Donald

## Abstract

In CASP16, we assessed the ability of computational methods to predict the distribution of relative orientations of two domains tethered by a flexible linker. The range of interdomain distances and orientations (poses) of such domain-linker-domain (D-L-D) proteins can play an important role in protein function, allostery, aggregation, and the thermodynamics of binding. The CASP16 Conformational Ensembles Experiment included two challenges to predict the interdomain pose distribution of a Staphylococcal protein A (SpA) D-L-D construct, called ZLBT-C, in which two of SpA’s five nearly identical domains are connected by either (1) a six-residue wild-type (WT) linker (kadnkf), or (2) an all-glycine (Gly6) linker. The wild-type linker has a highly conserved sequence and is thought to contribute to the energetic barrier for binding with host antibodies. Ground truth was provided by nuclear magnetic resonance (NMR) residual dipolar coupling (RDC) data on WT protein and small angle X-ray scattering (SAXS) data on both proteins in solution. Twenty-five predictor groups submitted 35 sets of predicted conformational distributions, in the form of population-weighted finite ensembles of discrete structures. Unlike traditional CASP assessments that compare predicted atomic models to experimental atomic models, the accuracy of these predictions was assessed by back-calculating NMR RDCs and SAXS curves from each ensemble of atomic models and comparing these results to respective experimental data. Accuracy was also assessed by using kernelization to compare ensembles to the continuous orientational distributions optimally fit to experimental data. In our assessment, predictions spanned a wide range of accuracy, but none were close fits to the combined NMR and SAXS data. In addition, none were able to recapitulate the observed difference between WT and Gly6 proteins, as observed in the SAXS data. These results, and our analysis, highlighted strengths and weaknesses, plus complementarity of NMR RDC and SAXS analysis.

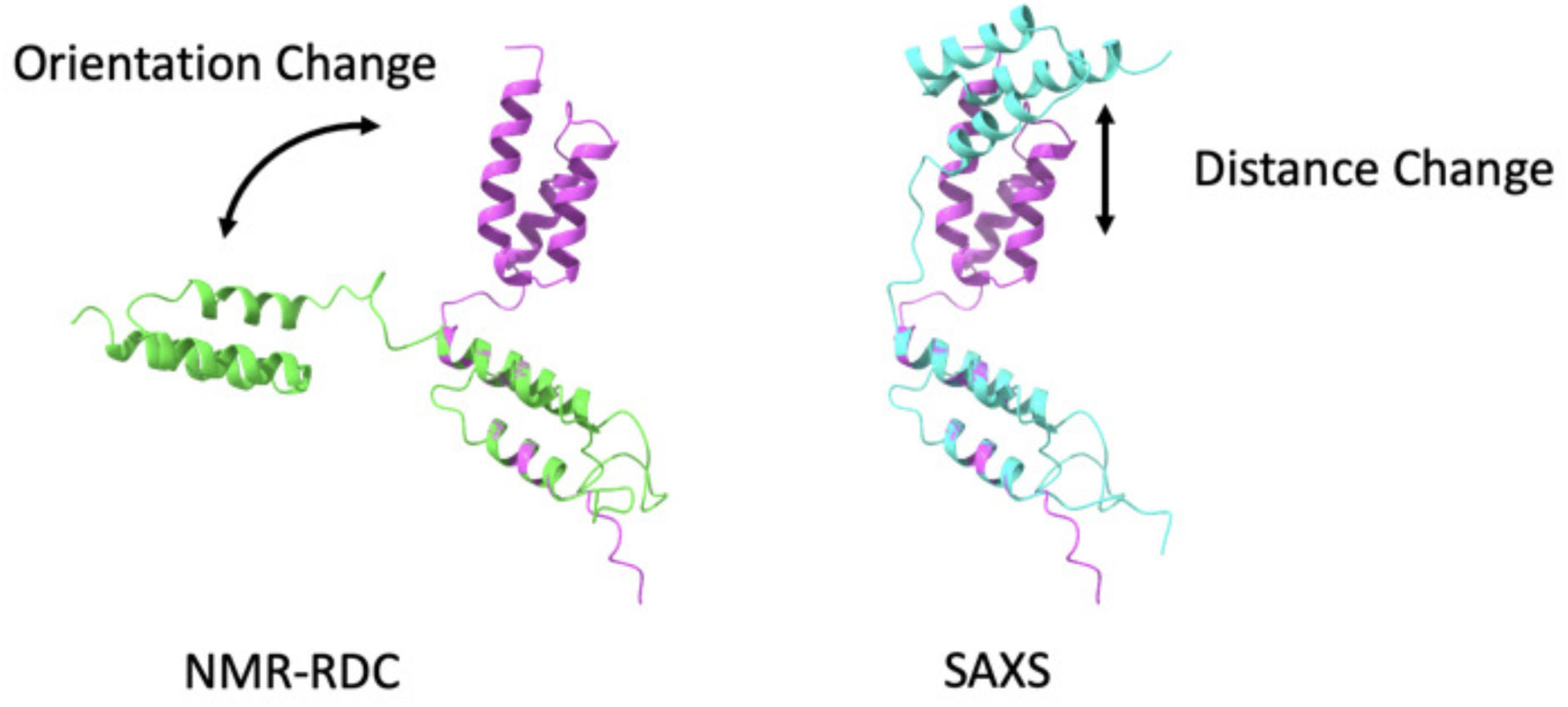

## 1 Introduction

Short, flexible sequences of amino acids often link more rigid domains, both in nature and in engineered proteins [25, 15, 53]. These linkers exist in constantly changing conformational ensembles and do not fold into a stable native structure [59]. Although linkers, like intrinsically disordered regions (IDRs) more generally, are considered as random coils to be described by polymer theory [24], they usually have constrained flexibility due to amino acid sequence constraints—both flexible yet not fully random. For such linkers, the size and shape of amino acid sidechains will constrain the conformation of the linker backbone and thereby the pose distributions of linked domains. If this pose distribution has functional significance, such as transcription activation, phase separation, or antibody binding, the sequence is likely conserved, which is atypical among flexible linkers [27, 34, 10, 47].

Characterizing and predicting not only the stiffness of linkers, but also the specific distributions of interdomain poses that they allow, is an important challenge [47, 7, 26, 1, 4]. Differences in these distributions among different linkers is determined by linker sequence as well as by the properties of the domains they connect. A better understanding of the relationship between linker sequence and interdomain pose distribution would contribute to our study of the natural biological systems containing such linkers; it would also allow us to engineer linkers to achieve desired pose distributions for improved binding in therapeutic applications or for optimized synthetic pathways in biomanufacturing.

Linked multidomain proteins belong to a larger class of proteins and protein regions possessing flexible and highly dynamic regions that do not fold into a stable native structure [59]. Such sequences longer than 30 amino acids are identified as intrinsically disordered proteins (IDPs) or IDRs [63, 66]. At least 40% of human proteins contain IDRs.

The two Critical Assessment of protein Structure Prediction (CASP) targets assessed in this report, T1200 and T1300, represent the first time that ensemble predictions have been assessed on the basis of how well each predicted ensemble, taken as a whole, fits experimental data collected from real ensembles in solution. Prior to CASP15, flexibility-related CASP challenges assessed the ability to predict which regions in a sequence were flexible (IDRs), but not the actual conformational distributions adopted by flexible regions [42, 41, 43, 11, 33, 40]. CASP15 introduced ensemble targets to CASP; however, these targets were assessed on the basis of individual models, not as conformational distributions [35].

T1200 and T1300 are two domain-linker-domain (D-L-D) proteins with the same rigid domains but different 6-amino-acid flexible linker regions. For these targets, an X-ray crystallographic structure of the two rigid domains of each construct is known and published in the Protein Data Bank (PDB) (IDs: 2LR2 [5]; 4NPD [20]). With the given sequence and reference structure of the rigid domain, predictors were instructed to predict how the linker sequence impacts the relative pose of the two well-folded protein domains by submitting an ensemble with population weights. The ensembles were to be assessed in relation to nuclear magnetic resonance (NMR) and small angle X-ray scattering (SAXS) experimental data, as well as to our previously modeled orientational distributions for one of the constructs [47], none of which predictors had access to.

The T1200 and T1300 targets are biologically relevant because they are constructs derived from protein A, which is a virulence factor in *Staphylococcus aureus*, binding to mammalian antibodies and inhibiting opsonization. Understanding the structure and function of protein A in *S. aureus* (SpA) could allow for the design of improved vaccines or therapeutics targeting *S. aureus* virulence. The N-terminal half of SpA (SpA-N) consists of five homologous three-helix domains connected by four homologous flexible linkers. These linkers are highly conserved in evolution, possibly because their constrained flexibility supports binding to antibodies such as to the Fc region of IgG [47].

The T1200 and T1300 targets correspond to two simplified D-L-D systems created to further investigate the role of these linkers in the flexibility and function of protein A, using two experimental methods (NMR and SAXS). The T1200 target is the D-L-D construct ZLBT-C. The C domain is the wild-type C domain of SpA-N; the ZLBT domain is similar to the B domain of SpA-N but includes a lanthanide binding tag (LBT) described by Barb and Subedi [6] and modified from Barb et al. [5] to facilitate partial alignment in the NMR magnetic field for purposes of residual dipolar coupling (RDC) experiments. The flexible linker between the two domains is the wild-type linker (kadnkf) in three of the four linkers of SpA-N. The T1300 target is identical to T1200 except that the linker sequence gggggg is substituted for kadnkf. The intent of this substitution is to increase flexibility [47]. The constructs for T1200 and T1300 are referred to here as “WT” and “Gly6,” respectively.

The NMR data used in this assessment comes from our prior work applying RDCs to characterize flexibility [47]. RDCs carry information about the orientation of internuclear bond vectors, such as N-H and C-H, with respect to the external magnetic field in an NMR experiment [13]. As molecules tumble and sample various conformations in solution, the angles between these bond vectors and the magnetic field change. The RDC experiment captures the average angle of these vectors over the experimental timescale, representing both time- and population-averaged molecular dynamics. This averaging reflects the distribution of interdomain orientations in the sample and can be represented as continuous probability distributions. Historically, these distributions of interdomain orientation have been represented implicitly in the form of finite, discrete, atomistic ensembles fit to RDC data [e.g. 28, 37]. In Qi et al. [47], we demonstrated benefits of representing these distributions explicitly using a family of low-dimensional continuous probability distributions fit to RDC data, termed continuous distributions of interdomain orientation (CDIOs). One limitation of RDC data is that its information is strictly orientational in nature and does not contain information on the relative probability of different interdomain translations. This limitation is partially addressed in the present study through the use of SAXS, a complementary technique that studies proteins in physiologically-relevant solution.

SAXS is a technique useful for studying flexible protein systems under relevant biological conditions [29, 12]. SAXS data contains information on the distribution of electron pair distances [46]. As the data is a sum from all molecules in solution, SAXS captures information from all conformations occurring in solution, containing information on both orientation and domain-domain distance, albeit at lower resolution then NMR RDC. For example, we showed in a CASP study that over half of the proteins adopted a different conformation in solution than in the crystal lattice [32], as was similarly observed in an NMR study [2]. This lack of agreement between solution and crystal data confounded earlier CASP attempts to use SAXS to improve prediction accuracy using crystallographic models as the ground truth. SAXS measures the intensity of X-rays scattered by the protein solution as a function of the scattering vector magnitude *q*, which is related to scattering angle *θ* and wavelength *λ* by *q* = 4*π* sin(*θ*)*/λ*. The variable *q* represents a point in reciprocal space, which is the Fourier transform of real space. By transforming reciprocal space information to real space by Fourier transformation, we can obtain a more intuitive distribution of pairwise distance for protein electrons. Importantly, SAXS reciprocal space curves can be calculated from atomic models and compared to experimental SAXS data, identifying atomic models or ensembles of atomic models that are consistent with protein solution data and can detect even small domain-domain shifts that alter multiple distances [16, 55, 17, 48]. Thus, SAXS is a useful tool in characterizing ensembles of proteins in distinct conformations.

RDC and SAXS information can be viewed as complementary (Figure 1). RDC information strictly relates to orientation in a global frame, whereas SAXS information relates to interatomic distances. SAXS also carries information about the relative orientation of the domains of a D-L-D system, because a change in this relative orientation necessarily changes the distribution of interatomic distances between the domains. However, the information is not residue specific and is convolved with the domain distances. Therefore, combined RDC and SAXS data are an attractive choice for a study whose goal is to assess the accuracy of prediction of interdomain pose distribution, where pose consists of interdomain translation and orientation.

**Figure 1:**
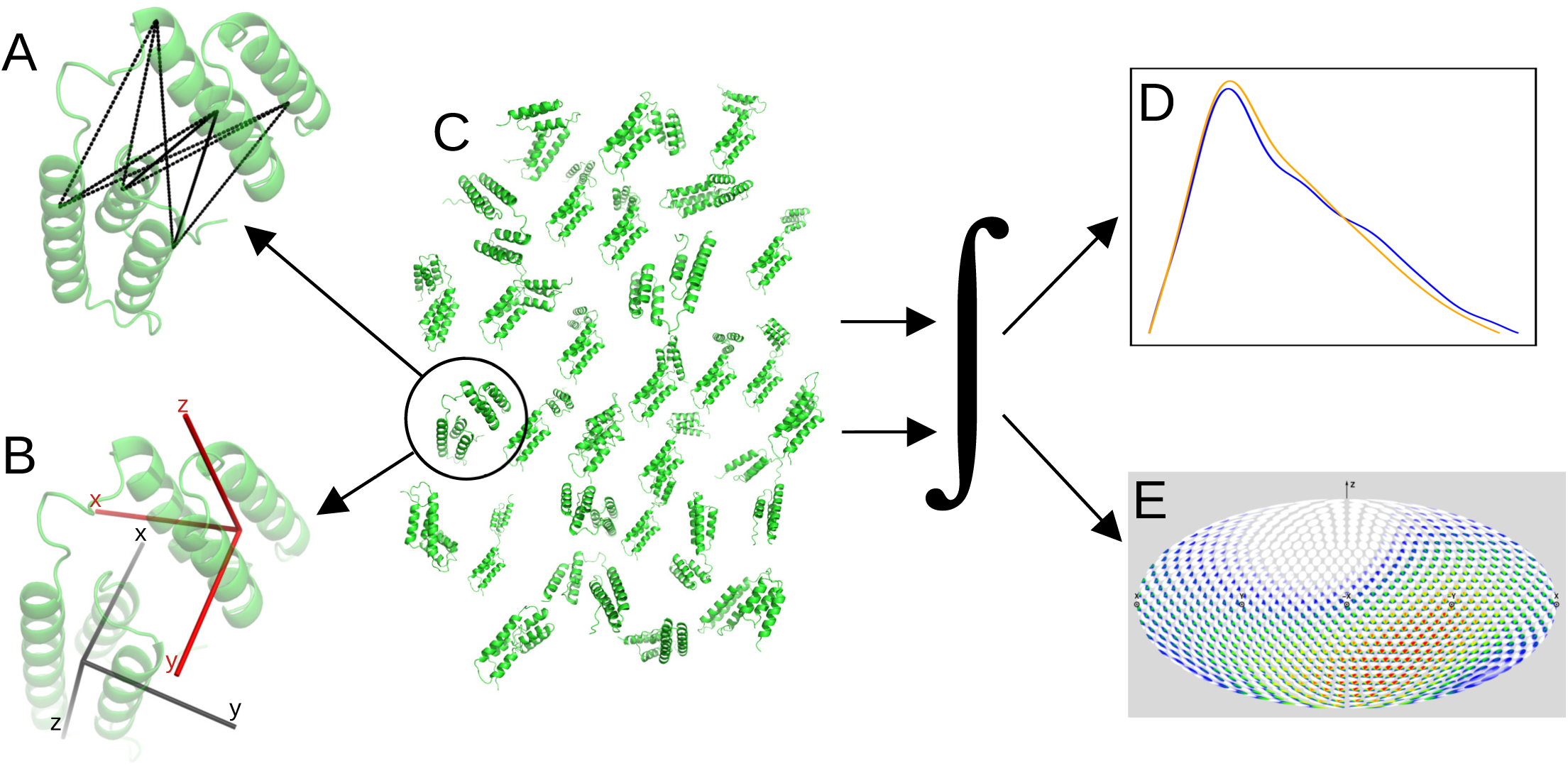
Cartoon of relationship between SAXS and NMR RDC information. For a given conformation in an ensemble (C), SAXS carries information about all pairwise interatomic distances (A), and RDCs carry information related to interdomain orientation (B). This information is integrated over the ensemble and, once analyzed, can yield models of the distributions of interatomic distances (i.e., SAXS *P* (*r*) curves, D) and interdomain orientation (i.e., CDIOs, E) for the entire ensemble. (Panels D and E are scaled-down images of Figure 4B and Supplemental Information Figure S5A. Molecule illustrations are derived from structures in ensemble 264.)

## 2 Materials and Methods

### 2.1 NMR Computational Methods

We used NMR RDC experimental data from our previous paper [47], which are deposited at BMRB (accession number 53002) [62]. The amino acid sequence of the wild-type ZLBT-C protein is mvdnkfnkeqqnafyeilhlpn lneeqrnafiqslkdyidtnndgayegdelqsanllaeakklndaqapkadnkfnkeqqnafyeilhlpnlteeqrngfiqslkd dpsvskeilaeakklndaqapk. The amino acid sequence for the Gly6 ZLBT-C variant is mvdnkfnkeqqnafyei lhlpnlneeqrnafiqslkdyidtnndgayegdelqsanllaeakklndaqapggggggnkeqqnafyeilhlpnlteeqrngfi qslkddpsvskeilaeakklndaqapk. NMR data collection for Gly6 is ongoing, so the NMR-related analysis and assessment in this experiment relates to the T1200 (WT) target.

The Saupe tensors used in our analysis are also the same as described in Qi et al. [47]. As described in that publication, experimental Saupe tensors for the ZLBT and C domains were fit to the RDCs measured in four alignment conditions for each domain, using singular value decomposition (SVD) [38, 47, 3]; SVD was then used to extract two effectively independent alignment conditions from experimental data to obtain orthogonal linear combinations (OLCs) of measured RDCs and corresponding Saupe tensors [50, 47]. The first two of these OLC Saupe tensors were used in the present analysis, and are reproduced in SI Table S5.

A branch-and-bound search algorithm was used to find CDIOs in the form of Bingham distributions that are close fits to these OLC RDCs, as described in Qi et al. [47]. Two distinct solutions were found with similar goodness-of-fit; these are qualitatively similar to the two reported in Qi et al. [47], and are named Solution 1 and Solution 2 accordingly.

Following Qi et al. [47, SI, Eq. (12)], the 4 × 4 parameter matrix *X* for a Bingham distribution can be decomposed as

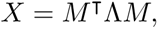

where *M* is a matrix representing a rotation in SO(4), the group of rotations in four-dimensional Euclidean space, and Λ is a diagonal matrix with constant trace specifying the variances along the four principle directions [47, SI]. *M* has 6 degrees of freedom and Λ has 3, giving *X* a total of 9 degrees of freedom [47, SI]. For Solution 1, we have

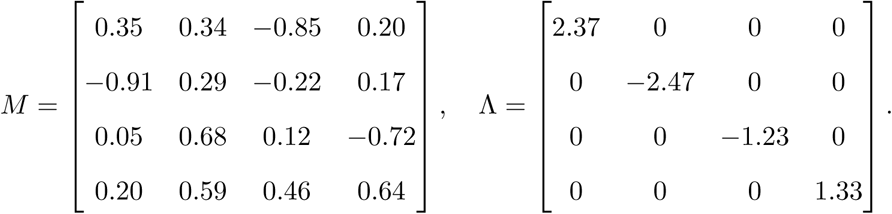

For Solution 2, we have

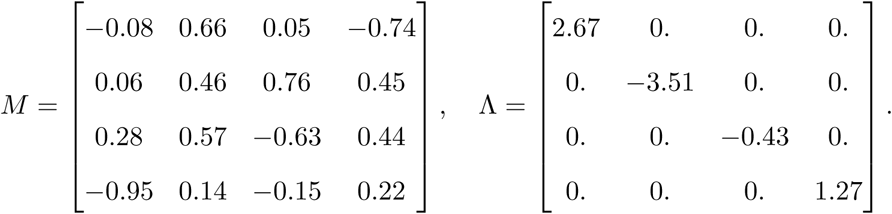

OLC RDC data back-calculated from these two solutions are a good fit for the experimental data, as shown in Figure 2. Supplemental Information (SI) Figure S5 shows these two solutions in the form of Disk-on-Sphere (DoS) visualizations, which are described in Qi et al. [47]. (The *M* matrices above are given in the coordinate system that corresponds to these DoS visualizations.)

**Figure 2:**
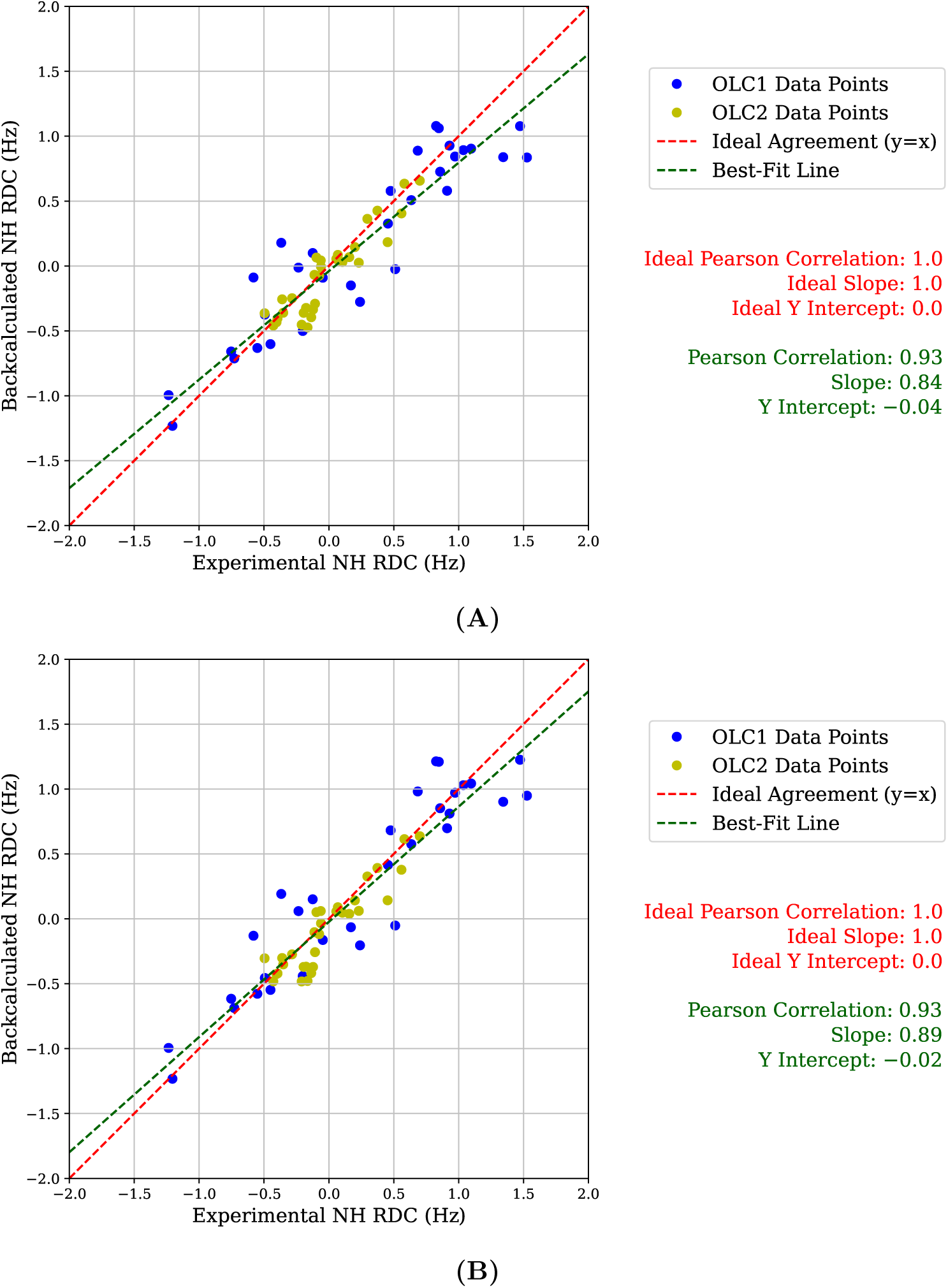
Comparing CDIO solutions to NMR RDC data. Linear regressions of OLC RDCs back-calculated from (A) Solution 1 and (B) Solution 2, which are our CDIOs optimally fit (using the algorithm in Qi et al. [47]) to our experimental NMR data, showing close fits to experimental RDCs.

If two distributions *ϕ*_1_ and *ϕ*_2_ each fit experimental RDCs, then any convex linear combination of them *αϕ*_1_ + (1 − *α*)*ϕ*_2_ is also a solution [47]. Therefore we also consider many such solutions for values of *α* between zero and one, in increments of 0.01. These solutions are not Bingham distributions, but mixtures of two Bingham distributions.

The experimental error of the RDCs is reported in Qi et al. [47] as 0.2 Hz. SI Section S6 explores implications of this uncertainty.

### 2.2 Assessment methods

A document provided to CASP predictors (see Specifications in SI) specified that backbone atoms of helices 2 and 3 of each domain be within 0.5 Å of reference models. The purpose of this requirement is to ensure meaningful analysis. Our assessment relies on attaching coordinate frames to each predicted rigid domain (Section 2.2.2), and the orientations of these frames must be comparable to the orientations described by our ground-truth CDIOs. Furthermore, our SAXS assessment assumes that differences between back-calculated and experimental data (Section 2.2.1) can be attributed to predicted linker conformations. These assumptions hold to the extent that the atoms in the relatively rigid domains of predicted structures can be aligned with reference structures.

In practice, we computed the atomic root mean square deviation (RMSD) of these backbone atoms for each domain, and then found the mean between these domains. Because the resulting distance was greater than 0.5 Å for 20 out of 35 predictors, we used a cutoff of 0.75 Å instead of 0.5 for each target. Because the goal of these CASP targets was purely to predict linker conformations, this atomic RMSD cutoff was used strictly as a filter to ensure meaningful analysis; it plays no other role in assessing the success of predictions.

#### 2.2.1 SAXS assessment

To evaluate the submitted ensembles, we use fast open-source X-ray scattering (FoXS) to convert the 3D protein structure to a theoretical scattering profile, which requires two parameters for each ensemble, *c*_1_ and *c*_2_, to estimate the possible water layer surrounding the protein [51, 52]. The water layer should be consistent for a given protein sequence and ensemble. Thus, to estimate the proper water layer parameters for the conversion, we conducted the BilboMD simulation with integrated SAXS fitting on reconstructed atomic models based on reference PDBs for both targets, and used the output parameters to preprocess all submitted ensembles [45]. We further calculated the weighted average based on the submitted population profile to obtain the predicted SAXS profile of each prediction group. Chi-squared (*χ*^2^) value is calculated between experimental and predicted data to evaluate the accuracy of the prediction, as has been described previously [51]. This *χ*^2^ metric is weighted based on the uncertainty of the experimental data for an accurate assessment. For experimental data and for data back-calculated from two example predictions, we also calculated the distribution of pairwise distances between protein electrons (*P* (*r*)), including uncertainty and maximum dimension, using the GNOM algorithm in the RAW software suite [56] (Figure 7).

#### 2.2.2 Assessing back-calculated NMR-RDC values

Our NMR-based assessments fall into two categories: those directly comparing against experimental measurements, and those involving ground-truth CDIO models. We address each in turn.

OLC RDCs (Section 2.1) for the C domain were compared to corresponding RDCs back-calculated from each predictor ensemble. To back-calculate RDCs for a given predictor ensemble, each structure in the ensemble was considered in turn. First, the orientation of each predicted domain was defined by attaching a coordinate frame to it as described in Qi et al. [47] and the Algorithm document in the SI. Then, the two reference domains (for this purpose, PDB IDs 1Q2N [67] and 4NPE [20]) were reoriented such that the relative orientation between them matched the relative orientation between the two domains of the predicted structure. That is, if the frames attached to the two domains in the predicted structure are represented as column vectors in matrices *F_PC_* and *F_PZ_* and are related by *F_PC_* = *RF_PZ_* for some rotation matrix *R*, then the two reference structures were reoriented such that *F_RC_* = *RF_RZ_*, where *F_RC_* and *F_RZ_* are, similarly, the domain-attached coordinate frames for the two reference structures. Once the reference structures were oriented in this manner, RDCs were back-calculated based on

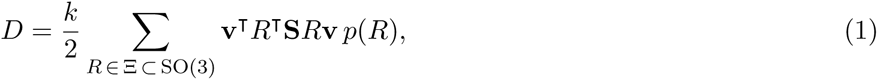

where *k* is the dipolar coupling constant (SI Section S2), Ξ is a finite sampling of SO(3), *p* : Ξ → R is a discrete probability density function (PDF), **v** is the unit bond vector, *R* is the rotation matrix defined above, and **S** is each of the two OLC Saupe tensors. *p*(*R*) is the population weight provided by predictors. When predictors did not provide population weights we assumed them to be uniform. This finite sum is a discrete approximation to the integral in SI Section S2 (SI Equation S8), in which the continuous PDF is replaced by population weights on a discrete set of orientations.

After back-calculating RDCs from a given predictor ensemble, we performed linear regression predicting back-calculated RDCs from experimental RDCs. If an ensemble were to perfectly reproduce the experimental data, then the slope of the best-fit line would be 1, the *y*-intercept would be 0, and the Pearson correlation would be 1. We considered each of these quantities separately, because each represents a different aspect in which back-calculated RDCs can be similar or dissimilar from experimental. For example, the back-calculated RDCs for predictor 331 (multicom ai) had the strongest Pearson correlation with the experimental RDCs of any predictor (*R* = 0.81); however, the slope of the best-fit line was 6.88, much higher than the slope of 1 that would result from a perfect match (Figure 3A). By contrast, the corresponding best-fit line for predictor 15 (pezyFoldings) had a slope of 0.99, the closest to 1.0 of any predictor. However, the Pearson correlation for their back-calculated RDCs was close to 0 (*R* = 0.07), indicating the near-unity slope of the best-fit line is not meaningful in the sense of signifying back-calculated RDCs that fit the experimental RDCs (Figure 3B).

**Figure 3:**
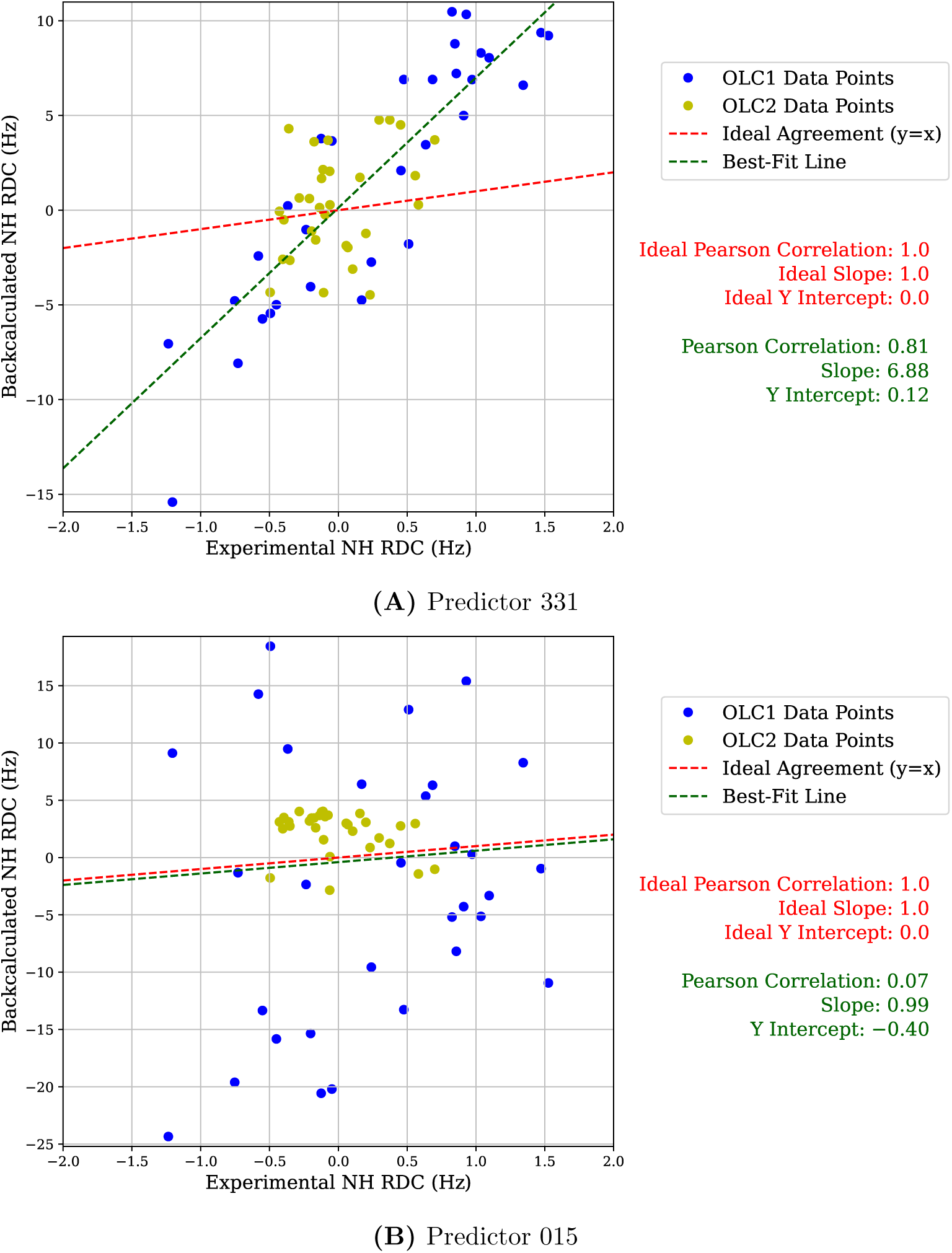
Examples of comparing experimental RDCs to RDCs back-calculated from ensemble predictions. (A) For ensemble 331, back-calculated RDCs plotted against experimental RDCs, along with the best-fit linear regression for these back-calculated RDCs as well as for the ideal case when the experimental and predicted RDCs are equal. (B) Same as above, for ensemble 015.

#### 2.2.3 Comparing predictions to CDIOs

Although predictors had the option to submit parameterized continuous distributions, all chose to submit discrete, finite ensembles of structural predictions. We interpret these submitted discrete ensembles as samples from an implied continuous distribution of conformations, which facilitates comparison with ground-truth CDIOs. Consistent with this view, we used kernel density estimation to convert each discrete ensemble into a continuous probability density function, taking a weighted sum of kernel distributions, one for each structure in a discrete ensemble. A narrow kernel function was chosen as described in SI Section S3.

To visualize these kernelized predictor ensembles in comparison with our solution CDIOs, we used the disk-on-sphere (DoS) approach we introduced in Qi et al. [47]. Instead of using a perspective projection, which requires two views per distribution (for front and rear views) and makes it hard to see densities on the edges of the projection, we used the Mollweide projection from cartography [54], originally introduced by Karl B. Mollweide in 1805. The correspondence between these projections is illustrated in SI Figure S5. This projection shows the entire sphere in one shape, and is also equal-area, simplifying the visual interpretation of different amounts of color in different parts of the sphere.

Each kernelized predictor distribution was visualized in comparison to one of the *α* values considered as convex combinations of Solution 1 and Solution 2 (Section 2.1, SI Figure S5). For visualization purposes, *α* was chosen such that, if the CDIO corresponding to that *α* were correct, the free energy difference between that solution and the given kernelized predictor distribution would be minimized. Free energy difference calculations are described in Qi et al. [47] and in SI Section S4. The distribution of these best-fit *α* values is shown in SI Figure S4.

A central concern of our work on the ZLBT-C system is to characterize how flexible the linker is, that is, in terms of relative interdomain orientation, how close to or far from isotropic its true CDIO is. In keeping with this emphasis, we assessed the accuracy of each predicted ensemble in this regard by defining “anisotropicity” and comparing the anistropicity of each predictor ensemble with the range of anisotropicities associated with our solution CDIOs for *α* values between 0 and 1. We define anisotropicity to be thermodynamically interpretable: We calculated the free energy difference (SI Section S4) between a given orientational distribution and the uniform distribution, under the provisional assumption for this purpose that the uniform distribution reflects the comparison energy field.

As our *α* parameter ranges from zero to one, this anisotropicity ranges from 0.09 to 0.42 kcal/mol. If a kernelized predictor distribution was within this range, this predictor was treated as correctly predicting isotropicity. Otherwise, each predictor was assessed based on the absolute value of the difference between their predicted anisotropicity and the nearest endpoint of this range. This approach ensures that predictors are assessed with maximum leniency by assuming their anisotropicity is correct if it is within the range of anisotropicities of our solution CDIOs. This assessment based on anisotropicity is the only element of our assessment that depends on our solution CDIO models; predictors are otherwise assessed based on comparisons between back-calculated and experimental data.

#### 2.2.4 Combining NMR-based assessments

Our assessments resulted in four values relating to NMR RDC data. Three of these relate to linear regressions of back-calculated on experimental RDCs: Pearson correlation coefficient, slope of best-fit line, and *y*-intercept of best-fit line. The fourth is our anisotropicity quantity, which relates to a free energy difference between a kernelized predicted ensemble and a uniform distribution. These four quantities must be combined for purposes of assessment. We chose to first scale each quantity according to the range of possible predicted ensembles, and to then combine them using Chebyshev distance [21]. Chebyshev distance treats the distance between two vectors as simply the maximum of the component-wise distances, so in this case, it corresponds to assessing each predicted ensemble based on its worst performance among the four quantities summarized above. This approach has two advantages: First, after scaling according to the worst that these quantities can be, comparing one quantity to another may be more meaningful. Second, because the final Chebyshev distance directly reflects only one of the underlying quantities, it does not implicitly assume that the quantities are independent. ^1^

This approach requires computing the worst that a hypothetical predictor could perform with respect to each of these quantities. Determining the worst possible Pearson correlation is not empirically straightforward, so we assume that a perfect anticorrelation (i.e., a Pearson correlation of -1) is possible. By contrast, the most extreme possible slopes and *y*-intercepts are straightforward to determine empirically. These occur when the magnitude of back-calculated RDCs is maximized, which in turn occurs when an ensemble consists of a single structure, resulting in no averaging of back-calculated RDCs. We back-calculated RDCs for single-structure ensembles whose interdomain orientations were each of 294,912 grid points on SO(3) [65]. Performing linear regression on each back-calculated RDC set against the experimental RDCs, the slope furthest from 1.0 was −14.53, and the *y*-intercept furthest from zero was 8.80.

## 3 Results

### 3.1 CASP experiment

The goal for these two CASP targets was to evaluate the prediction of the orientation and translation between the two rigid domains generated by different computational methods, as driven by the structural distribution of the six-residue flexible linker domain. The structures of the globular domains are deposited in the PDB as PDB IDs 2LR2 [5] and 4NPD [20]; predictors were given this information, in keeping with the CASP challenge to predict conformational diversity in the flexible linker, not in the rigid domains. Predictions were made using a variety of methods, mostly involving either molecular dynamics or deep learning (Table 1).

**Table 1:**
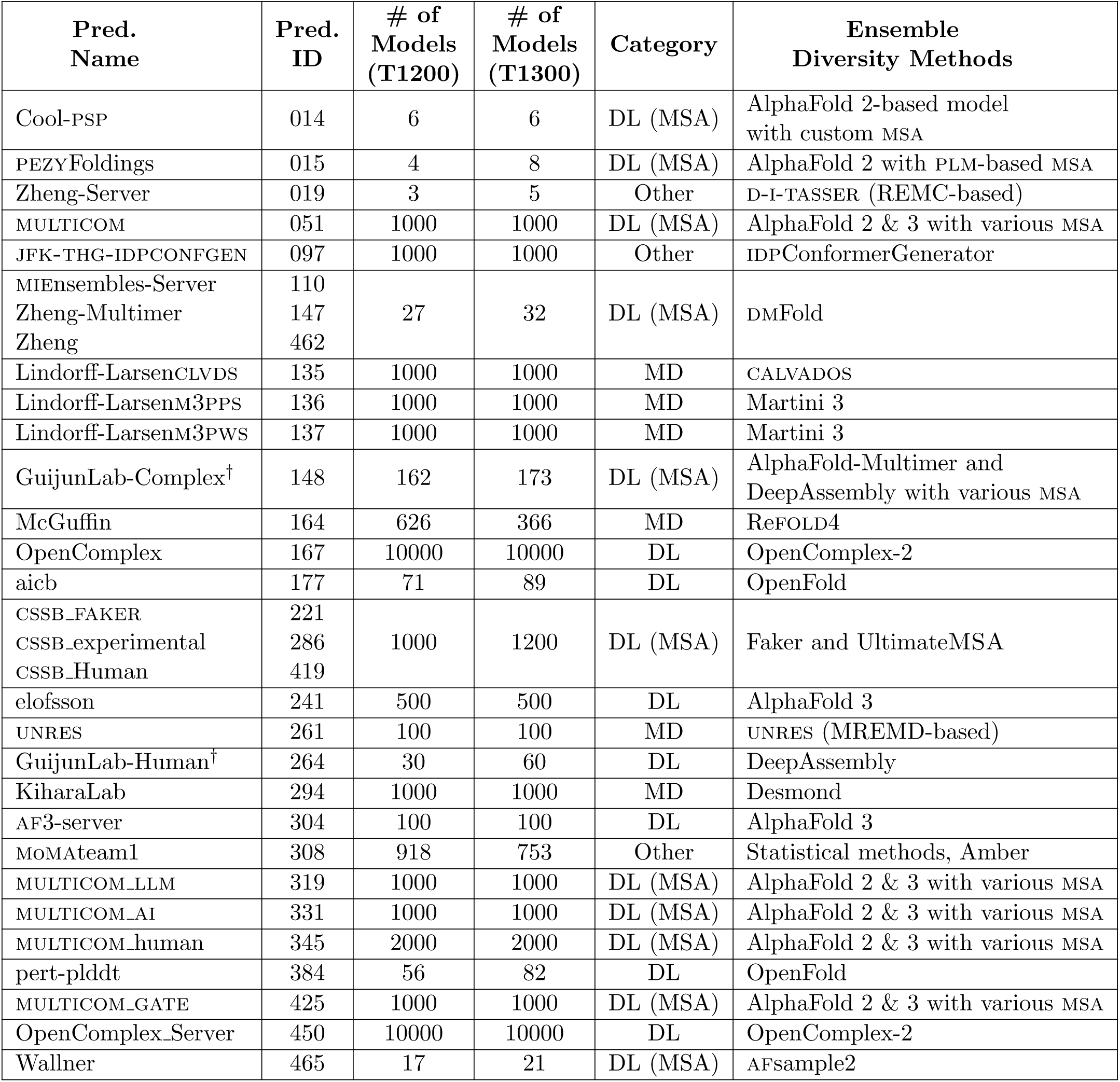
Summary of predicted ensembles meeting RMSD cutoff, including the number of atomic structural models in each ensemble and key prediction methods. This table omits information on refinement methods applied to each predicted structure in an ensemble; instead, it focuses on a key question relating to the overall shape of an ensemble: What methods were most important in generating a pool of decoys (“Ensemble diversity methods”)? For qualitative analysis purposes, we categorized predictions according to methods used to generate a pool of decoys: “DL” for deep learning with default MSA, “DL (MSA)” for deep learning with custom MSA, “MD” for molecular dynamics, and “Other” for other methods such as Monte Carlo. ^†^Ensemble 148 (target T1200) and ensemble 264 (both targets) contained mixed-sequence predictions and were not assessed; see Section 3.1.

Predictors were invited to submit predictions either in the form of discrete ensembles of PDB-format structures with accompanying population weights or in the form of Bingham orientational distributions [9, 36]. All predictors chose to submit discrete ensembles, suggesting this is more familiar for prediction algorithms. The number of discrete structures per ensemble ranged from 3 to 10,000.

Predictors were instructed (see Specifications in SI) to ensure that atomic backbone RMSDs should match the reference structures within 0.5 Å, because deviations would call into question the validity of back-calculated SAXS and RDC data, as discussed in Sections 2.2.1 and 2.2.2. However, 20 out of the 35 predicted ensembles had RMSDs above this 0.5 Å, so we filtered according to a cutoff of 0.75 Å, which reduced the number of ensembles excluded from consideration to 8, for both T1200 and T1300 targets. Excluded ensembles were numbers 2, 3, 4, 5, 84, 88, 100, and 139, with average RMSDs ranging from 0.96 to 1.75 Å (SI Table S1).

Three ensembles (148 for target T1200, and 264 for targets T1200 and T1300) each contained mixtures of structures with the WT and Gly6 sequences. The purpose of this CASP experiment was to predict conformation based on linker sequence, and our assessment presupposed that submitted structures’ sequences would match the target for which they were submitted. To ensure meaningful assessment, these mixed-sequence ensembles were therefore excluded from the assessment. Assessments made of these ensembles before this issue was recognized are found in the initial version of the preprint of this article [39].

### 3.2 SAXS

The Size-Exclusion-Coupled (SEC)-SAXS data were collected on the T1200 and T1300 systems at the SIBYLS beamline 12.3.1 at the Advanced Light Source Synchrotron [49] (details to be published in an experimental paper). The columns were equilibrated in the same buffer used for the NMR samples except for the addition of glycerol: 25 mM 3-(N-morpholino)propanesulfonic acid (MOPS) pH 7.2, 100 mM KCl, and 1% glycerol. The proteins were stoichiometrically monodisperse, based on co-collected multi-angle light scattering (MALS), SAXS-calculated molecular weight, and linear Guinier curves.

The T1200 (WT) and T1300 (Gly6) targets were similar to each other in terms of SAXS results, with the SAXS data in reciprocal and real space indicating that T1200 was more globular and T1300 more extended (Figure 4). Tellingly, the real space analysis that describes the pairwise distance distribution of protein residues revealed an important difference: the *P* (*r*) curve of T1200 had visible shoulders after the main peak, which were not as prominent in the curve for the T1300 sample (Figure 4B, arrows). While the main peak represents the intradomain distances, the peaks at a longer distance represent interdomain distances. The visible shoulders in the experimental data for T1200 suggest a population with longer preferred interdomain distances while the lack of shoulders is consistent with the loss of preferred distances in T1300. This finding supports the model that the conserved WT linker in T1200 constrains the conformational space, while the Gly6 replacement in T1300 reduces that constraint. This difference was important for assessing the ensemble predictions of each predictor.

**Figure 4:**
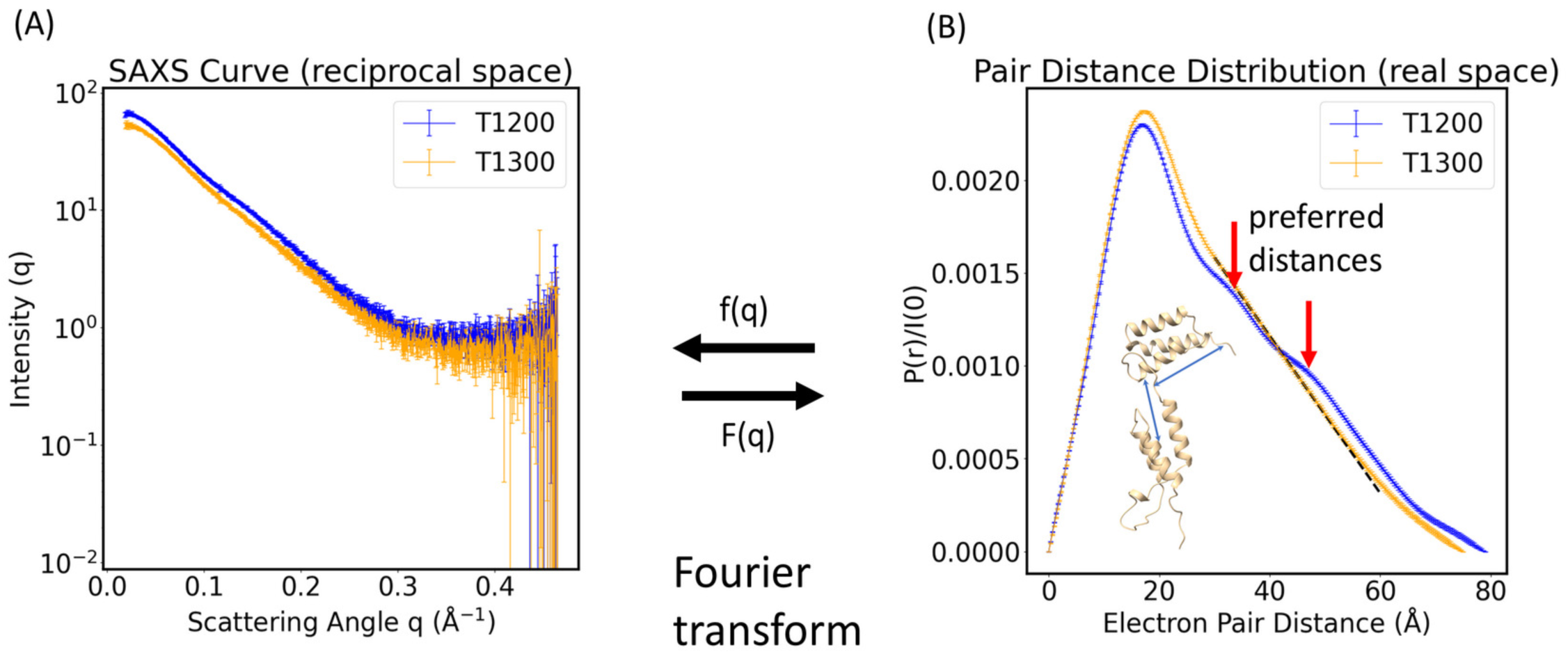
SAXS experimental data for targets T1200 and T1300. (A) The reciprocal space data for T1200 and T1300, with experimental standard deviation. (B) The real space data for T1200 and T1300, calculated from the reciprocal space data, with bars (very small) for estimated error as described in Section 2.2.1. The *y*-axis represents the normalized probability of the pair distance distribution.

Based on these SAXS data, we conducted a two-step evaluation of the predictors. First, we evaluated the overall quality of the predictors’ models using the *χ*^2^ metric in reciprocal space and filtered out models that diverged from the experimental data. The *χ*^2^ metric provides a rapid assessment of the overall structure. However, the *χ*^2^ metric can obscure subtle yet significant differences in the scattering. So, to detect whether those models with reasonable *χ*^2^ values actually captured the structural preferences of the protein ensemble, we compared the reciprocal and real space curves of the top predictors with corresponding experimental data, as a function of the scattering vector magnitude *q*.

To assess the overall quality of predictions, we calculated SAXS curves from the ensembles of atomic models provided by the predictors using FoXS and compared them to the experimental data using a standard workflow (Figure 5). The commonly-used *χ*^2^ metric was employed as a measure of accuracy; note that some of our predictors may have calibrated/trained their models with other SAXS data using this metric. *χ*^2^ evaluates the difference between model and experimental curves and is thus dominated by low *q* and interdomain distances [31]. Typically, a *χ*^2^ value between 1 and 2 is considered reasonable, with the caveat that *χ*^2^ is a function of the experimental error in the denominator. Models or model ensembles for the same protein based on the same experiment and analyzed using the same experimental error are comparable; however, *χ*^2^ values cannot be compared between different experiments. In this evaluation, 16 of 27 predictors had at least one prediction with a *χ*^2^ value between 3 and 10 (Figure 6, panels A, B, and C). The largest *χ*^2^ value exceeded 200 for the worst single PDB among all the conformations submitted, which is surprising given that the domain structures are known and only the structural conformations of 6 residues are unknown. As a reference, the ranking in the plot is sorted by the root mean square average of the *χ*^2^ values for T1200 and T1300.

**Figure 5:**
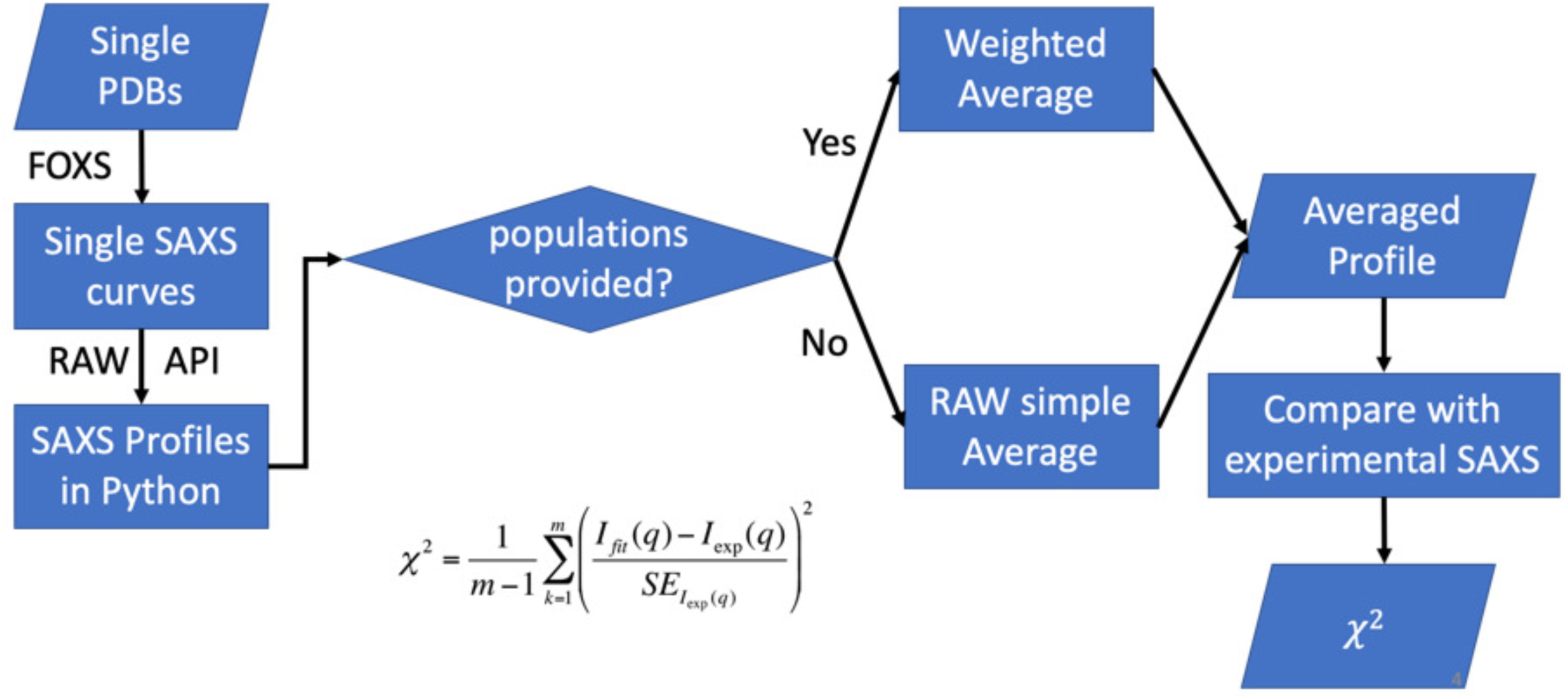
SAXS evaluation workflow. The single PDBs of each group were converted to single SAXS curves with FoXS for preprocessing (Left). The converted SAXS curves were averaged based on the population file provided with the PDBs (Right). The same evaluation workflow is applied on both targets. In the *χ*^2^ difference comparison between experimental data and reciprocal space curves calculated from predicted ensemble atomic models, *m* is the number of experimental data points, *I*_fit_(*q*) is the predicted intensity of each group’s ensembles models, *I*_exp_(*q*) is the experimental intensity, *SE_I_*_exp(_*_q_*_)_ is the standard error of each experimental data point, *q* is the scattering vector magnitude.

**Figure 6:**
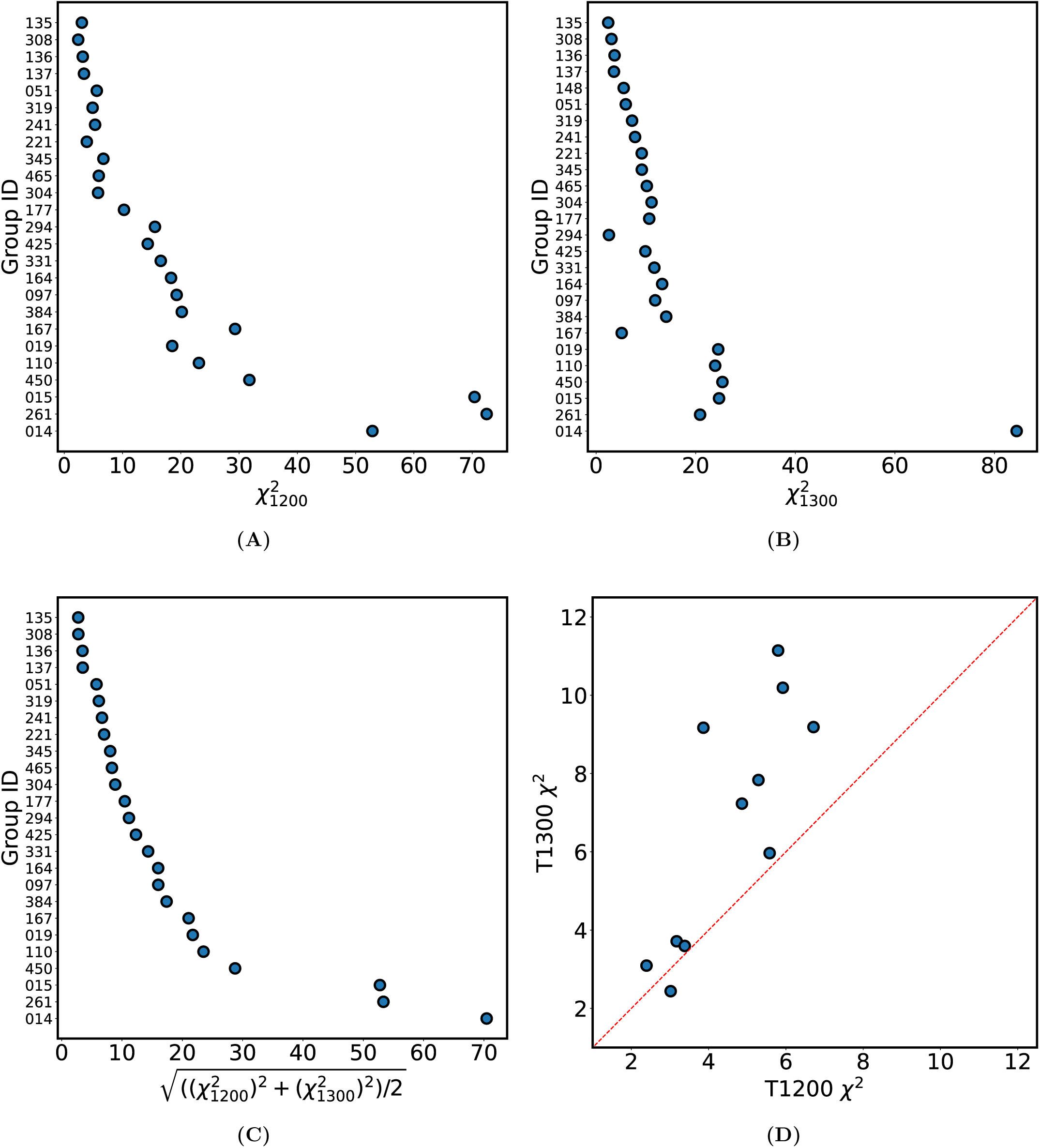
Comparison of ensemble-based predictions of SAXS scattering curves with ground truth using the *χ*^2^ metric (Section 2.2.2). (A) Average *χ*^2^ for T1200 (WT). (B) Average *χ*^2^ for T1300 (Gly6). (C) Root mean square average of the *χ*^2^ values for T1200 and T1300. (D) Comparison of predictor performance on WT vs. mutants for the predictions whose T1300 *χ*^2^ values were less than 12. Each point represents a predictor group. The *y*-axis shows the *χ*^2^ value for T1300, the Gly6 mutant. The *x*-axis shows the *χ*^2^ value for T1200, the WT protein. The dashed red line represents the diagonal, indicating the same performance on both targets. Panel (B) includes predictor 148, whose submission for target T1300, but not T1200, was valid (Section 3.1).

To see if any predictor model can reveal the difference between the two targets, T1200 and T1300, we plotted the *χ*^2^ for the two targets for groups that have *χ*^2^ lower than 10 for at least one target (Figure 6D). The data points for groups that perform similarly on both targets would fall near the diagonal. About half of the predictors performed better on the WT compared to the Gly6 mutant. This suggests that prediction methods may be more accurate in modeling endogenous linkers similar to those found in nature as opposed to artificial constructs such as Gly6 where there are no evolutionary relationships.

Because *χ*^2^ is a single metric for a three-dimensional comparison, we looked at the reciprocal and real space curves for predicted vs. experimental data for the two best-scoring predictors 135 (Lindorff-Larsenclvds) and 308 (momateam1) (Figure 7). More informative than visual inspection of the overlain curves, in terms of accuracy, is the clear bias in the residual difference between predicted and experimental data in reciprocal space (Δ*I*(*q*)*/σ*(*q*)) (Figure 7A,B). If the predictions were accurate, the residuals would fall with equal probability above and below the diagonal. For top scoring predictors 135 and 308, there are large residuals falling in the low-*q* region, which indicates an inaccurate prediction of the interdomain distance. Since the structures of the rigid domains were given, the mid- to high-*q* regions show relatively good predictions of the intradomain distance.

**Figure 7:**
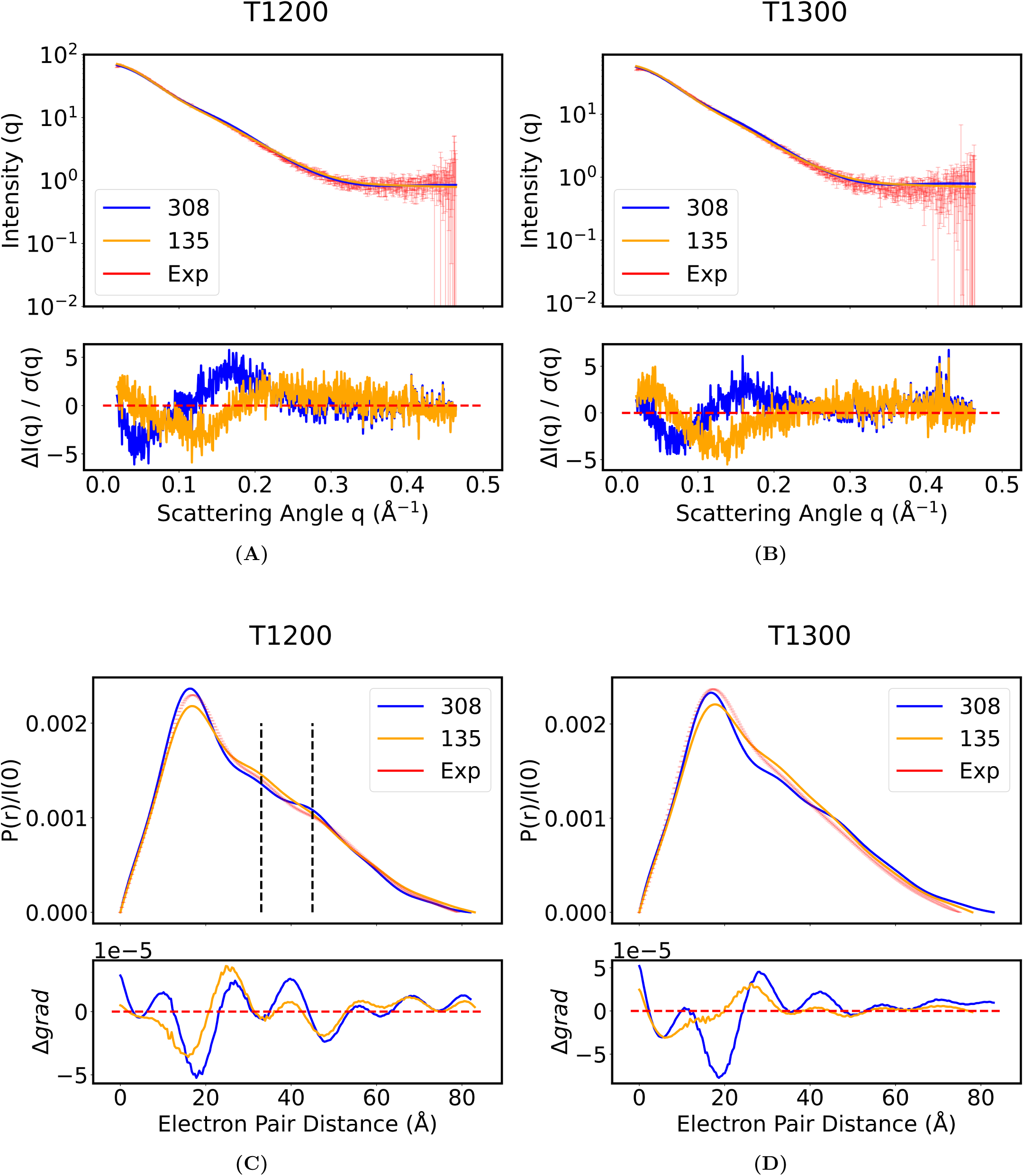
Comparison of two predicted ensembles (135 and 308) with experimental (“Exp”) SAXS data for T1200 and T1300 showing biased differences. SAXS curves were calculated from predictor’s ensemble of atomic models and overlaid with experimental data in reciprocal (A,B) and real (C,D) space for T1200 (A,C) and T1300 (B,D). Residual graphs between predictions and experimental are shown below each SAXS plot. Panels (A) and (B) show bars for experimental standard deviations, and panels (C) and (D) show bars (very small) for estimated error as described in Section 2.2.1.

We further plotted the change of the derivative in the *P* (*r*) curve to see if predictors captured the semi-stabilized domain-domain orientations indicated by the shoulders at 33 and 49 Å in the real space data (Figure 4B). These shoulders represent the preferred population, especially in the WT. For the WT predictions, we observed shoulders (Figure 7C), but they were at incorrect distances. The Gly6 predictions also had shoulders. However, shoulders were not present in the experimental data, indicating that the loss of preferred distance was not captured in these predictions (Figure 7D). No metrics that can quantitatively measure bias or preferred distances have been developed for SAXS analysis, hence our analysis here, based on SAXS data alone, can only qualitatively conclude that the predictors captured neither the preferred distance of the WT protein nor the loss of this preferred distance in the Gly6 mutant.

### 3.3 NMR

Predictions were compared to NMR RDC data both directly and through comparison with models as described in Materials and Methods. Briefly, a linear regression of back-calculated RDC values on experimental RDC values was computed for each predictor, and the slope and *y*-intercept of this regression were considered, along with the corresponding Pearson correlation. In addition, kernelized predicted ensembles were compared to uniform distributions in terms of free energy as a measure of anisotropicity. The ranges of these four quantities across predictors are shown in Table 2, along with ideal and worst possible values or range of values for that quantity, and SI Table S3 shows these four quantities for all predictors. The four quantities were normalized according to their worst possible values and then combined using Chebyshev distance, selecting for each predictor their worst performing score among the four (SI Table S4).

**Table 2:** Quantities used to compute Chebyshev distances. The first two columns are the minimum and maximum values of each quantity among all the predictions. The ideal value represents the quantity for perfect agreement with ground truth. Calculation of worst possible values is described in Section 2.2.4. Chebyshev distances were computed by scaling the prediction’s values for each quantity by the range from ideal to worst possible. The overall distance is the highest of these four values. SI Table S4 lists each of these values and the overall distance for each prediction.

| Quantity | Minimum<br>among<br>predictors | Maximum<br>among<br>predictors | Ideal value | Worst<br>possible<br>value |
| --- | --- | --- | --- | --- |
| Slope for RDC regression | -3.0 | 7.6 | 1.0 | -14.5 |
| Y-intercept for RDC regression | -3.7 | 1.7 | 0.0 | -8.8 |
| Pearson correlation for RDCs | -0.76 | 0.81 | 1.0 | -1.0 |
| Anisotropy (kcal/mol) | 0.40 | 2.45 | 0.09 to 0.42 | 2.50 |

Results of analysis for predictor 345 (multicom human), which had the smallest Chebyshev distance from ideal, are shown in Figure 8, with similar figures for every predictor in SI Section S7. Figure 8A shows the set of back-calculated RDCs plotted against experimental RDCs and with information shown regarding the linear regression. Figure 8B shows this predictor’s kernelized distribution in DoS form, along with the DoS for the solution CDIO corresponding to *α* = 0.98 (Figure 8C), which is the value of *α* for which the corresponding solution CDIO had the smallest free energy difference with predictor 345’s kernelized distribution. It is also clear from the DoS’s that this prediction is considerably less isotropic than the corresponding *α* = 0.98 solution CDIO, and it is anisotropicity that limits this predictor’s Chebyshev distance score, which is 0.30. Chebyshev distances range from 0.30 for predictor 345 (multicom human) to 0.98 for predictor 014 (Cool-psp). This latter ensemble’s Chebyshev distance is limited by its near-maximal anisotropicity; it consists of six structures, all similarly oriented.

**Figure 8:**
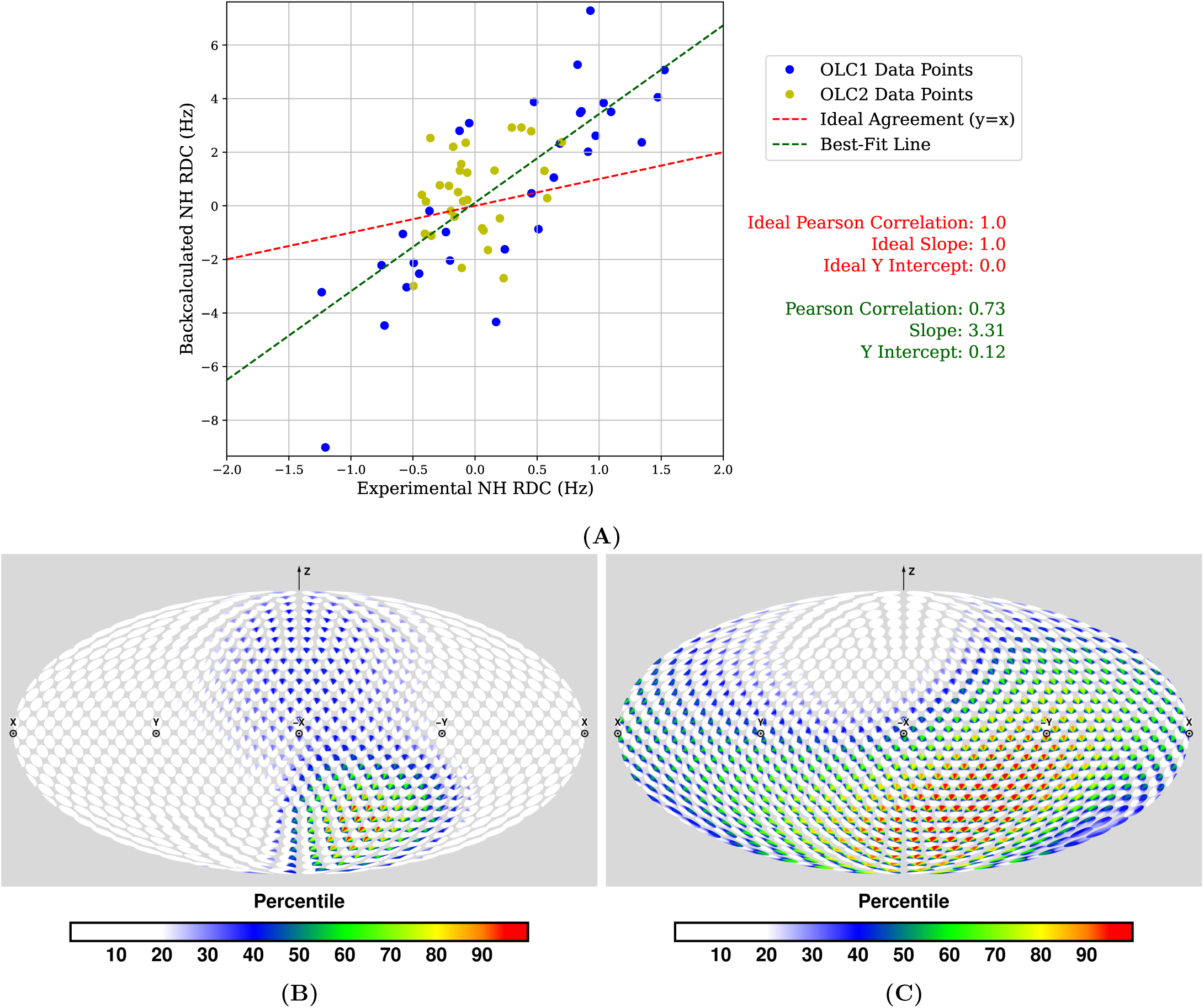
Representative data and assessment for predictor 345. (A) back-calculated RDCs plotted vs. experimental RDCs. *Green*: best-fit linear regression for back-calculated RDCs. *Red* : ideal regression line when the experimental and predicted RDCs are in perfect agreement (equal). (B) kernelized distribution of 345’s interdomain orientations. (C) mixture of 2 ground truth solution CDIOs (SI Figure S5, [47]) corresponding to *α* = 0.98, the value of *α* minimizing the free energy difference between the resulting *α*-mixture CDIO vs. the kernelized predictor distribution. (B,C) are disk-on-sphere (DoS) figures [47], in the Mollweide cartographic projection [54].

### 3.4 Joint evaluation of NMR and SAXS

An accurate prediction would be consistent with both SAXS and NMR RDC experimental data. These two types of data provide complementary information: SAXS data contain residue-nonspecific distance and orientational information of the entire protein, while NMR RDC provides residue-specific orientational information between the two domains. Both SAXS and NMR RDC data reflect distance and/or orientational information integrated, both temporally and spatially, over the conformational variation within an ensemble. NMR RDC data has relatively little noise, whereas SAXS data has relatively high noise.

SI Figure S5 shows that the CDIOs best fit to NMR RDC data demonstrate preference for some orientations over others. Similarly, the WT *P* (*r*) curve from SAXS in Figure 4B shows that there are populations in the WT ensemble demonstrating preferred interdomain distances; no such interdomain preferences were visually detectable in the *P* (*r*) curve for the Gly6 mutant. These observations confirm our previous descriptions of the WT linker as neither rigid nor completely flexible, and are consistent with the hypothesis that the conserved sequence of the WT linker encodes constrained flexibility. Thus, an ideal prediction consistent with data from both experiments would not describe an isotropic ensemble with uniform orientational and distance distributions. Instead, it would describe a somewhat anisotropic ensemble exhibiting these structural preferences in terms of both distance and orientation.

Figure 9 shows Chebyshev distances reflecting NMR-based analysis plotted against *χ*^2^ statistics reflecting SAXS analysis. We colored the dot for each prediction according to a broad categorization of the methods used to predict the diversity of linker conformations for each ensemble, as shown in Table 1. Where possible, we categorized these methods as being primarily based on molecular dynamics (blue), deep learning with default multi-sequence alignment (MSA) (green), and deep learning with other MSA methods (yellow-green). Light blue indicates a method that did not fall into these three categories; specifically, the d-i-tasser workflow used by predictor 019 (Zheng-Server), which uses replica exchange Monte Carlo (REMC) as the primary method for generating diverse conformations [68], IDPConformerGenerator, used by predictor 097 (jfk-thg-idpconfgen), which predicts ensembles of IDPs by sampling torsion angles found in the RCSB Protein Data Bank [57], and a workflow combining molecular dynamics with a statistical method used to predict conformations of IDPs, used by predictor 308 (momateam1) [23].

**Figure 9:**
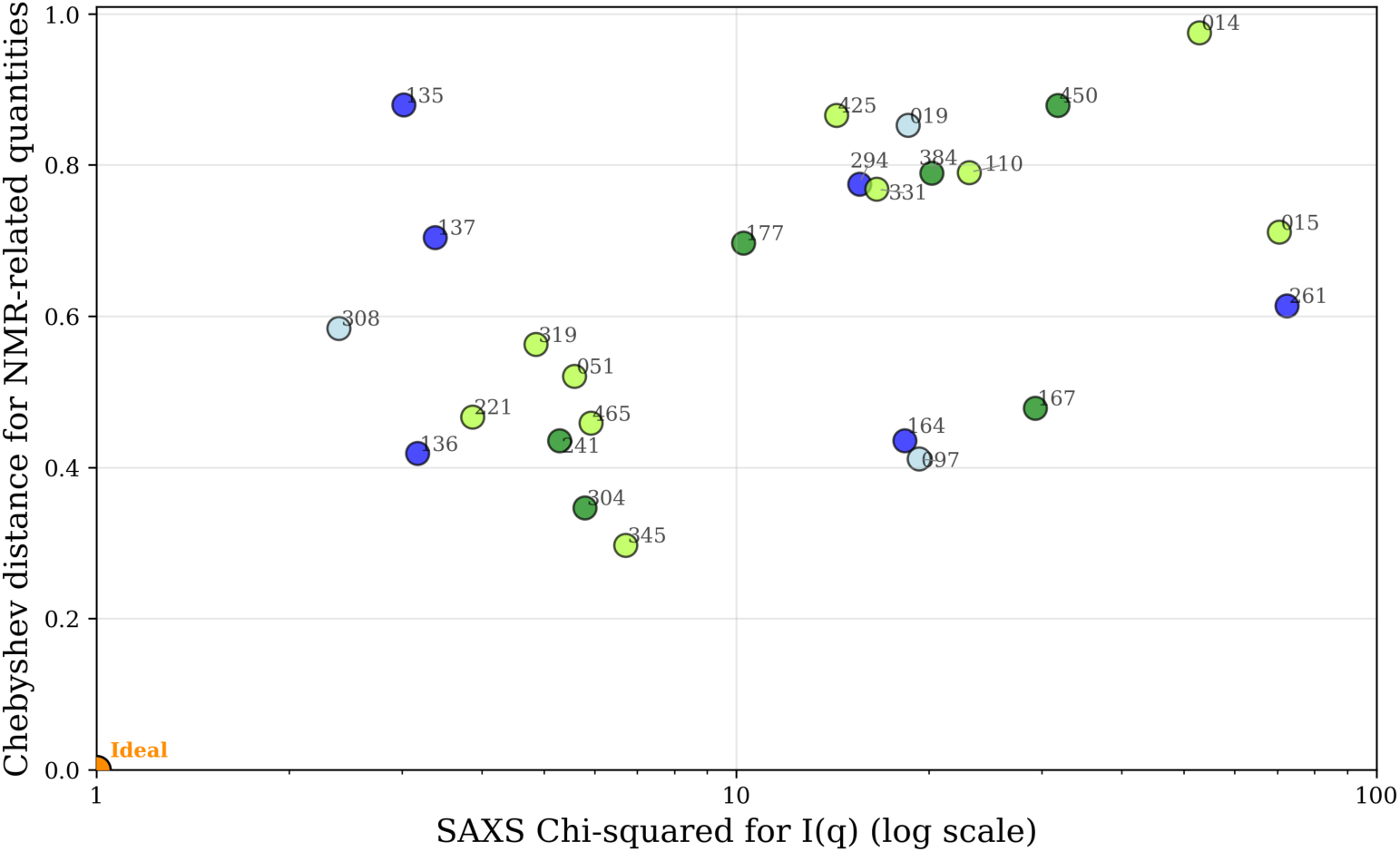
For each predictor, Chebyshev distances reflecting NMR-based analysis vs. *χ*^2^ statistics reflecting SAXS analysis. An ideal prediction would lie at the lower-left corner of the plot, with a Chebyshev distance of zero and a *χ*^2^ of one. *χ*^2^ statistics are plotted on a log scale to reflect that values greater than 10 are considered a poor fit to data. Dot color for each predictor reflects a broad categorization of methods used to predict the diversity of conformations for each ensemble (*blue*: molecular dynamics; *green*: deep learning; *yellow-green*: deep learning with custom MSA; *light blue*: other) and orange indicates the ideal values for a perfect prediction.

For each predictor, panels summarizing the various analyses described in this section are shown in SI Section S7.

## 4 Discussion

### 4.1 Comparison with other CASP challenges

In a typical CASP assessment, predictors have submitted one or more structures, each of which was compared individually to atomistic models. Even ensemble targets have taken this approach, for example, by comparing each predicted structure one-on-one with each member of a ground-truth ensemble. Our assessment is the first to assess each predicted population-weighted ensemble as an interconverting temporal or molecular thermodynamic ensemble. We assessed each predicted ensemble against both experimental SAXS and NMR RDC data as well as to ensemble models fit to experimental data.

In comparing predictions to NMR data, our assessment is somewhat similar to a previous CASP experiment that evaluated prediction models against NMR NOESY peak lists, backbone chemical shifts, and RDC data [30]. Unlike that experiment, which compared singleton atomistic structures to NMR observables, we treat the submitted set of population-weighted structures as a thermodynamic ensemble. Our assessment also differs by incorporating SAXS data as well as NMR.

Other past CASP challenges have used SAXS data [32, 44]. However, these were not blind assessments; rather, SAXS data were provided to predictors in advance, and comparisons were made largely on the basis of fit of predicted atomistic models to experimentally determined atomistic models from X-ray crystallography. These past CASP experiments demonstrated possible pitfalls of non-blind prediction: when initial structures with incorrect protein topology were adjusted to fit SAXS data, good fits could be achieved through distortion of the structure without correction of topological errors. By assessing data-blind predictions, we increased the likelihood that predictions that are good fits to data reflect more realistic models, as opposed to overfitting.

Our assessment’s use of back-calculated data allowed for comparisons of entire ensembles vs. ensemble-averaged NMR RDC and SAXS experimental data. Such whole-ensemble comparisons are necessary for our goal of assessing predictions of continuously varying interdomain pose. This flexible interdomain structure is not naturally described by fixed, finite numbers of discrete conformations but rather by continuous probability distributions. Therefore, discrete, pairwise comparisons between atomic structural models are not adequate to assess these predictions.

### 4.2 Evaluation of predictions

For this assessment, a good prediction must be close in terms of back-calculated SAXS curves, back-calculated RDCs, and anisotropicity. None of the predicted ensembles were close fits in all these measures. This lack of clear success in prediction resembles the first years of CASP.

Ensemble 345 (multicom human) is an example of a prediction that performed well by some measures but not others. The kernelized distribution based on this ensemble is shown in Figure 8 in comparison to the interpolation between Solutions 1 and 2 that is closest to it, in free energy terms (*α* = 0.98, which is nearly the same as Solution 1). Although the modes of the distributions in Figure 8B and Figure 8C are similar, comparing these two panels makes clear that ensemble 345 is too narrow; that is, it is less isotropic than either Solution 1 or 2 or any interpolation between them. Some qualitative aspects of the prediction can be observed in the best-fit regression for this ensemble’s back-calculated RDCs. In particular, the relatively good prediction of the Solution 1 mode is reflected in a relatively high Pearson correlation (0.73), and the prediction’s overly-narrow spread is reflected in an overly steep slope of 3.31. This steep slope derives from the overly large magnitudes of the back-calculated RDCs, which often results from a lack of averaging of RDCs corresponding to different orientations (Equation 1). Two other multicom predictions are even more dramatic in having accurate mode but overly-narrow distributions, which are similarly reflected in good Pearson correlations but very steep slopes (SI Figures S28 and S31).

While superficially, it appears from Figure 8B that ensemble 345 has a second mode similar to Solution 2 (just north of the equator of the DoS sphere), that is not the case. Close examination of the coloration within the relevant discs shows that this mode is rotated roughly 180^◦^ around the helical axis from the mode of Solution 2. This secondary mode of ensemble 345 and that of Solution 2 therefore have nearly orthogonal orientations (i.e., their corresponding quaternion representations are orthogonal as vectors in S^3^ ∈ R^4^), just as they would be if the disks representing them were on opposite sides of the DoS sphere.

Our results show that it remains challenging to predict, from sequence alone, ensembles of linker conformation (and, more generally, D-L-D protein structure) that are consistent with experimental data. This difficulty persisted even though predictors were given high resolution models of the two globular domains based on X-ray crystallography. The remaining challenge was thus to predict the conformational distribution of the 6 linker residues or, alternatively, the relative translation and orientation between the two relatively rigid domains. It is striking that a 6-residue structure prediction problem remains challenging for current methods. It is much easier to create ensembles consistent with data when RDC and/or SAXS data, at the level of ensembles or individual structures, are available prior to prediction for fitting and/or comparison.

Figure 9 summarizes our assessments, with a broad categorization of prediction methods indicated by different colors of dots. The small number of predictors in each category, combined with the diversity of methods within each category, preclude meaningful statistical analysis. If a clear pattern were qualitatively visible in the locations of differently-colored dots, we might be able to draw preliminary conclusions about strengths and weaknesses of different prediction methods, with respect to the relative orientation and translation of the two domains of the D-L-D system. No such clear patterns emerged in the data overall; however, we make some qualitative observations based on the top predictors with respect to the NMR RDC and SAXS axes of this figure.

Four of the nine non-deep-learning-based predictions were the best-performing predictions in terms of fitting SAXS data. These four predictions all used either energy functions tuned to SAXS data [60, 14] or methods known to produce predictions with good agreement to SAXS data [23]. In addition, these four predictions were also among the most isotropic of all ensembles considered, reflecting an enigmatic relationship between the anisotropicity of predictor ensembles and their closeness of fit to the SAXS data (SI Figure S6).

More broadly, neither molecular dynamics nor deep learning methods seemed to dominate the best predictions with respect to both SAXS and NMR data. The top four predictions with respect to SAXS data were based primarily or partly on molecular dynamics and were not among the best predictions with respect to NMR data. Conversely, the top three predictions with respect to NMR data were based on deep learning and were not among the best predictions with respect to SAXS data.

We have deliberately eschewed any single combined scoring for both SAXS *χ*^2^ statistics with NMR-based Chebyshev distances, because these two quantities reflect performance with respect to two completely different experimental modalities. Whenever entities are scored using two incommensurable values, it is possible to define a Pareto frontier of those entities with the property that every other entity is worse than it on at least one of the two scores. Figure 9 makes such a Pareto frontier clearly visible; it contains predictions 308, 136, 304, and 345.

Before we decided to use Chebyshev distance to combine three different quantities associated with best fits between back-calculated RDCs and experimental RDCs, we considered using more conventional measures of RDC closeness such as RDC RMSD [61, 22], *Q*-factor [58, 19], and *R*_dip_ [58, 18] (all of which are, in the context of fixed experimental data, approximately constant multiples of each other). If predictions had been closer fits to experimental data, these quantities may have been useful measures of that fit. However, no predictors’ back-calculated RDC sets were qualitatively close to the OLC RDCs. Those predictors whose back-calculated RDC sets were of similar magnitude to the OLC RDCs did not correlate well with the latter, and vice versa. As a result, variation in RDC RMSD, *Q*-factor, or *R*_dip_ was dominated by the magnitude of back-calculated RDCs rather than more meaningful closeness between the back-calculated and OLC RDC sets. Because our experimental RDCs are relatively small, the dominance of RDC magnitude in these measures had the practical effect of heavy bias toward more isotropic predicted ensembles, in which averaging resulted in back-calculated RDCs that were smaller in magnitude but generally uncorrelated with experimental RDCs. *Q*-factor and *R*_dip_ are reported in SI Table S2 alongside SI Figure S7, which illustrates how RDC RMSD and quantities proportional to it are not meaningful when back-calculated RDCs for different predictors have different ranges from each other and are also poor fits to experimental data.

### 4.3 Value of continuous distributions

Ensembles of discrete structures are commonly used to express various kinds of flexibility in protein structure, including that of flexible linkers. Such ensembles have the advantage that they are compatible with various computational tools designed to work with atomic models such as those deposited in the PDB. However, ensemble models fail to express the fundamentally continuous nature of protein flexibility or explicitly give the range of possible motion. In addition, they can express flexibility with many redundant degrees of freedom, specifying the positions of every atom when the relevant information may be expressible, for example, by a much smaller number of torsion angles. Some ensembles are designed to fit NMR and/or SAXS data with a minimal number of conformational models; such minimal ensembles may be easier for a human to understand [45], but sacrifice the ability to express flexibility or entropy in detail.

Continuous probability distributions offer descriptions of protein flexibility that can be at once precise, compact, and relatively simple. Such distributions are consistent with the assumption that flexibility is continuous in reality. In addition, choosing a parameterized family of distributions (such as Binghams over SO(3)) to represent potential solutions naturally regularizes the problem, potentially making proposed distributions more parsimonious by allowing for fewer degrees of freedom compared to discrete ensembles. Finally, as our study also demonstrates, there are practical analytical advantages to considering protein flexibility in terms of continuous distributions. For example, modes are well-defined on continuous distributions, and two continuous distributions can be compared in terms of their free energy difference via integration, given a known energy field (or the assumption that one of the distributions reflects this field). There remains the issue of comparing experimental SAXS data to probability distributions that include translational and rotational preferences, requiring further work in this area.

In the context of our assessment, the use of CDIOs facilitates the interpretation of ensembles’ degree of agreement with the NMR data. By comparing panels (C) and (E) in SI Figures S8 to S33, predictors may gain insight into changes to their predicted distribution of conformations that would yield closer fits to experimental data. Indeed, it is simpler to characterize the deficiencies of ill-fitting predicted ensembles in terms of distribution properties than in terms of the properties of individual structures. Although our assessment is primarily based on direct comparisons with experimental data, systematic, CDIO-based comparisons are also possible, and an example is described in SI Section S5.

The interpretation of best-fit solution CDIOs is complicated in this experiment, as it was in Qi et al. [47], by the presence of two CDIOs that are close fits to experimental data. The presence of two solutions with good fit is, at this point, an empirical observation; further work is needed to interpret this observation. It remains to be seen whether other D-L-D systems, including the T1300 (Gly6) construct, also yield two CDIO solutions that are close fits to RDC data. Further observations may shed light on the question of whether these multiple solutions relate to the “ghost” solutions described by Bertini et al. [8]. If so, it may be possible to rule out some solutions with RDCs measured using a fifth experimental alignment condition. A related avenue of future research would investigate other CDIO solutions that are not as close fits to the experimental data as the two considered in Qi et al. [47].

### 4.4 Value of predicting/understanding flexibility

In general, sequence conservation in flexible regions of proteins suggests a role in biological function, such as involvement in an interface. In the case of SpA-N, the linker sequences studied here are likely to be conserved because the constraints they place on flexibility help determine functionally significant interdomain pose distributions. However, more research is needed to clarify the relationship between sequence, flexibility, and function even in this particularly well-studied system.

The ability to predict interdomain pose distribution from sequence would be an important step toward understanding these natural systems, and the results of this study demonstrate that such predictive ability is still an open problem. In addition to contributing to our understanding of natural systems, such predictive ability would be beneficial in designing engineered systems. For example, therapeutics could be designed to have flexibility matching known pose distributions of target proteins or for optimized orientation of active sites to pass intermediates from one protein to another [64].

Our present study points the way toward improving our ability to predict pose distribution from sequence by demonstrating that experimental methods and robust biophysical measurements exist to empirically ground flexibility predictions. We have shown that both NMR RDCs and SAXS scattering curves can be back-calculated from ensemble predictions and used to assess the quality of flexible ensemble predictions. We expect hybrid collaborations combining these experimental methods with computational models to continue to play an important role in the advancement of predicting protein flexibility.

Further work is needed to solve the reverse problem of flexible structure determination based on NMR and SAXS data. In Qi et al. [47], we introduced a powerful method for determining orientational distributions from NMR RDC data. It would be better to have a method for determining not just an orientational distribution (over SO(3)) but a pose distribution (over SE(3)); such a pose distribution would represent a joint distribution of orientation and translation. Because RDCs carry only orientational information, other data sources would be needed to fit such a joint distribution. In principle, pseudocontact shifts caused by a paramagnetic lanthanide would contain both orientational and distance information; however, we see these shifts only in the ZLBT domain and none in the C domain. The idea of using SAXS information along with RDCs to fit a joint pose distribution is appealing, because the distribution of interatomic distances changes with both the relative orientation and translation of the two rigid domains; therefore, SAXS should in principle be sensitive to changes in pose distribution. Whether the information carried by SAXS data has sufficient resolution to allow fitting a joint pose distribution is an open question. Because SE(3) has six dimensions, as opposed to the three dimensions of SO(3), any reasonable selection of a family of distributions over SE(3) will have more parameter degrees of freedom than the nine of the Bingham distribution. Our fitting of Bingham distributions to RDC data relies on the novel algorithm described in Qi et al. [47], and fitting pose distributions to RDC and SAXS data is likely to similarly require mathematical and algorithmic advances.

## 5 Conclusion

CASP16 targets T1200 and T1300 asked participants to predict the conformational distribution of two flexible linkers, each part of a D-L-D construct designed to explore the flexibility of SpA-N. These targets represent the first blind challenge to predict population-weighted ensembles of structures to be compared both to experimental data as well as to ensemble models. The introduction of such targets to CASP recognizes the importance of predicting distributions of conformations realized in solution.

None of the predictions were close fits either for the experimental data or for our optimally-fit orientational distributions. These results demonstrate that predicting the distribution of poses of rigid domains connected by flexible linkers remains an unsolved problem. We conclude that NMR RDC and SAXS data work well together as complementary experimental modalities reflecting different aspects of the pose distribution of D-L-D systems. Further work is needed to fit continuous, low-dimensional pose distributions to SAXS and NMR RDC information. Given the success of AlphaFold and similar models in predicting static structures, the prediction of protein conformational dynamics driving biological function is an important frontier in structural biology. We hope that future CASPs continue to build on the new directions our experiment has established in addressing this challenge.

## Data availability

Both predictions and SAXS data will be uploaded to the CASP website before publication. NMR RDC data are available at BMRB (accession number 53002) [62]. All data processing scripts will be uploaded to https://github.com/donaldlab/CASP16 upon acceptance.

## Acknowledgments

We thank Henry Childs for help in writing the abstract of this report, and all members of the Donald Lab for comments and discussion. We also thank Greg Hura for his work with Susan E. Tsutakawa in initiating CASP- SAXS.

We are grateful to John Moult, Krzysztof Fidelis, and Andriy Kryshtafovych for their work in organizing CASP. We also thank all the predictors who made this work possible by participating in these CASP targets, and whose submitted abstracts we used to categorize prediction methods.

This work was supported by the National Institutes of Health (R35 GM-144042 to Bruce R. Donald and NIGMS R35 GM141818 to Gaetano T. Montelione). This work was also supported by the U.S. Department of Energy, Offices of Biological and Environmental Research; Basic Energy Sciences and Advanced Scientific Computing Research through the Biopreparednesss Research Virtual Environment (BRaVE) under contract number DE- AC02-O5CH11231 to Taskforce 5 (DOE-BRAVET5) and to Integrated Diffraction Analysis Technologies (IDAT). In addition, this work was supported by NCI P01 CA092584 (to Susan E. Tsutakawa and Michal Hammel) and 1R01GM137021 (To Susan E. Tsutakawa)

## Notes

^1^They are not independent; for example, as Pearson correlation gets close to zero, slope becomes less meaningful.

