## Supplemental Information for "Predicting pose distribution of protein domains connected by flexible linkers is an unsolved problem"

This Supplementary Information (SI) briefly elaborates specific points in the main text, along with additional figures. It also contains an assessment figure for every predictor, with panels showing the results of our assessment for that predictor with respect to comparisons based directly on SAXS and NMR RDC as well as comparisons with CDIO models optimally fit to NMR RDC data. Finally, for this CASP experiment, to completely and rigorously report the precise experimental conditions in the form of the specifications given to the predictors, as a reference, it also contains two documents that were provided to all participants in this CASP target: a "Specifications" document detailing the requirements of the challenge, and an "Algorithm" document explaining our procedure for determining the relative alignment of ZLBT and C domains of a given model of either the T1200 or T1300 system.

#### S1 OLC calculations

For clarity in the main text, we use the term "OLC RDCs" to refer to the sets of RDC-based numbers used in comparing back-calculated and experimental values. Each of these values is actually a linear combination of experimental RDCs. Like RDCs, they have units of hertz, and they represent the RDCs corresponding to putative mutually orthogonal alignment conditions.

†These authors contributed equally to this work

The calculation of these OLC RDCs is mostly described in Qi et al. [3], which cites Ruan and Tolman [4] for the underlying method. However, the precise method used in Qi et al. [3] is a variation of that reported by Ruan and Tolman [4]: Rather than perform SVD on the RDC data sets themselves, SVD is instead performed on vectorized Saupe tensors fit to the ZLBT domain RDCs, with similar results.

### S2 RDC background

The experimental residual dipolar coupling (RDC) between nuclei $i$ and $j$ is given by:

$$d_{ij} = \frac{\mu_0 \gamma_i \gamma_j \hbar}{4\pi^2 r_{ij}^3} \left\langle \frac{3\cos^2(\psi_{ij}) - 1}{2} \right\rangle, \tag{S1}$$

where $\mu_0$ is the magnetic permeability of free space, $\gamma_i, \gamma_j$ are the gyromagnetic ratios of the nuclei, $\hbar$ is the reduced Planck's constant, $r_{ij}$ is the length of the covalent bond between the nuclei (assumed to be known from high-resolution crystal structures), and $\psi_{ij} \in \mathrm{S}^1$ is the angle between the bond vector and magnetic field vector.

Defining the dipolar coupling constant as:

$$k = \frac{\mu_0 \gamma_i \gamma_j \hbar}{4\pi^2 r_{ij}^3}, \tag{S2}$$

the RDC equation is equivalent to:

$$d = \frac{k}{2} \mathbf{v}^\intercal \mathbf{S} \mathbf{v}, \tag{S3}$$

where $k$ is the dipolar coupling constant, $\mathbf{v}$ is the unit bond vector, and $\mathbf{S}$ is the Saupe tensor that describes the average molecular alignment [1].

If $\mathbf{S}$ is the Saupe tensor describing a rigid domain with partial alignment (the ZLBT domain in the present case), and if $p(R)$ is the probability distribution function describing the orientation $R$ of a second rigid domain relative to the first (the C domain in our case), then for RDCs in the second domain we can write:

$$d = \frac{k}{2} \int_{\mathrm{SO}(3)} \mathbf{v}^\intercal R^\intercal \mathbf{S} R \mathbf{v}\, p(R)\, dR. \tag{S4}$$

(We abuse notation here by using $R$ to refer both to an element of SO(3) and to the matrix representation of that element.)

### S3 Predictor ensemble kernelization

As explained in Section 2.2.3, we cannot use integrals to directly compare discrete ensembles to continuous distributions such as CDIOs. Therefore, we use kernel density estimation [2] to estimate continuous probability density functions of interdomain orientation from the discrete probability functions directly implied by predicted population-weighted ensembles. By doing so, we assume that each interdomain orientation in a discrete ensemble is a sample drawn from a distribution that is a sum of kernel functions, one centered at each of the discrete orientations.

Formally, we are given a finite set of $n$ protein structures with interdomain orientations $\theta_1$, ..., $\theta_n \in \mathrm{SO}(3)$ and population weights $w_1$, ..., $w_n$. Kernel density estimation gives us a continuous probability density function $f : \mathrm{SO}(3) \to \mathbb{R}$ such that $f$ is the sum of $n$ kernel functions $g_i$, each with mode $\theta_i$ and weight $w_i$.

It is natural to choose a Bingham distribution for each kernel function $g_i$, for the same reason that we use Bingham distributions for our CDIOs, which is that the Bingham distribution is the natural analog of the Gaussian distribution over SO(3). Therefore, our kernel density estimate for a given discrete ensemble will be

$$f(\theta) = \sum_{i=1}^{n} g_i(\theta) \propto \sum_{i=1}^{n} w_i \exp(\theta^{\intercal} A_i \theta) \tag{S5}$$

for any $\theta \in \mathrm{SO}(3)$. (We abuse notation here by using $\theta$ to represent both an element of SO(3) and its quaternion representation. In addition, we omit the details of the hypergeometric normalization constants associated with Bingham distributions. In the work described here, we normalize the distributions by dividing by the numerical integral of the unnormalized values.)

It remains to choose the parameter matrix $A_i$ for each kernel function. The interdomain orientation $\theta$ of a given structure in a predicted ensemble implies no information about the relative likelihood of any two different orientations $\zeta_1$ and $\zeta_2$, if the angular distance between $\theta$ and $\zeta_1$ is the same as the angular distance between $\theta$ and $\zeta_2$. (The angular distance between two orientations is the angle of the axis-angle rotation taking one to the other; if the two orientations are represented as rotation matrices $P$ and $Q$, this angle is

$$\cos^{-1}\left(\frac{(\mathrm{Tr}(PQ^{\intercal}) - 1)}{2}\right), \tag{S6}$$

where Tr is the trace of a matrix.) Therefore, it is natural to choose a kernel that has as its mode the orientation of the given predicted structure and exhibits circular symmetry around this mode. We also want this kernel to be small in variance (much smaller, for example, than the ground-truth CDIOs). This small variance corresponds to the interpretation of the interdomain orientation of a single predicted structure as suggesting that very similar interdomain orientations are also somewhat likely, but saying little about less similar interdomain orientations. The parameter matrix of a Bingham distribution fitting this description is

$$A = \lambda\, \mathbf{v} \otimes \mathbf{v}, \tag{S7}$$

where $\mathbf{v}$ is the mode of this distribution, and $\lambda$ is a parameter controlling the spread of this distribution around its mode. Note that the mode corresponds to the largest-eigenvalue eigenvector of $A$, and $\mathbf{v} \otimes \mathbf{v} = \mathbf{v}\mathbf{v}^{\intercal}$ is the tensor or outer product of this mode vector with itself.

To choose a $\lambda$ to use for the kernelization of predictor ensembles, we varied $\lambda$ according to $-(2^i), \forall i \in \{0, 1, \cdots, 10\}$; for each $\lambda$, we took 8192 random samples from each of our solution CDIOs, similar to the maximum size (10,000) of a predictor ensemble. We kernelized these samples using $\lambda$, and computed the free energy difference between the resulting distribution and the corresponding solution CDIO. We chose the value of $\lambda$ that resulted in the smallest such free energy difference; this $\lambda$ value was $-32$. In other words, we chose $\lambda$ to be approximately that which would allow a maximally-sized ensemble to best reproduce one of our CDIO solutions.

#### S4 Free energy integral comparisons

All free energy differences between two continuous probability distributions of interdomain orientation were calculated according to

$$\Delta F = RT \int_{SO(3)} p_2(\theta) \ln \frac{p_2(\theta)}{p_1(\theta)}\, d\theta \tag{S8}$$

which is simplified from the derivation in Qi et al. [3], where $R$ is the gas constant, $T$ is temperature (assumed here to be 25 °C), $p_2$ is a probability distribution being compared to $p_1$, and $\theta$ is a given orientation. The integral was approximated by a Riemann sum over a grid of 36,864 points on SO(3). These points, generated by the algorithm described in Yershova et al. [6], are at the centers of approximately equal-sized volume elements.

#### S5 Using CDIOs to characterize predictions

**Algorithm 1** This procedure is used to plot each prediction in Figure S1.

1. For each kernelized predicted distribution, we select from among our CDIO solutions the one with which the prediction has the smallest free energy difference.
2. Each prediction is rotated such that its mode is equal to the mode of the selected CDIO solution for that prediction.
3. The free energy difference between this rotated prediction and the selected solution is plotted on the $x$-axis.
4. The angular distance (as given by Equation (S6)) between the modes of the (un-rotated) prediction and the selected CDIO solution is plotted on the $y$-axis.

CDIOs may be useful in interpreting how given ensemble predictions could be improved; Figure S1 shows an analysis illustrating this utility. A given predictor's location in Figure S1 relates to the CDIO solution with the smallest free energy difference with the predicted ensemble, as in panel (E) of the figures in Section S7. Location on the $y$-axis reflects the accuracy of the mode of the predictor's ensemble, with respect to this closest CDIO solution. Location on the $x$-axis reflects a comparison between the prediction and the closest CDIO solution after rotating the former so that its mode matches the CDIO solution. The $x$-axis thus reflects the accuracy of the shape of the prediction and the orientations of its principle axes of variance, independently of the location of its mode. (In other words, if a predictor predicted a distribution that was correct except for having the wrong mode, they would be plotted somewhere along the $y$-axis of this figure.) From this figure, one can see that a given predicted ensemble would be more similar to a best-fit solution CDIO if, for example, its mode were rotated or its shape were changed. Such interpretation is complicated, of course, by the existence of more than one solution CDIO that is a close fit to the experimental data. Moreover, the information shown in this figure is not part of our assessment, which is summarized by main text Figure 9.

#### S6 Uncertainty analysis

Qi et al. [3] report 0.2 Hz experimental error, as an overall estimate for all RDCs. This error is shown in Figure S2, which is otherwise identical to main text Figure 2.

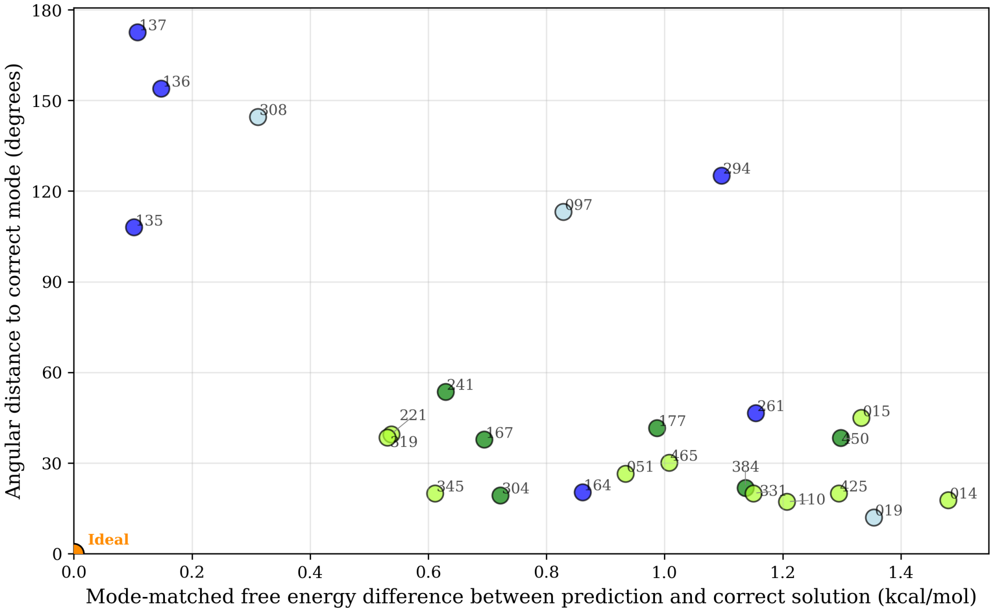


**Figure S1: For each predictor, mode-matched free energy difference and mode angular distance, both in comparison to the closest correct solution.** Algorithm 1 is used to compute the accuracy of each predicted ensemble's mode, based on the angular distance between the mode and that of the best fit solution (y-axis). Algorithm 1 is also used to compare the shape of the predicted distribution and the orientations of its principle axes of variance to those of the best-fit solution by superimposing the modes of each (x-axis). As in main text Figure 9, dot color for each predictor reflects a broad categorization of methods used to predict the diversity of conformations for each ensemble (*blue*: molecular dynamics; *green*: deep learning; *yellow-green*: deep learning with custom MSA; *light blue*: other).

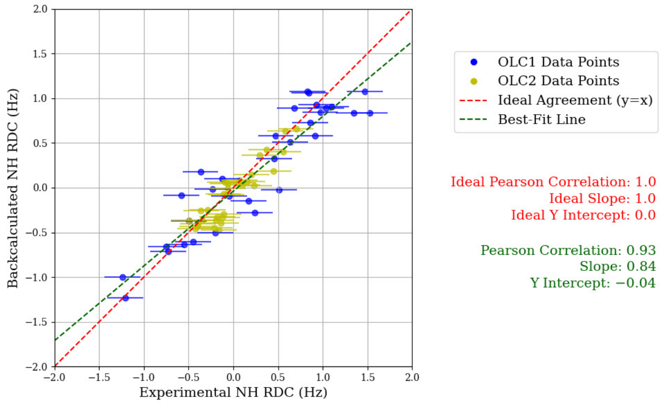


**(A)** Solution 1

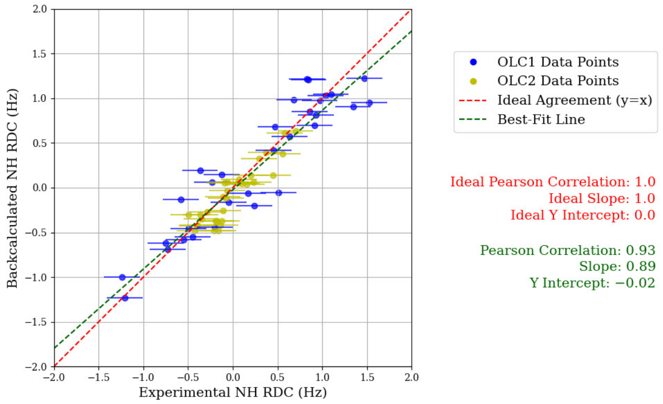


**(B)** Solution 2

**Figure S2: Comparing CDIO solutions to NMR RDC data, with experimental errors shown.** Linear regressions of OLC RDCs back-calculated from (A) Solution 1 and (B) Solution 2, as in main text Figure 2, with ±0.2 Hz estimated experimental error shown as horizontal bars.

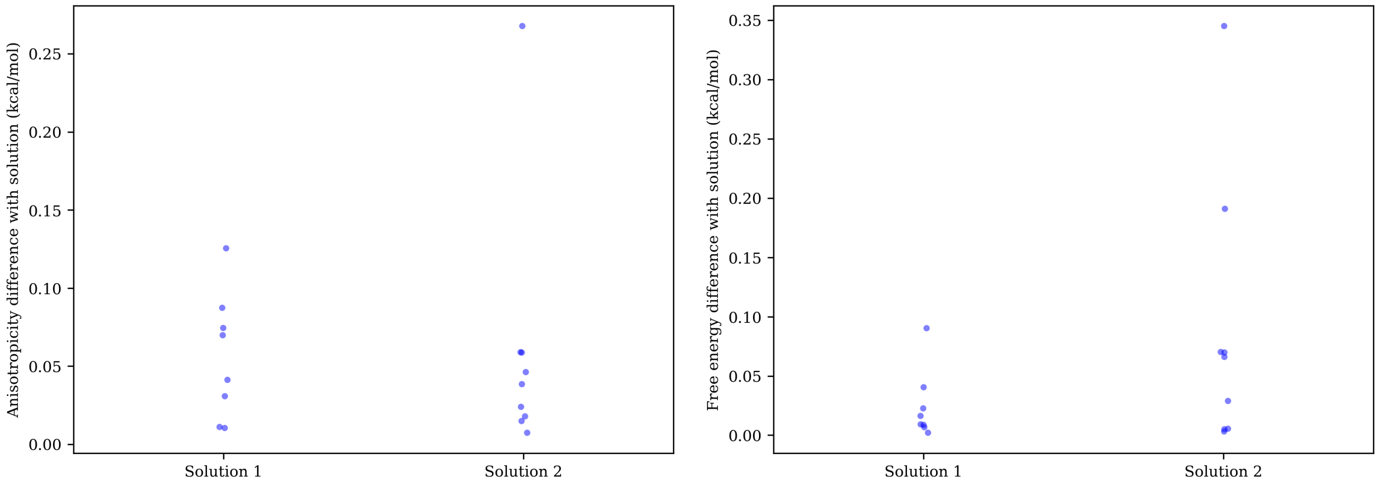


**(A)** **(B)**

**Figure S3: Best-fit CDIO solutions are robust to experimental uncertainty.** We generated 10 sets of RDCs, perturbed from the experimental RDCs according to the uncertainty reported in Qi et al. [3], and found best-fit CDIOs for each. The 8 best-fit CDIOs similar to Solution 1 and the 9 best-fit CDIOs similar to Solution 2 are plotted in terms of their differences with the solution CDIOs. (A) Anisotropicity difference between perturbation-based CDIOs and solution CDIOs, where anisotropicity is defined as in main text Section 2.2.3. (B) Free energy differences between the perturbation-based CDIOs and solution CDIOs, computed with the solution CDIOs treated as correct, as described in Sec. S4.

To investigate the robustness of our best-fit CDIO solutions with respect to experimental uncertainty, we fit CDIO solutions to perturbed sets of RDCs, similar to the approach described in Tripathy et al. [5]. To perturb the experimental RDCs, we applied Gaussian noise with a standard deviation of 0.2 Hz to each RDC value, according to the reported experimental error in Qi et al. [3]. We generated 10 sets of RDCs perturbed in this manner, and applied to each set the method from Qi et al. [3] to search for best-fit Bingham distributions, as with the measured RDCs [3]. In 7 out of these 10 sets, the top two qualitatively distinct CDIOs were similar to Solution 1 and Solution 2 as reported in main text Section 2.1. In 2 of the 10 sets, only one close-fitting distinct CDIO was found, and this CDIO was similar to Solution 2. In 1 of the 10 sets, the best-fit CDIO was similar to Solution 1, and a second close-fitting CDIO was found that resembled neither Solution 1 nor Solution 2. In summary, 8 CDIOs similar to Solution 1 were found and 9 similar to Solution 2.

Figure S3 quantifies the similarity of the perturbation-based CDIOs to the Solution 1 and Solution 2 CDIOs using difference in anisotropicity (main text Section 2.2.3) and in free energy (Section S4). Anisotropicity is used as part of our assessment of predictions, and free energy differences were used in Qi et al. [3] to investigate biological function, and are used to select nearest solutions for visualization in panel (E) of Figures S8 to S33 in Section S7. Differences in anisotropicity shown in Figure S3A are small compared to the anisotropicity differences between predictor ensembles and the CDIO solutions fit to experimental data, as seen in Figure S6. (A few predicted ensembles were very close to our anisotropicity range for CDIO solutions fit to data; however, Table S4 shows that in these cases, their Chebyshev distances were determined by poor agreement between back-calculated and experimental RDCs rather than by anisotropicity. Therefore, Chebyshev distances for these predictions are not affected by small changes to the range of anisotropicity considered as correct.) Free energy differences shown in Figure S3B are mostly small compared to the differences in free energy between predicted ensembles and CDIO solutions fit to data; these latter differences between predicted ensembles and best-fit solutions range from 0.17 to 1.57 kcal/mol. These results demonstrate that our best-fit CDIO solutions are robust to experimental uncertainty.

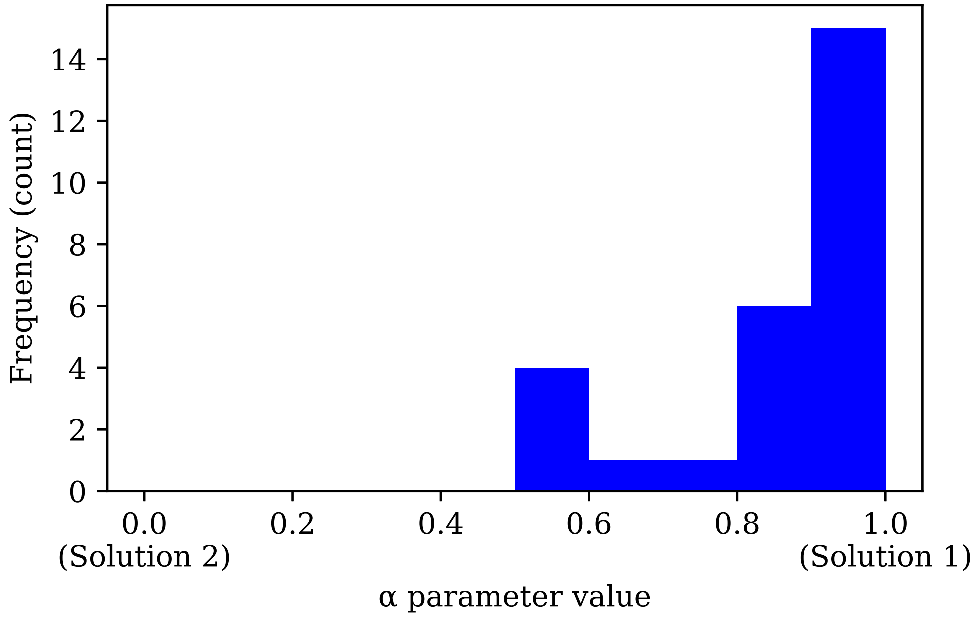


**Figure S4: Distribution of $\alpha$ values for predicted ensembles.** For each predictor, the value of $\alpha$ was chosen that was associated with the smallest difference in free energy between the corresponding mixture CDIO (as the distribution for comparison) and the kernelized predictor CDIO. The frequencies of these alpha values are shown in bins of 0.1 width.

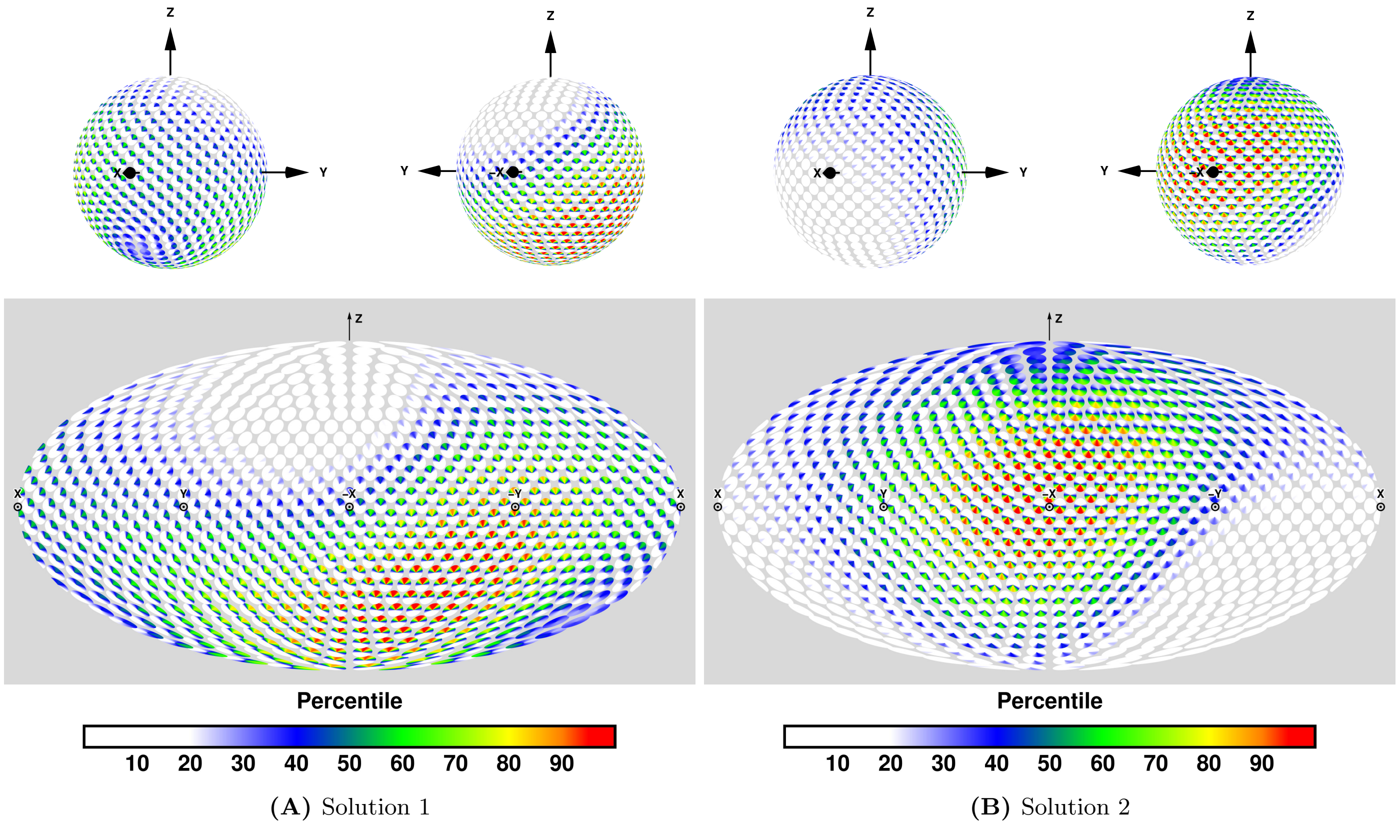


**Figure S5: Disk-on-Sphere representations of Bingham CDIOs optimally fit to NMR RDC data, for (A) Solution 1 and (B) Solution 2.** For each solution, a perspective projection is shown (top), front and back, and the same information in Mollweide projection (bottom). Colors indicate percentiles; a given orientation $R$ is assigned a color such that the corresponding percentage of total probability mass is found at orientations less likely than $R$. This color scheme is the same as that used in Qi et al. [3].

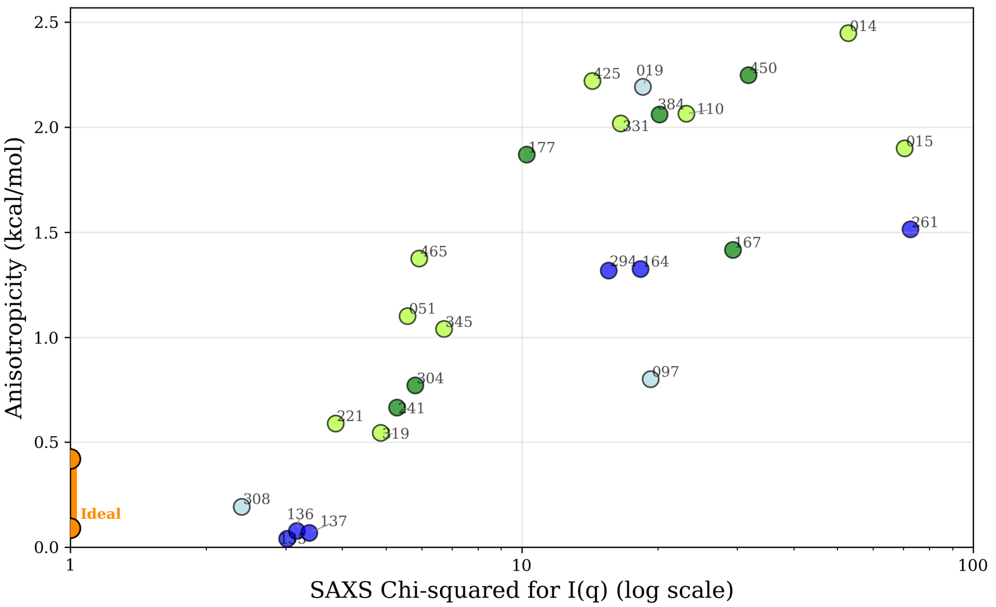


**Figure S6: For each predictor, anisotropicity vs. $\chi^2$ statistics reflecting SAXS analysis.** Anisotropicity was calculated based on NMR RDC data as explained in main text Section 2.2.3. Ideal predictions would lie along the orange line segment in the lower left, with anisotropicity between 0.09 and 0.42 kcal/mol, and with a $\chi^2$ of one. As in main text Figure 9, dot color for each predictor reflects a broad categorization of methods used to predict the diversity of conformations for each ensemble (*blue*: molecular dynamics; *green*: deep learning; *yellow-green*: deep learning with custom MSA; *light blue*: other).

| Pred. Name | Pred. ID | # of Models (T1200) | # of Models (T1300) | Category | Ensemble Diversity Methods | Model Filtering | Weighting | Rigid Domain RMSD (Å) (T1200) | Rigid Domain RMSD (Å) (T1300) |
|---|---|---|---|---|---|---|---|---|---|
| JFK-THG-AMBER | 002 | 1000 | 1000 | Other | IDPConformerGenerator / Amber | Clustering | | 1.19 | 1.64 |
| JFK-THG-AMBERstable | 003 | 1000 | 1000 | Other | IDPConformerGenerator / Amber | Clustering | | 1.21 | 1.66 |
| JFK-THG-CHARMM | 004 | 1000 | 1000 | Other | IDPConformerGenerator / Charmm | Clustering | | 1.23 | 1.03 |
| JFK-THG-CHARMMstable | 005 | 1000 | 1000 | Other | IDPConformerGenerator / Charmm | Clustering | | 1.30 | 1.05 |
| Cool-PSP | 014 | 6 | 6 | DL (MSA) | AlphaFold 2-based model with custom MSA | Clustering | Built-in scoring | 0.60 | 0.64 |
| PEZYFoldings | 015 | 4 | 8 | DL (MSA) | AlphaFold 2 with PLM-based MSA | | pLDDT scores | 0.48 | 0.46 |
| Zheng-Server | 019 | 3 | 5 | Other | D-I-TASSER (REMC-based) | Clustering (SPICKER) | Cluster population | 0.57 | 0.58 |
| MULTICOM | 051 | 1000 | 1000 | DL (MSA) | AlphaFold 2 & 3 with various MSA | | Various scoring | 0.46 | 0.45 |
| Vendruscolo | 084 | 33 | 38 | | Not disclosed | Not disclosed | Not disclosed | 1.19 | 1.08 |
| orangeballs | 088 | 500 | 500 | DL | AlphaFlow | | | 0.96 | 1.08 |
| JFK-THG-IDPCONFGEN | 097 | 1000 | 1000 | Other | IDPConformerGenerator | Weighting normalization | | 0.41 | 0.41 |
| zurite_lab | 100 | 100 | 100 | | Not disclosed | Not disclosed | Not disclosed | 1.60 | 1.45 |
| MIEnsembles-Server<br>Zheng-Multimer<br>Zheng | 110<br>147<br>462 | 27 | 32 | DL (MSA) | DMFold | Clustering (SPICKER) | pLDDT scores | 0.57 | 0.59 |
| Lindorff-LarsenCLVDS | 135 | 1000 | 1000 | MD | CALVADOS | | | 0.18 | 0.18 |
| Lindorff-LarsenM3PPS | 136 | 1000 | 1000 | MD | Martini 3 | | | 0.16 | 0.16 |
| Lindorff-LarsenM3PWS | 137 | 1000 | 1000 | MD | Martini 3 | | | 0.16 | 0.15 |
| DeepFold-refine | 139 | 2000 | 2000 | MD | Amber | | | 1.13 | 1.75 |
| GuijunLab-Complex† | 148 | 162 | 173 | DL (MSA) | AlphaFold-Multimer and DeepAssembly with various MSA | Hierarchical networks | | 0.59 | 0.58 |
| McGuffin | 164 | 626 | 366 | MD | ReFOLD4 | | ModFOLD9 | 0.69 | 0.67 |
| OpenComplex | 167 | 10000 | 10000 | DL | OpenComplex-2 | Clustering | | 0.34 | 0.34 |
| aicb | 177 | 71 | 89 | DL | OpenFold | Clustering (MaxCluster) | Amber | 0.75 | 0.69 |
| CSSB_FAKER<br>CSSB_experimental<br>CSSB_Human | 221<br>286<br>419 | 1000 | 1200 | DL (MSA) | Faker and UltimateMSA | | | 0.58 | 0.57 |
| elofsson | 241 | 500 | 500 | DL | AlphaFold 3 | Clustering | | 0.58 | 0.57 |
| UNRES | 261 | 100 | 100 | MD | UNRES (MREMD-based) | Clustering | WHAM | 0.05 | 0.05 |
| GuijunLab-Human† | 264 | 30 | 60 | DL | DeepAssembly | GraphCPLMQA | | 0.59 | 0.58 |
| KiharaLab | 294 | 1000 | 1000 | MD | Desmond | | | 0.00 | 0.00 |
| AF3-server | 304 | 100 | 100 | DL | AlphaFold 3 | Clustering | | 0.58 | 0.57 |
| MOMAteam1 | 308 | 918 | 753 | Other | Statistical methods, Amber | Rigid domain RMSD | Cluster population | 0.70 | 0.68 |
| MULTICOM_LLM | 319 | 1000 | 1000 | DL (MSA) | AlphaFold 2 & 3 with various MSA | | GATE,<br>pLDDT,<br>AlphaFold 3 ranking | 0.46 | 0.45 |
| MULTICOM_AI | 331 | 1000 | 1000 | DL (MSA) | AlphaFold 2 & 3 with various MSA | | pLDDT scores | 0.47 | 0.45 |
| MULTICOM_human | 345 | 2000 | 2000 | DL (MSA) | AlphaFold 2 & 3 with various MSA | | pLDDT scores with human intervention | 0.47 | 0.45 |
| pert-plddt | 384 | 56 | 82 | DL | OpenFold | Clustering (MaxCluster) | Amber | 0.75 | 0.61 |
| MULTICOM_GATE | 425 | 1000 | 1000 | DL (MSA) | AlphaFold 2 & 3 with various MSA | | GATE | 0.47 | 0.45 |
| OpenComplex_Server | 450 | 10000 | 10000 | DL | OpenComplex-2 | Confidence scores | | 0.28 | 0.28 |
| Wallner | 465 | 17 | 21 | DL (MSA) | AFsample2 | Rosetta Clustering | Cluster population | 0.22 | 0.22 |

**Table S1: Information about each predicted ensemble, including the number of atomic structural models in each predicted ensemble, key prediction methods, and rigid domain RMSD.** The "Rigid domain RMSD" column gives the average across the two rigid domains of atomic backbone RMSD for certain residues (Section 2). This quantity is used to filter out predicted ensembles with rigid domain distortions too large for reliable analysis. It is not a measure of prediction accuracy, because rigid domain reference structures were provided to predictors; therefore, this RMSD was 0 for predictors who used the reference structures in their predictions. Rows are shaded in gray for ensembles which were above the cutoff of 0.75 Å and therefore not further analyzed. With respect to prediction methods, this table omits information on refinement methods applied to each predicted structure in an ensemble; instead, it focuses on three key questions relating to the overall shape of an ensemble: What methods were most important in generating a pool of decoys ("Ensemble diversity methods")? What methods, if any, were used to filter this pool of decoys ("Model filtering")? What methods, if any, were used to determine the population weight associated with each structure in the final ensemble ("Weighting")? For qualitative analysis purposes, we categorized predictions according to methods used to generate a pool of decoys: "DL" for deep learning with default MSA, "DL (MSA)" for deep learning with custom MSA, "MD" for molecular dynamics, and "Other" for other methods such as Monte Carlo. †Ensemble 148 (target T1200) and ensemble 264 (both targets) contained mixed-sequence predictions and were not assessed; see Section 3.1.

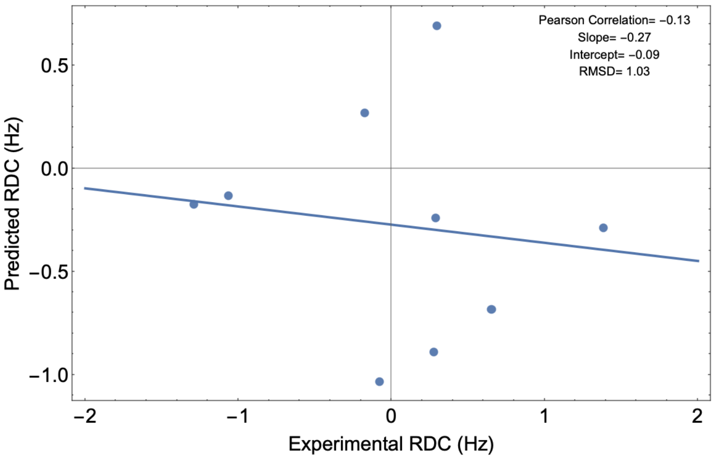


**(A)**

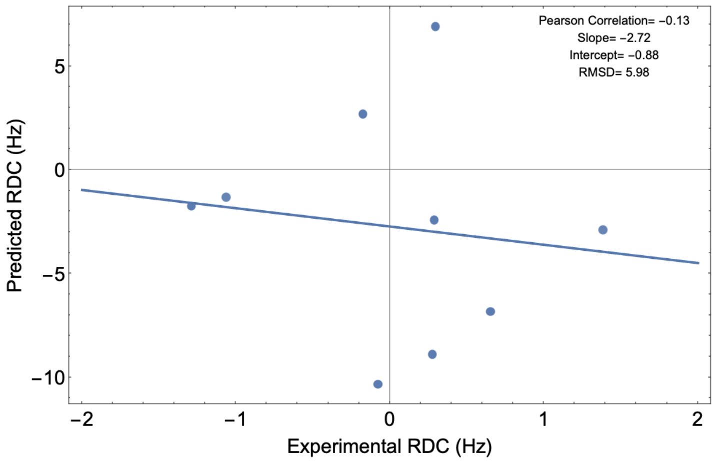


**(B)**

**Figure S7: RDC RMSD, $Q$-factor, and $R_{\mathbf{dip}}$ are not meaningful when RDCs are poor fits to experimental values.** Each panel above shows a different hypothetical set of back-calculated vs. experimental RDCs. The two sets of hypothetical predicted RDCs are proportional to each other. Both sets are completely uncorrelated with experimental RDCs, and neither is a meaningfully better fit to the data than the other. However, the set with smaller range (A) has a much smaller RDC RMSD than the set with larger range (B), and the same would be true for any measure proportional to RMSD (such as $Q$-factor) or approximately proportional to RMSD (such as $R_{\text{dip}}$).

| Predictor ID | $Q$ | $R_{\mathbf{dip}}$ |
|---|---|---|
| 002 | 11.87 | 10.6 |
| 003 | 13.61 | 12.38 |
| 004 | 7.75 | 5.85 |
| 005 | 10.39 | 7.81 |
| 014 | 12.16 | 10.47 |
| 015 | 13.95 | 11.41 |
| 019 | 8.82 | 8.18 |
| 051 | 3.68 | 3.91 |
| 084 | 3.84 | 3.06 |
| 088 | 3.94 | 3.37 |
| 097 | 7.07 | 5.28 |
| 100 | 2.87 | 2.3 |
| 110 | 8.67 | 8.36 |
| 135 | 1.41 | 1.23 |
| 136 | 0.99 | 0.99 |
| 137 | 1.34 | 1.08 |
| 139 | 3.48 | 4.07 |
| 164 | 8.11 | 6.41 |
| 167 | 6.8 | 6.63 |
| 177 | 12.67 | 10.45 |
| 221 | 2.28 | 2.43 |
| 241 | 2.43 | 2.78 |
| 261 | 5.23 | 5.38 |
| 294 | 6.02 | 4.76 |
| 304 | 2.55 | 2.63 |
| 308 | 1.81 | 1.46 |
| 319 | 2.37 | 2.3 |
| 331 | 7.63 | 7.41 |
| 345 | 3.86 | 4.08 |
| 384 | 10.01 | 8.79 |
| 425 | 8.53 | 8.31 |
| 450 | 14.17 | 11.56 |
| 465 | 4.39 | 3.88 |

**Table S2: $Q$-factor and $R_{\text{dip}}$ for each predicted ensemble.** These metrics for comparing structural models to experimental data are not meaningful, because RDCs back-calculated from predictions depart substantially from experimental values and thus are unsuitable here for relative scoring (see Figure S7).

| Predictor ID | Pearson correlation | Slope (Hz/Hz) | $y$-intercept (Hz) | Anisotropicity (kcal/mol) |
|---|---|---|---|---|
| 014 | 0.43 | 5.42 | 0.93 | 2.45 |
| 015 | 0.07 | 0.99 | -0.40 | 1.90 |
| 019 | 0.75 | 7.11 | 0.38 | 2.19 |
| 051 | -0.04 | -0.15 | 0.43 | 1.10 |
| 097 | 0.18 | 1.16 | -1.97 | 0.80 |
| 110 | 0.64 | 5.86 | 0.61 | 2.06 |
| 135 | -0.76 | -0.38 | 0.03 | 0.04 |
| 136 | 0.16 | 0.06 | 0.17 | 0.08 |
| 137 | -0.41 | -0.23 | -0.05 | 0.07 |
| 164 | 0.24 | 1.90 | -2.13 | 1.33 |
| 167 | 0.15 | 0.94 | 1.72 | 1.42 |
| 177 | 0.45 | 5.59 | -3.32 | 1.87 |
| 221 | 0.07 | 0.14 | 0.08 | 0.59 |
| 241 | 0.13 | 0.31 | 0.09 | 0.66 |
| 261 | -0.23 | -1.14 | 0.68 | 1.51 |
| 294 | -0.55 | -2.59 | -1.39 | 1.32 |
| 304 | 0.31 | 0.84 | -0.03 | 0.77 |
| 308 | -0.17 | -0.23 | 0.12 | 0.19 |
| 319 | -0.13 | -0.26 | 0.11 | 0.55 |
| 331 | 0.81 | 6.88 | 0.12 | 2.02 |
| 345 | 0.73 | 3.31 | 0.12 | 1.04 |
| 384 | 0.63 | 6.84 | -1.11 | 2.06 |
| 425 | 0.81 | 7.60 | -0.16 | 2.22 |
| 450 | 0.12 | 1.70 | -0.54 | 2.25 |
| 465 | 0.64 | 3.20 | -0.65 | 1.37 |

**Table S3: Pearson correlation, slope, $y$-intercept (Section 2.2.2) and anisotropicity (Section 2.2.3) for each predictor.** These values were compared to ideal values and normalized as described in Section 2.2.4 with resulting values in Table S4.

| Predictor ID | Chebyshev distance | Limiting quantity | Normalized distance, Pearson correlation | Normalized distance, slope | Normalized distance, $y$-intercept | Normalized distance, anisotropicity |
|---|---|---|---|---|---|---|
| 014 | 0.98 | Anisotropicity | 0.28 | 0.28 | 0.11 | 0.98 |
| 015 | 0.71 | Anisotropicity | 0.47 | 0.00 | 0.05 | 0.71 |
| 019 | 0.85 | Anisotropicity | 0.13 | 0.39 | 0.04 | 0.85 |
| 051 | 0.52 | Pearson Correlation | 0.52 | 0.07 | 0.05 | 0.33 |
| 097 | 0.41 | Pearson Correlation | 0.41 | 0.01 | 0.22 | 0.18 |
| 110 | 0.79 | Anisotropicity | 0.18 | 0.31 | 0.07 | 0.79 |
| 135 | 0.88 | Pearson Correlation | 0.88 | 0.09 | 0.00 | 0.03 |
| 136 | 0.42 | Pearson Correlation | 0.42 | 0.06 | 0.02 | 0.01 |
| 137 | 0.70 | Pearson Correlation | 0.70 | 0.08 | 0.01 | 0.01 |
| 164 | 0.44 | Anisotropicity | 0.38 | 0.06 | 0.24 | 0.44 |
| 167 | 0.48 | Anisotropicity | 0.43 | 0.00 | 0.20 | 0.48 |
| 177 | 0.70 | Anisotropicity | 0.28 | 0.30 | 0.38 | 0.70 |
| 221 | 0.47 | Pearson Correlation | 0.47 | 0.06 | 0.01 | 0.08 |
| 241 | 0.44 | Pearson Correlation | 0.44 | 0.04 | 0.01 | 0.12 |
| 261 | 0.61 | Pearson Correlation | 0.61 | 0.14 | 0.08 | 0.53 |
| 294 | 0.78 | Pearson Correlation | 0.78 | 0.23 | 0.16 | 0.43 |
| 304 | 0.35 | Pearson Correlation | 0.35 | 0.01 | 0.00 | 0.17 |
| 308 | 0.58 | Pearson Correlation | 0.58 | 0.08 | 0.01 | 0.00 |
| 319 | 0.56 | Pearson Correlation | 0.56 | 0.08 | 0.01 | 0.06 |
| 331 | 0.77 | Anisotropicity | 0.09 | 0.38 | 0.01 | 0.77 |
| 345 | 0.30 | Anisotropicity | 0.14 | 0.15 | 0.01 | 0.30 |
| 384 | 0.79 | Anisotropicity | 0.18 | 0.38 | 0.13 | 0.79 |
| 425 | 0.87 | Anisotropicity | 0.10 | 0.42 | 0.02 | 0.87 |
| 450 | 0.88 | Anisotropicity | 0.44 | 0.05 | 0.06 | 0.88 |
| 465 | 0.46 | Anisotropicity | 0.18 | 0.14 | 0.07 | 0.46 |

**Table S4: Components of the Chebyshev distance used to rank each predictor.** The last four columns relate to the quantities by which the predicted ensembles were compared with RDC data (see Section 2.2.2 and Table S3). As described in Section 2.2.4, the raw values of these quantities are normalized according to their worst possible values, and the largest of these four determines the Chebyshev distance (Section 2.2.4).

| Alignments | $S_{xx} \times 10^4$ | $S_{yy} \times 10^4$ | $S_{zz} \times 10^4$ | $S_{xy} \times 10^4$ | $S_{xz} \times 10^4$ | $S_{yz} \times 10^4$ |
|---|---|---|---|---|---|---|
| Z Domain (OLC1) | -8.6551 | 14.9983 | -6.34323 | -20.5640 | -0.41839 | 0.36701 |
| Z Domain (OLC2) | 5.04792 | -2.43413 | -2.61378 | -1.55826 | 2.56783 | 2.28739 |
| C Domain (OLC1) | -0.521156 | 1.57063 | -1.04947 | 0.771357 | 0.0267095 | 0.524178 |
| C Domain (OLC2) | 0.357427 | -0.54729 | 0.18986 | 0.036935 | -0.0785261 | -0.454419 |

**Table S5: The first two OLC Saupe tensors from Qi et al. [3] for each domain are used in this analysis.**

### S7 Graphical summaries of analyses for each predicted ensemble

This section contains a figure for each predicted ensemble, in order of prediction number, with panels summarizing key analyses (Figures S8 to S33). Summary statistics of the data in (Figures S8 to S33) are given in Figures 6 and 9, Figures S4 and S6, and Tables 2 to S4. Here (Figures S8 to S33) they are broken down and visualized on a per-predictor basis. In addition, Table S1 gives the predictor name for each prediction number, along with methods used and number of structures in each predicted ensemble.

For each figure in this section, panels include:

(A) Predicted vs. experimental SAXS $I(q)$ curves for target T1200 (Section 3.2). The top plot shows the experimental $I(q)$ curve in red and in blue shows the corresponding curve back-calculated from the predicted ensemble. For an ideal prediction (and setting aside the noise in the experimental curve), the red and blue curves would line up precisely. Because even small deviations in these curves are significant, the lower plot of this panel shows the difference between the back-calculated and experimental $I(q)$ curves, after normalizing by the standard deviation of the data as a function of scattering vector magnitude $q$. For an ideal prediction, the blue line would lie exactly on the red dashed zero line, indicating zero discrepancy between predictor's model and the SAXS experimental measurement.

(B) Similar to (A), for target T1300.

(C) Disk-on-Sphere (DoS) visualization [3] of predicted distribution of interdomain orientations, calculated from predicted ensemble using a kernel density estimate (Section 2.2.3), for target T1200.

(D) Similar to (C), for target T1300.

(E) DoS visualization of the solution CDIO, which is a mixture of our Solution 1 and Solution 2 CDIOs (Figure S5). The proportions of this mixture are defined by the $\alpha$ parameter shown in the panel caption (Section 2.2.3). $\alpha$ is chosen for each predictor such that the free energy difference between the corresponding solution CDIO and the kernelized predicted ensemble in panel (C) is minimized (Sections 2.2.3 and S4).

(F) Comparison of back-calculated and experimental NMR RDCs, for target T1200 (Section 2.2.2). The green line shows a linear regression of back-calculated vs. experimental values, and the red line shows the linear regression for a perfect prediction in which back-calculated and experimental RDCs were equal. We invite predictors to compare the fits in this panel to main text Figure 2, which shows the fit to our continuous distribution CDIO models.

(G) Color legend for panels (C), (D), and (E). As in Figure S5, colors indicate percentiles; a given orientation $R$ is assigned a color such that the corresponding percentage of total probability mass in the distribution is found at orientations less likely than $R$. These legends are reproduced in each figure for convenience; the same color mapping is used both throughout this paper as well as in Qi et al. [3].

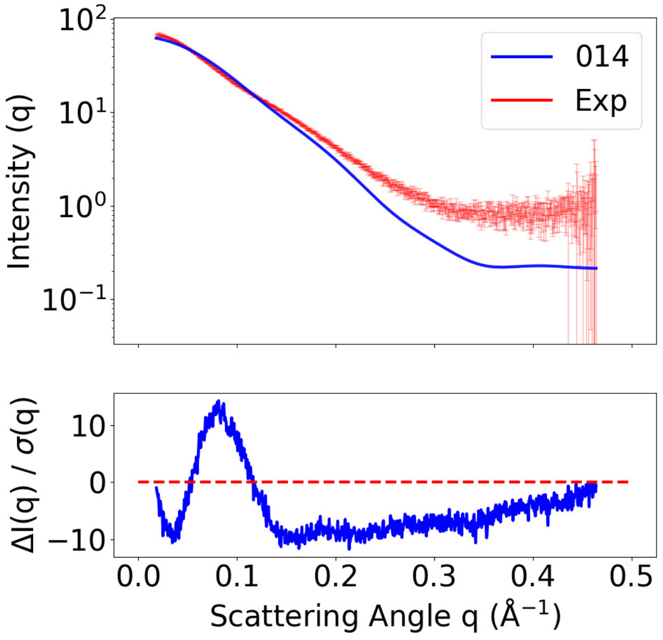


**(A)** SAXS curves, T1200

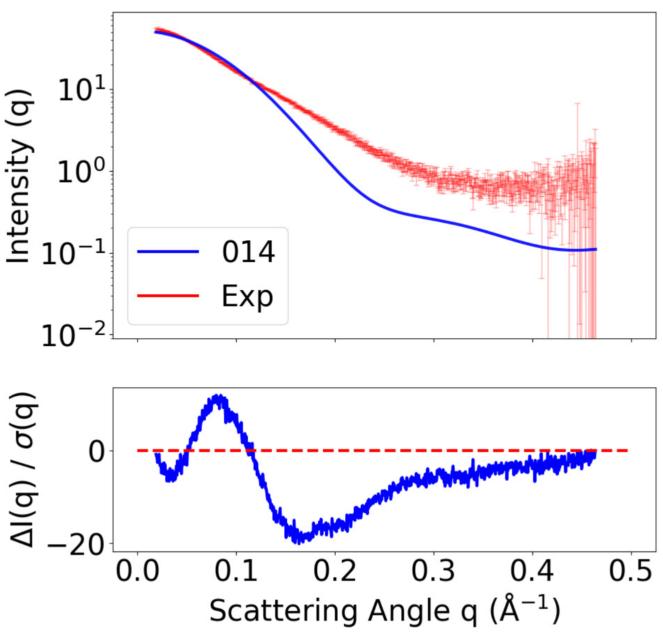


**(B)** SAXS curves, T1300

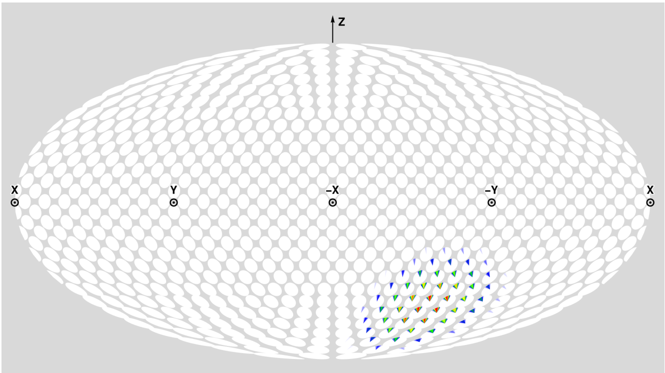


**(C)** Prediction DoS, T1200

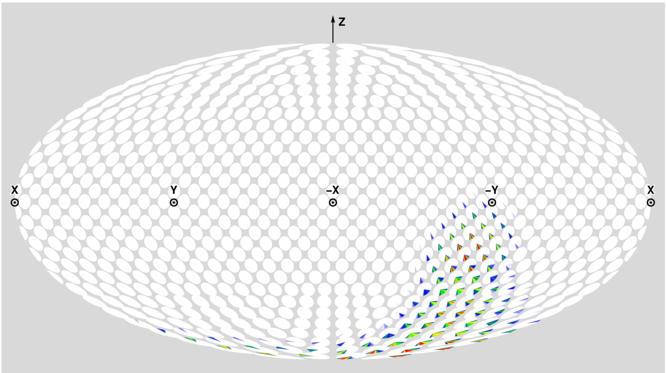


**(D)** Prediction DoS, T1300

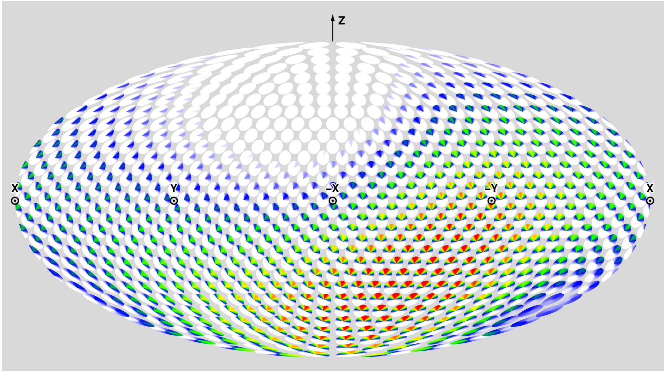


**(E)** Solution CDIO DoS ($\alpha = 1.00$)

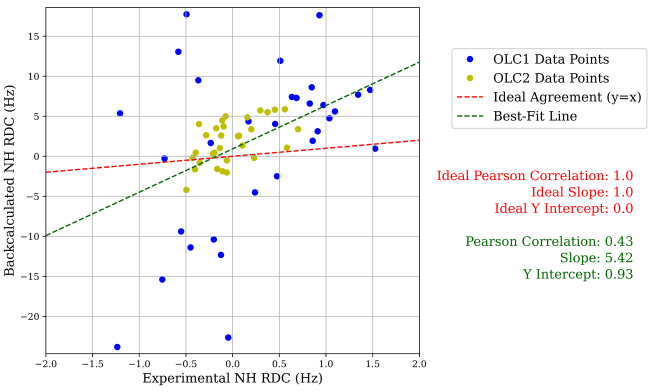


**(F)** Prediction RMB Plot, T1200

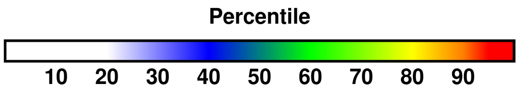


**(G)** Color percentiles for DoS

**Figure S8: Summary of analysis for predictor Cool-PSP (014).** For T1200 and T1300 respectively, (A) and (B) compare back-calculated and experimental SAXS data in the form of $I(q)$ curves and residuals, with bars showing experimental standard deviation (Section 3.2). For T1200 and T1300 respectively, (C) and (D) show disk-on-sphere (DoS) visualizations of kernel density estimates of predicted ensembles (Section 2.2.3). For T1200, (E) shows the solution CDIO for $\alpha = 1.00$, which is the parameter corresponding to the smallest free energy difference with the predicted ensemble (Section 2.2.3). For T1200, (F) compares back-calculated and experimental NMR RDC data (Section 2.2.2). (G) shows the percentile color legend for (C), (D), and (E).

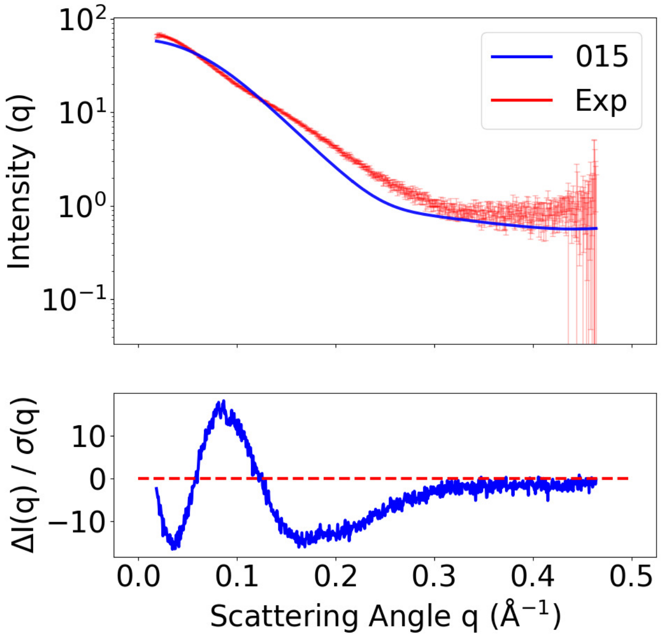


**(A)** SAXS curves, T1200

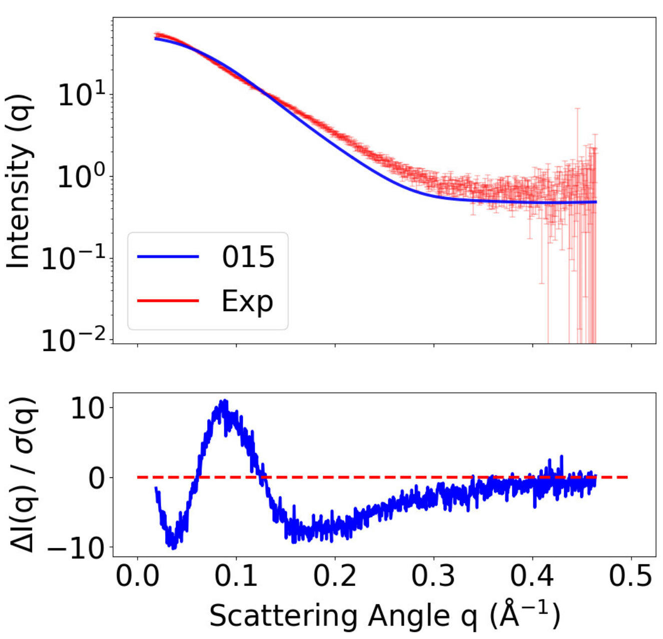


**(B)** SAXS curves, T1300

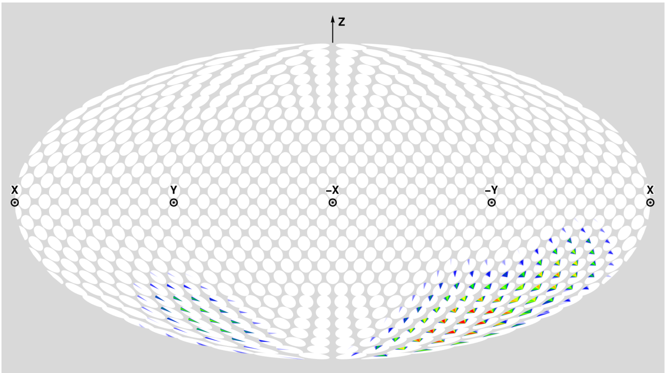


**(C)** Prediction DoS, T1200

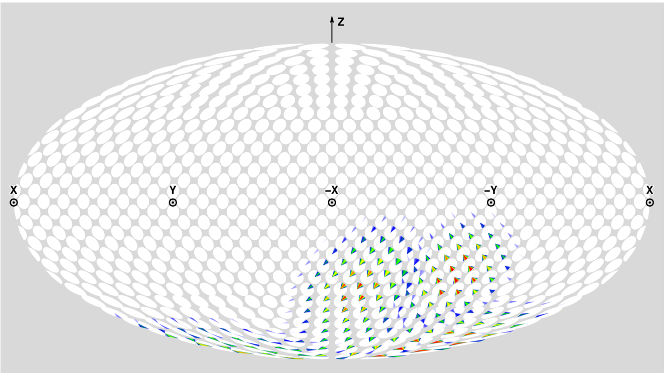


**(D)** Prediction DoS, T1300

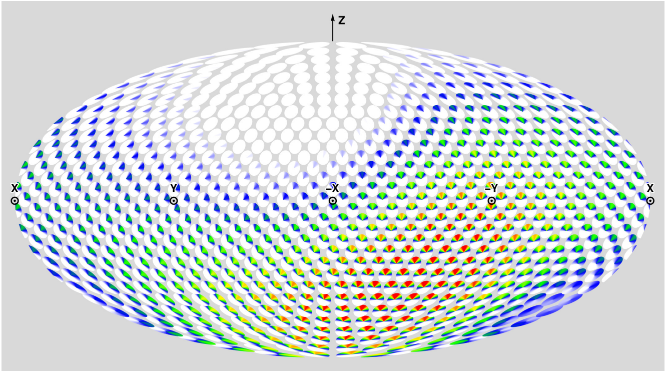


**(E)** Solution CDIO DoS ($\alpha = 1.00$)

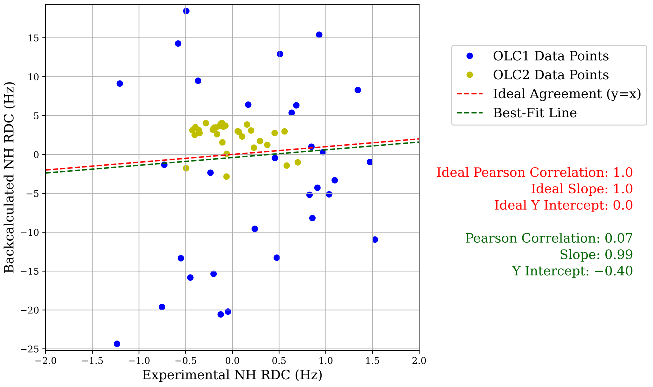


**(F)** Prediction RMB Plot, T1200

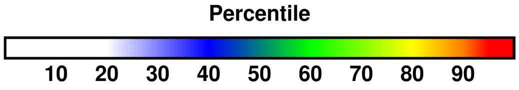


**(G)** Color percentiles for DoS

**Figure S9: Summary of analysis for predictor PEZYFoldings (015).** For T1200 and T1300 respectively, (A) and (B) compare back-calculated and experimental SAXS data in the form of $I(q)$ curves and residuals, with bars showing experimental standard deviation (Section 3.2). For T1200 and T1300 respectively, (C) and (D) show disk-on-sphere (DoS) visualizations of kernel density estimates of predicted ensembles (Section 2.2.3). For T1200, (E) shows the solution CDIO for $\alpha = 1.00$, which is the parameter corresponding to the smallest free energy difference with the predicted ensemble (Section 2.2.3). For T1200, (F) compares back-calculated and experimental NMR RDC data (Section 2.2.2). (G) shows the percentile color legend for (C), (D), and (E).

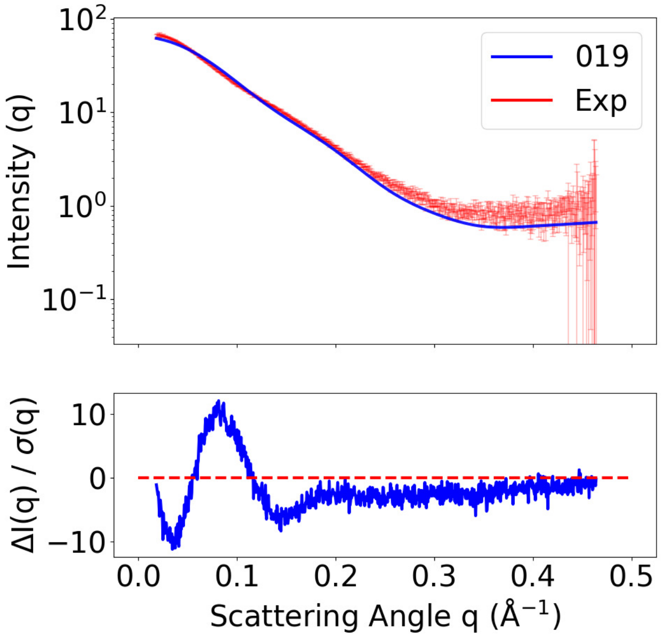


**(A)** SAXS curves, T1200

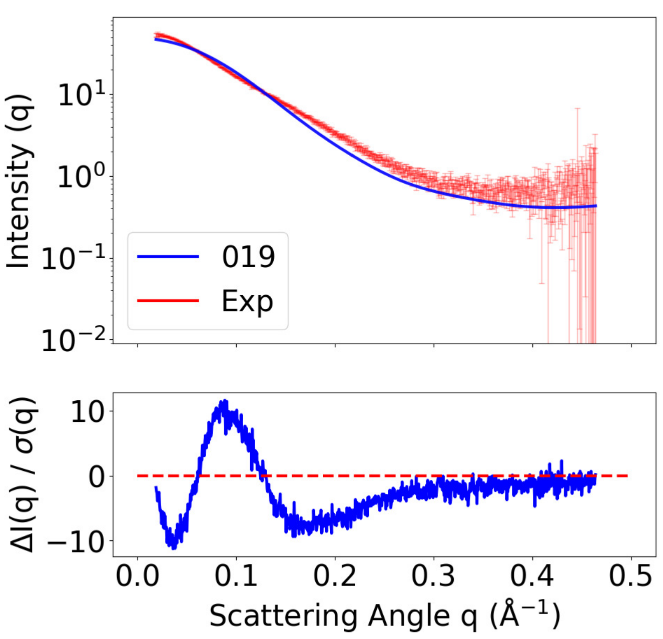


**(B)** SAXS curves, T1300

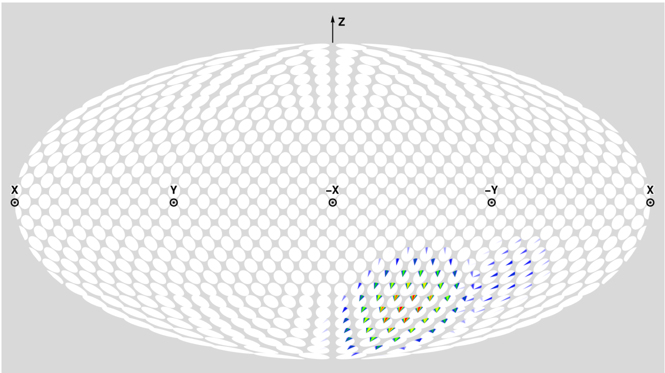


**(C)** Prediction DoS, T1200

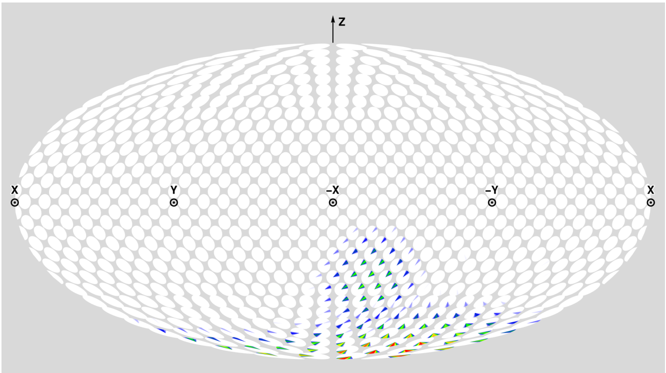


**(D)** Prediction DoS, T1300

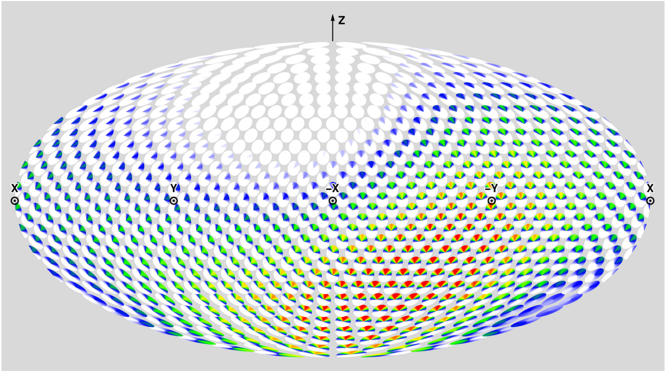


**(E)** Solution CDIO DoS ($\alpha = 1.00$)

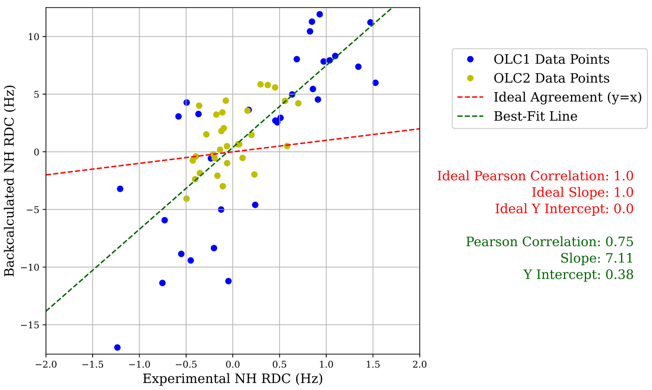


**(F)** Prediction RMB Plot, T1200

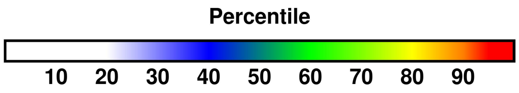


**(G)** Color percentiles for DoS

**Figure S10: Summary of analysis for predictor Zheng-Server (019)**. For T1200 and T1300 respectively, (A) and (B) compare back-calculated and experimental SAXS data in the form of $I(q)$ curves and residuals, with bars showing experimental standard deviation (Section 3.2). For T1200 and T1300 respectively, (C) and (D) show disk-on-sphere (DoS) visualizations of kernel density estimates of predicted ensembles (Section 2.2.3). For T1200, (E) shows the solution CDIO for $\alpha = 1.00$, which is the parameter corresponding to the smallest free energy difference with the predicted ensemble (Section 2.2.3). For T1200, (F) compares back-calculated and experimental NMR RDC data (Section 2.2.2). (G) shows the percentile color legend for (C), (D), and (E).

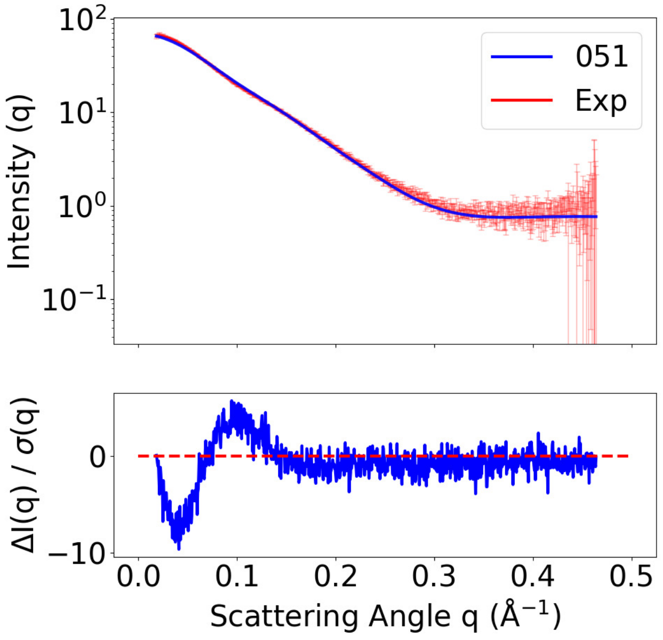


**(A)** SAXS curves, T1200

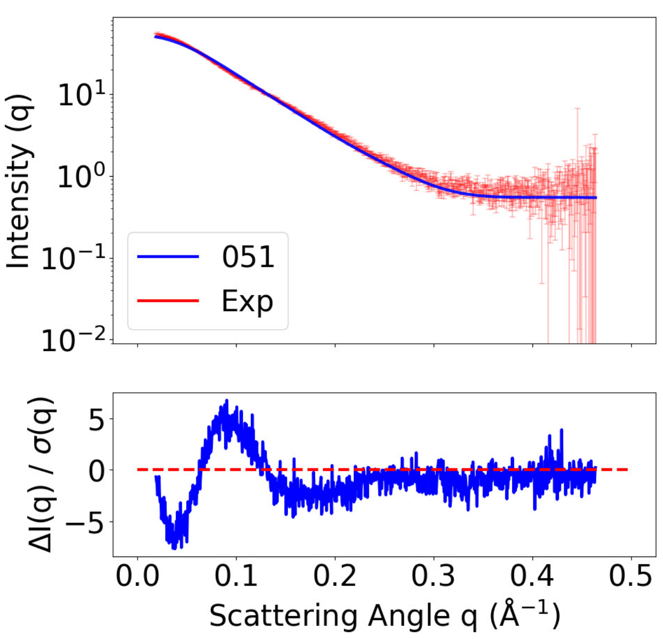


**(B)** SAXS curves, T1300

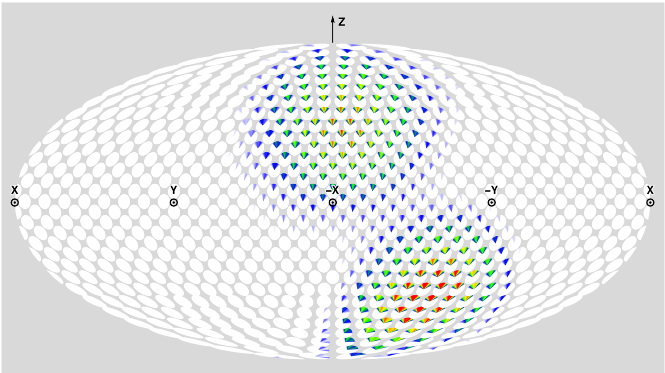


**(C)** Prediction DoS, T1200

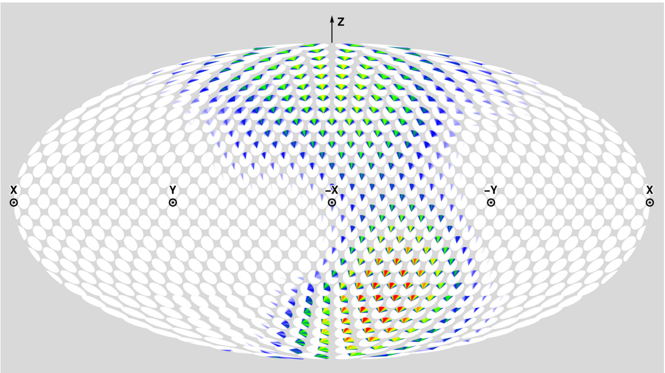


**(D)** Prediction DoS, T1300

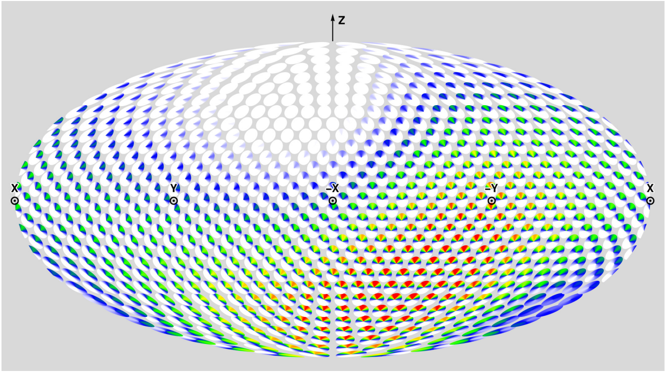


**(E)** Solution CDIO DoS ($\alpha = 0.93$)

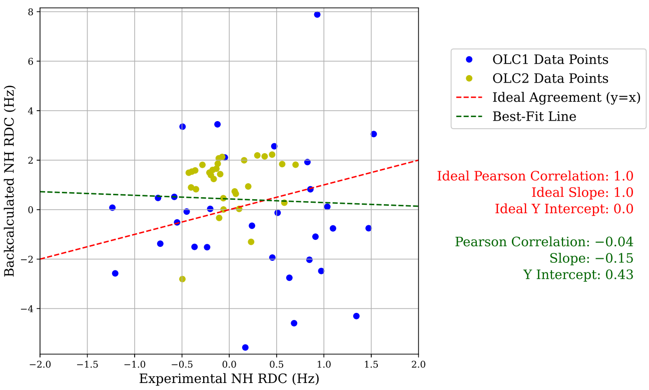


**(F)** Prediction RMB Plot, T1200

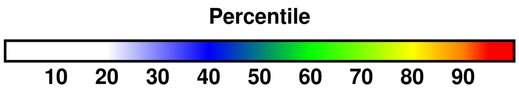


**(G)** Color percentiles for DoS

**Figure S11: Summary of analysis for predictor MULTICOM (051).** For T1200 and T1300 respectively, (A) and (B) compare back-calculated and experimental SAXS data in the form of $I(q)$ curves and residuals, with bars showing experimental standard deviation (Section 3.2). For T1200 and T1300 respectively, (C) and (D) show disk-on-sphere (DoS) visualizations of kernel density estimates of predicted ensembles (Section 2.2.3). For T1200, (E) shows the solution CDIO for $\alpha = 0.93$, which is the parameter corresponding to the smallest free energy difference with the predicted ensemble (Section 2.2.3). For T1200, (F) compares back-calculated and experimental NMR RDC data (Section 2.2.2). (G) shows the percentile color legend for (C), (D), and (E).

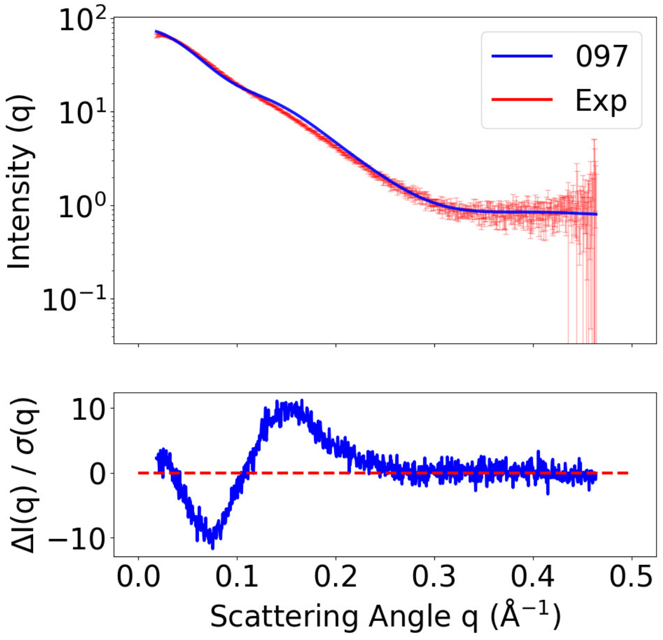


**(A)** SAXS curves, T1200

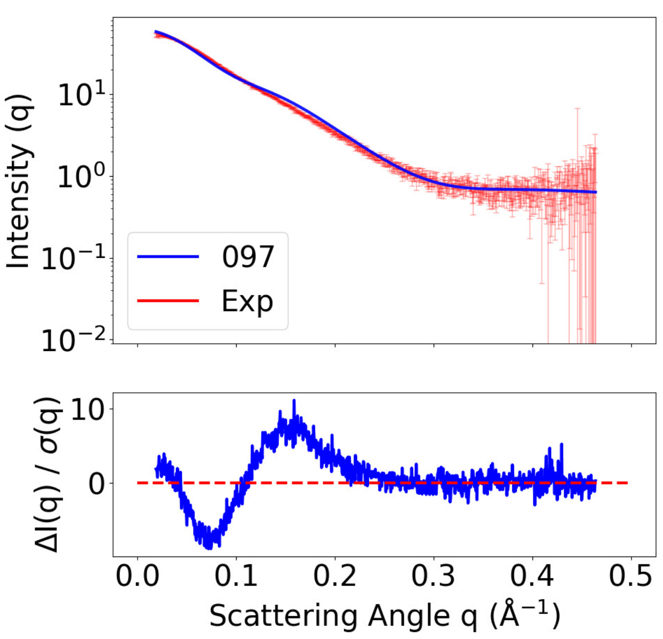


**(B)** SAXS curves, T1300

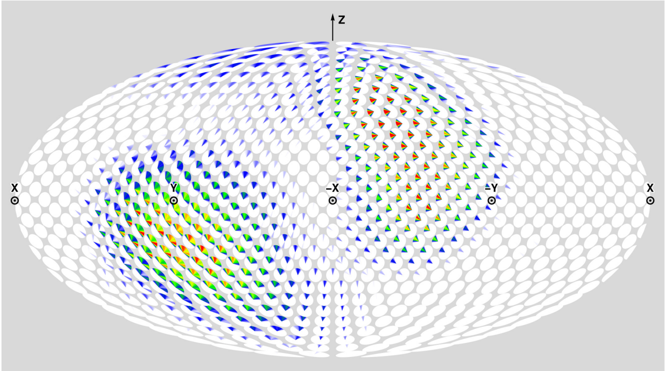

**(C)** Prediction DoS, T1200

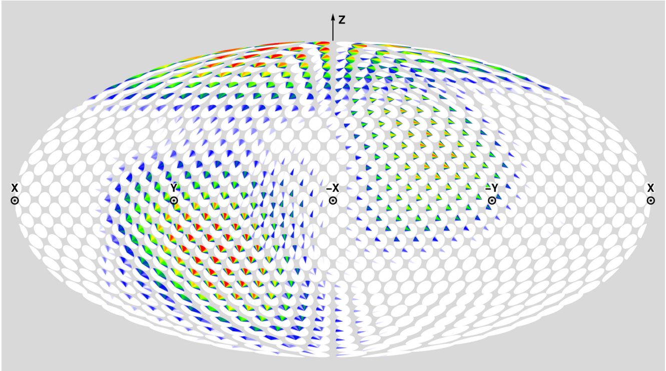

**(D)** Prediction DoS, T1300

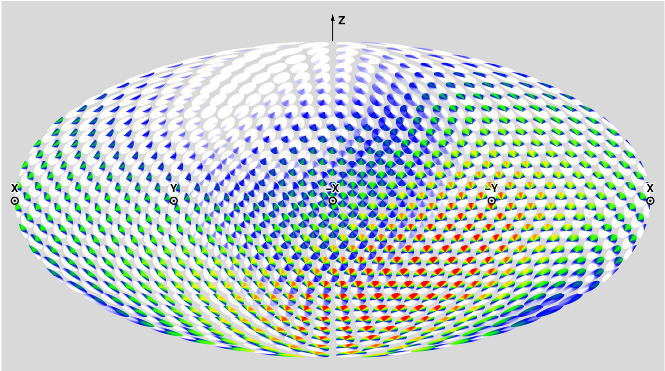

**(E)** Solution CDIO DoS ($\alpha = 0.80$)

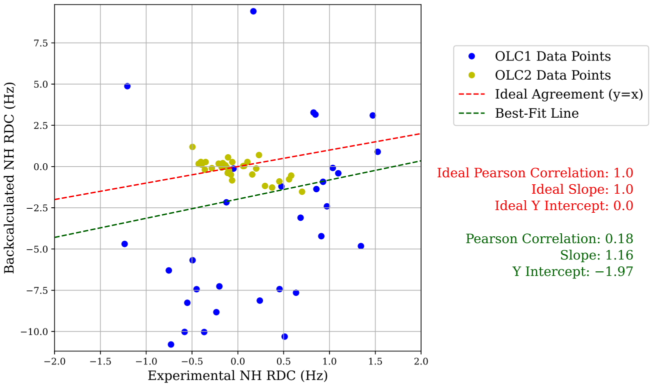


**(F)** Prediction RMB Plot, T1200

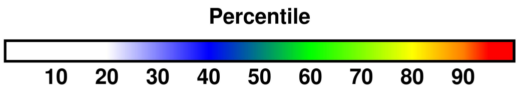


**(G)** Color percentiles for DoS

**Figure S12: Summary of analysis for predictor JFK-THG-IDPCONFGEN (097).** For T1200 and T1300 respectively, (A) and (B) compare back-calculated and experimental SAXS data in the form of $I(q)$ curves and residuals, with bars showing experimental standard deviation (Section 3.2). For T1200 and T1300 respectively, (C) and (D) show disk-on-sphere (DoS) visualizations of kernel density estimates of predicted ensembles (Section 2.2.3). For T1200, (E) shows the solution CDIO for $\alpha = 0.80$, which is the parameter corresponding to the smallest free energy difference with the predicted ensemble (Section 2.2.3). For T1200, (F) compares back-calculated and experimental NMR RDC data (Section 2.2.2). (G) shows the percentile color legend for (C), (D), and (E).

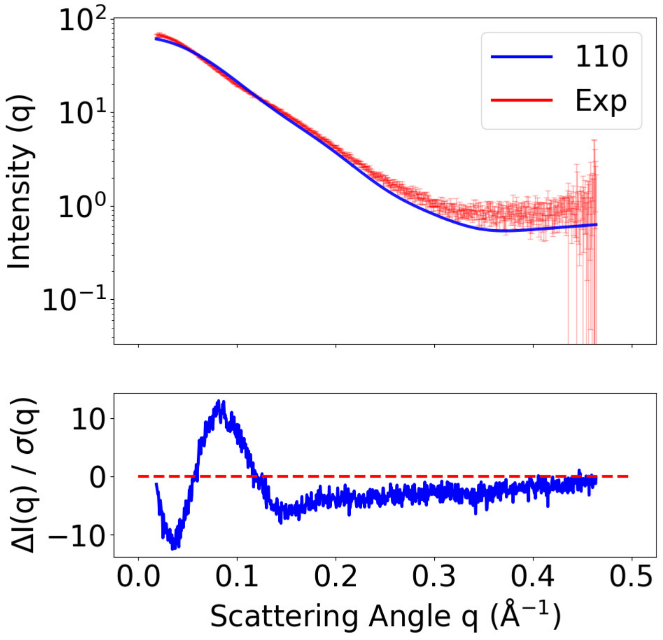


**(A)** SAXS curves, T1200

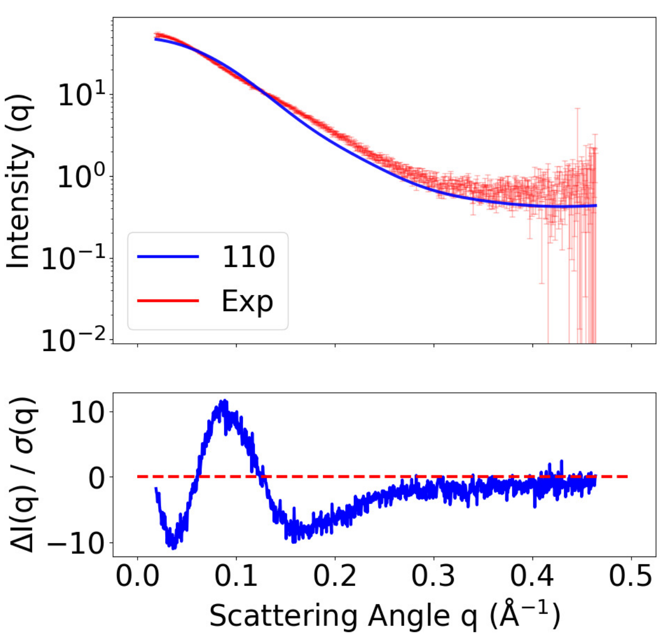


**(B)** SAXS curves, T1300

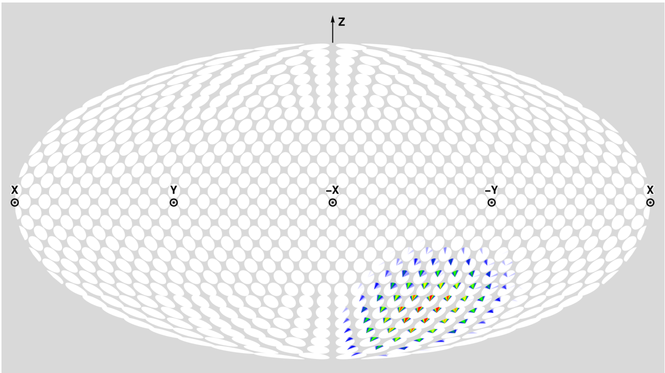


**(C)** Prediction DoS, T1200

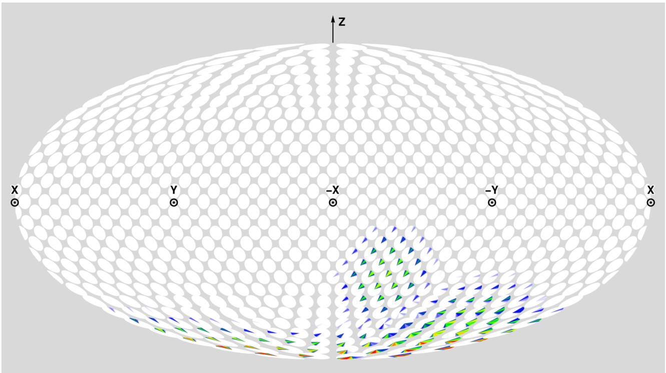


**(D)** Prediction DoS, T1300

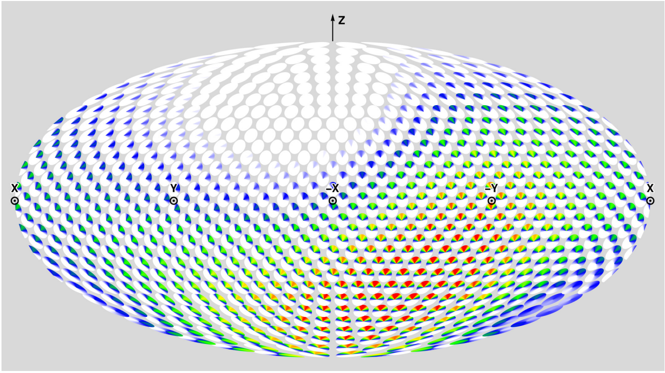


**(E)** Solution CDIO DoS ($\alpha = 1.00$)

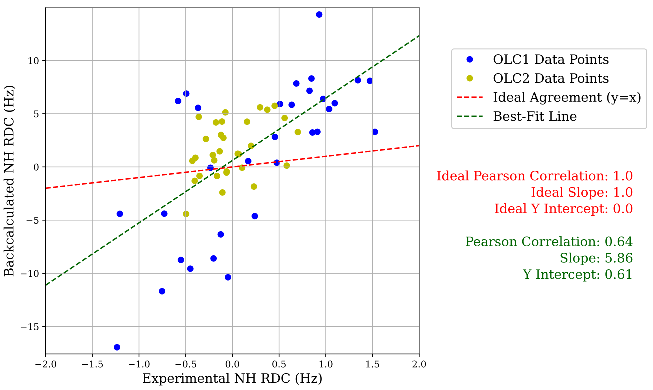


**(F)** Prediction RMB Plot, T1200

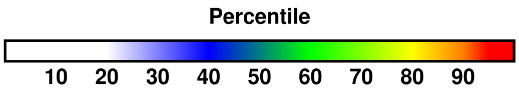


**(G)** Color percentiles for DoS

**Figure S13: Summary of analysis for predictor MIEnsembles-Server/Zheng-Multimer/Zheng (110)**. For T1200 and T1300 respectively, (A) and (B) compare back-calculated and experimental SAXS data in the form of $I(q)$ curves and residuals, with bars showing experimental standard deviation (Section 3.2). For T1200 and T1300 respectively, (C) and (D) show disk-on-sphere (DoS) visualizations of kernel density estimates of predicted ensembles (Section 2.2.3). For T1200, (E) shows the solution CDIO for $\alpha = 1.00$, which is the parameter corresponding to the smallest free energy difference with the predicted ensemble (Section 2.2.3). For T1200, (F) compares back-calculated and experimental NMR RDC data (Section 2.2.2). (G) shows the percentile color legend for (C), (D), and (E).

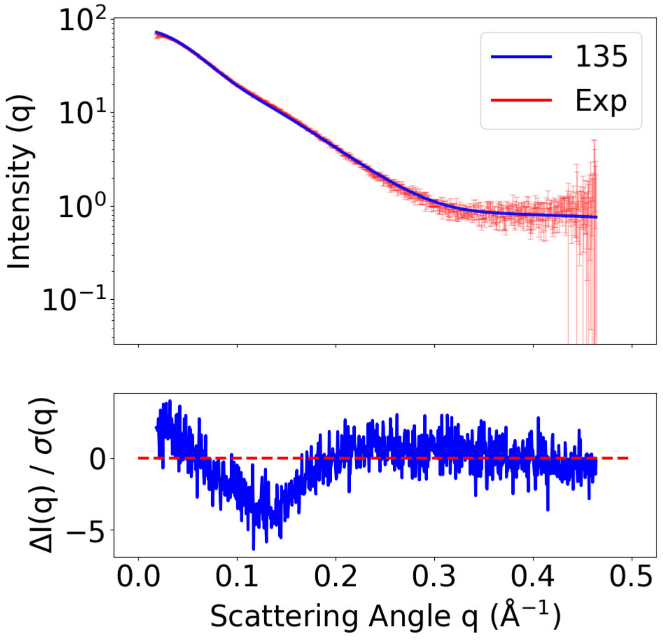


**(A)** SAXS curves, T1200

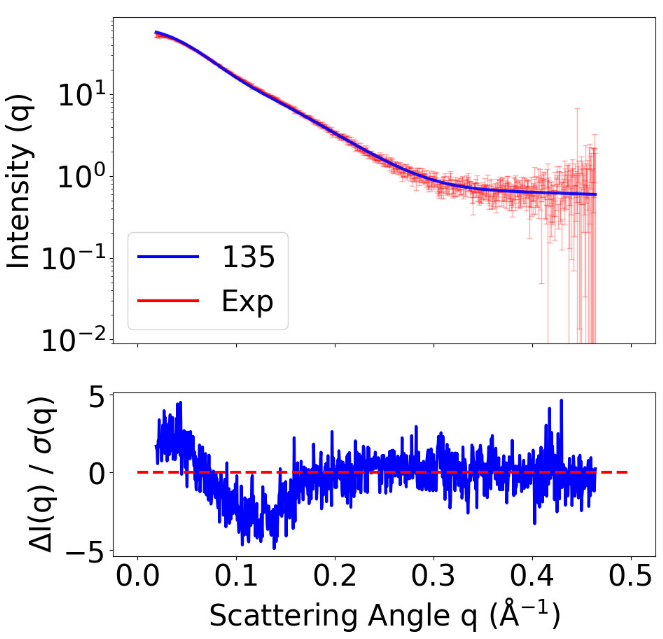


**(B)** SAXS curves, T1300

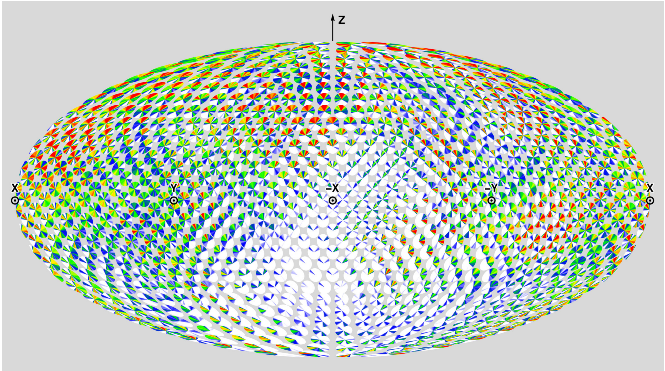

**(C)** Prediction DoS, T1200

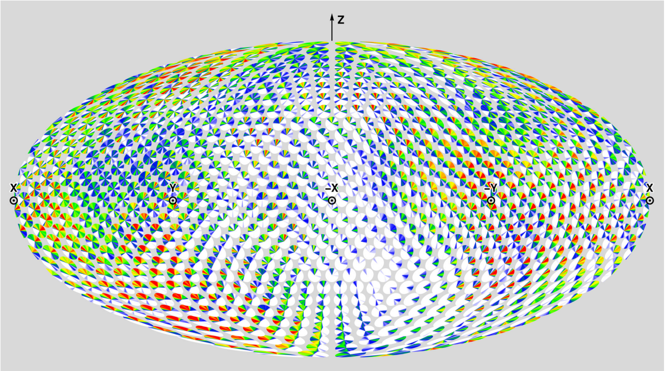

**(D)** Prediction DoS, T1300

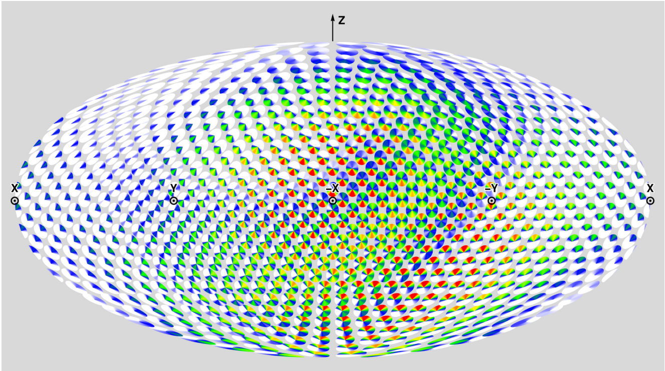

**(E)** Solution CDIO DoS ($\alpha = 0.54$)

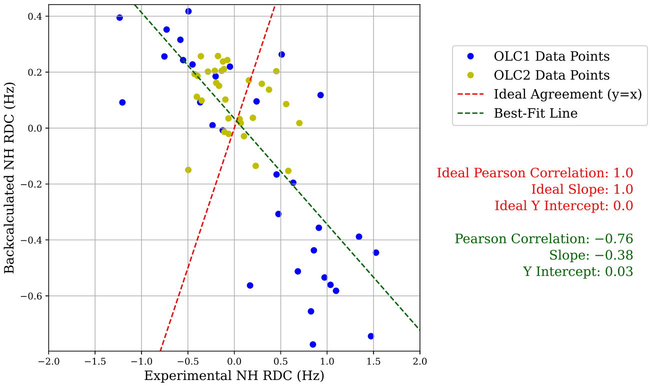


**(F)** Prediction RMB Plot, T1200

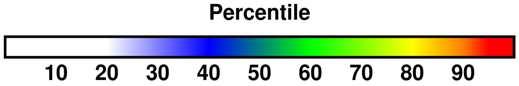


**(G)** Color percentiles for DoS

**Figure S14: Summary of analysis for predictor Lindorff-Larsenclvds (135).** For T1200 and T1300 respectively, (A) and (B) compare back-calculated and experimental SAXS data in the form of $I(q)$ curves and residuals, with bars showing experimental standard deviation (Section 3.2). For T1200 and T1300 respectively, (C) and (D) show disk-on-sphere (DoS) visualizations of kernel density estimates of predicted ensembles (Section 2.2.3). For T1200, (E) shows the solution CDIO for $\alpha = 0.54$, which is the parameter corresponding to the smallest free energy difference with the predicted ensemble (Section 2.2.3). For T1200, (F) compares back-calculated and experimental NMR RDC data (Section 2.2.2). (G) shows the percentile color legend for (C), (D), and (E).

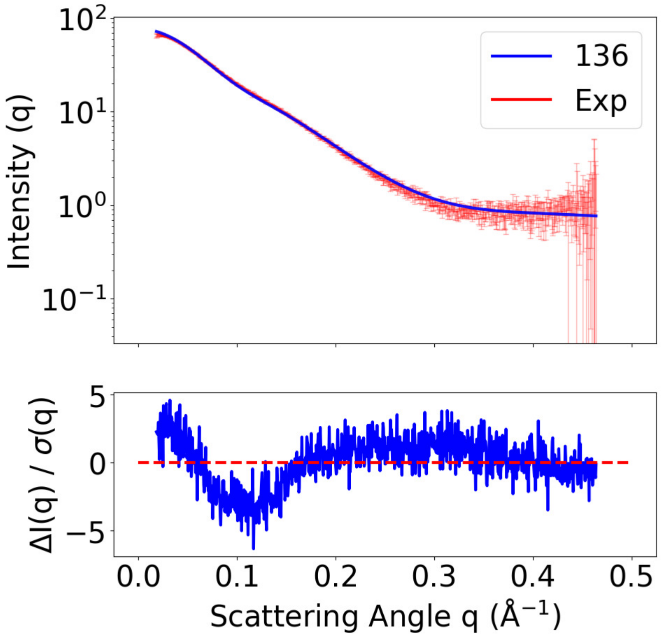


**(A)** SAXS curves, T1200

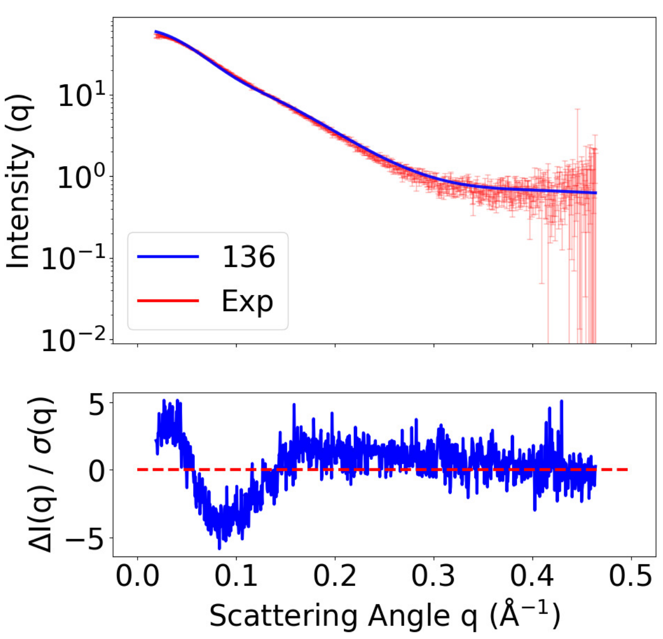


**(B)** SAXS curves, T1300

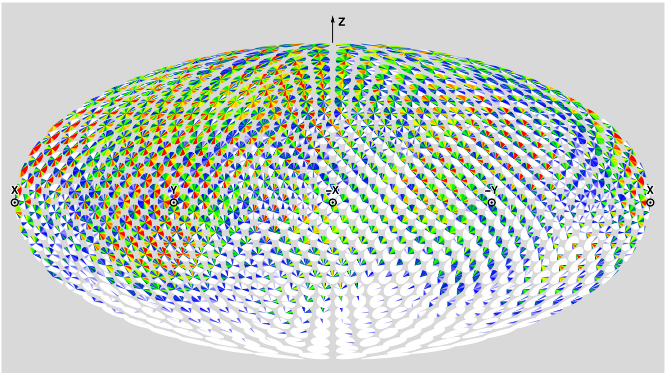

**(C)** Prediction DoS, T1200

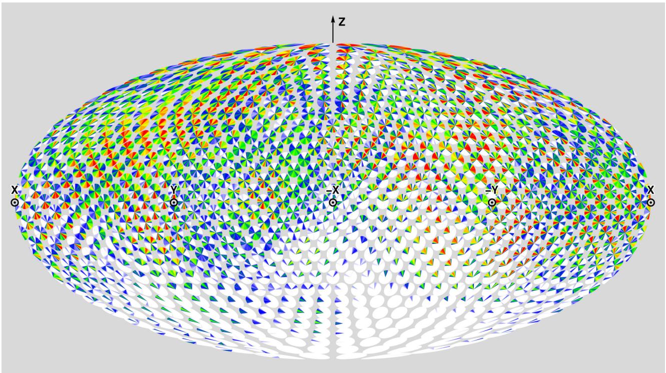

**(D)** Prediction DoS, T1300

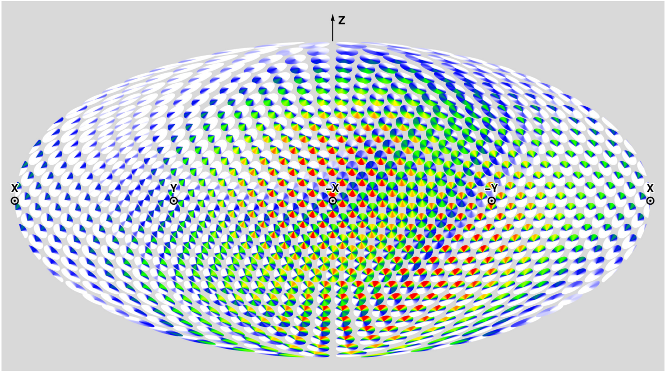

**(E)** Solution CDIO DoS ($\alpha = 0.54$)

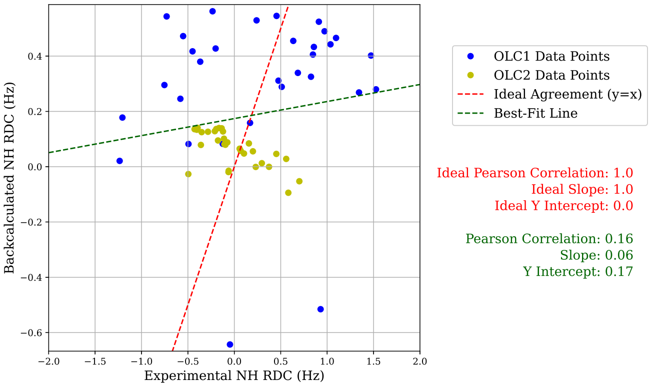


**(F)** Prediction RMB Plot, T1200

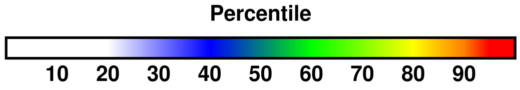


**(G)** Color percentiles for DoS

**Figure S15: Summary of analysis for predictor Lindorff-LarsenM3PPS (136)**. For T1200 and T1300 respectively, (A) and (B) compare back-calculated and experimental SAXS data in the form of $I(q)$ curves and residuals, with bars showing experimental standard deviation (Section 3.2). For T1200 and T1300 respectively, (C) and (D) show disk-on-sphere (DoS) visualizations of kernel density estimates of predicted ensembles (Section 2.2.3). For T1200, (E) shows the solution CDIO for $\alpha = 0.54$, which is the parameter corresponding to the smallest free energy difference with the predicted ensemble (Section 2.2.3). For T1200, (F) compares back-calculated and experimental NMR RDC data (Section 2.2.2). (G) shows the percentile color legend for (C), (D), and (E).

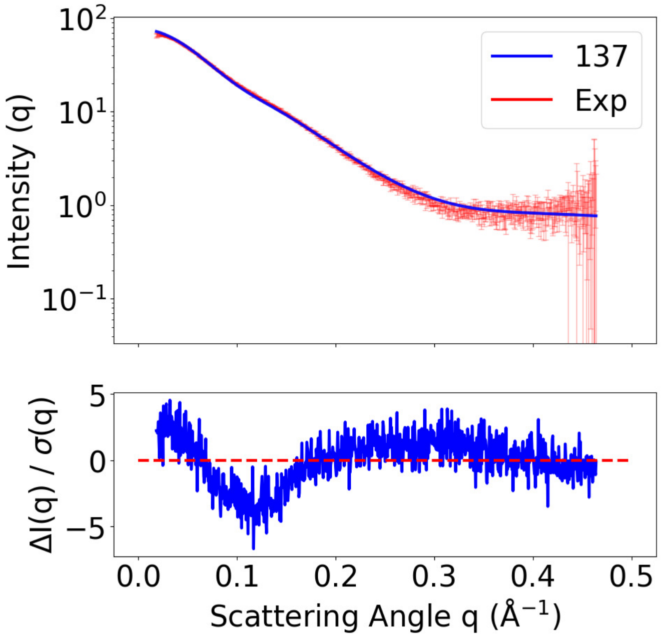


**(A)** SAXS curves, T1200

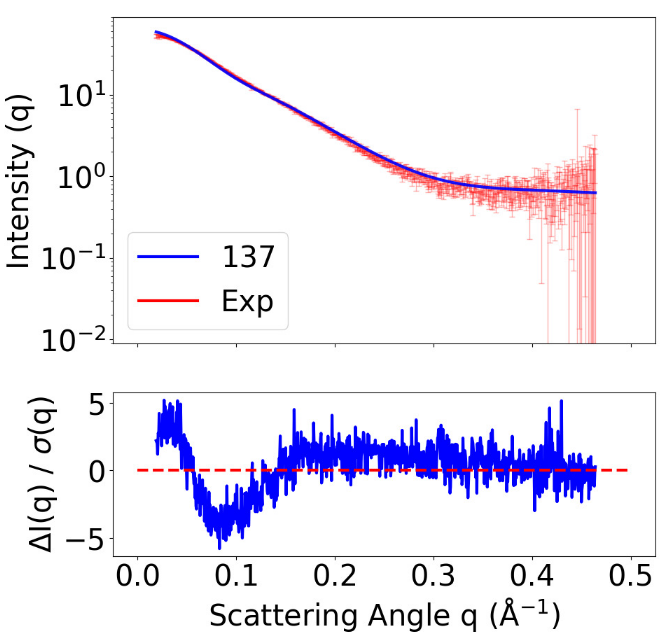


**(B)** SAXS curves, T1300

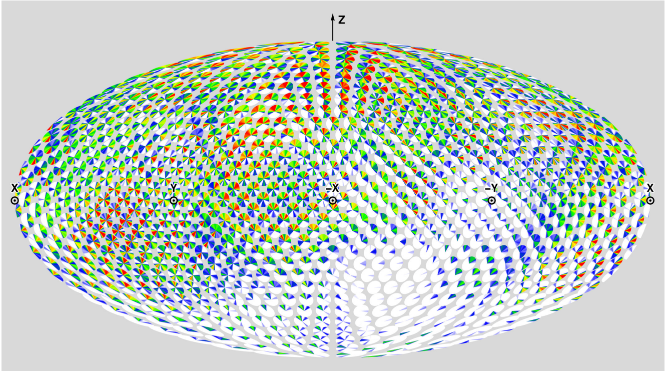

**(C)** Prediction DoS, T1200

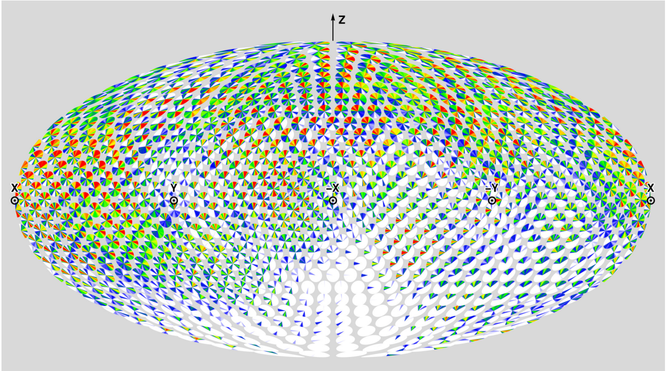

**(D)** Prediction DoS, T1300

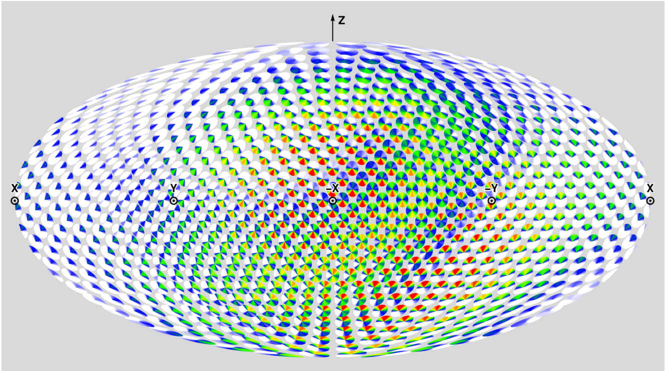

**(E)** Solution CDIO DoS ($\alpha = 0.53$)

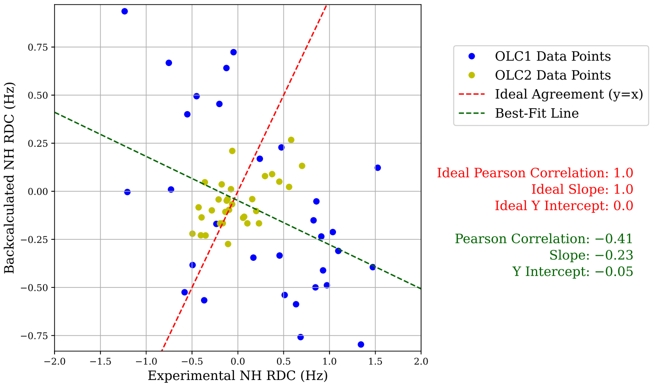


**(F)** Prediction RMB Plot, T1200

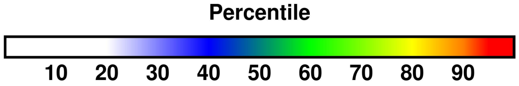


**(G)** Color percentiles for DoS

**Figure S16: Summary of analysis for predictor Lindorff-Larsen**M3PWS **(137)**. For T1200 and T1300 respectively, (A) and (B) compare back-calculated and experimental SAXS data in the form of $I(q)$ curves and residuals, with bars showing experimental standard deviation (Section 3.2). For T1200 and T1300 respectively, (C) and (D) show disk-on-sphere (DoS) visualizations of kernel density estimates of predicted ensembles (Section 2.2.3). For T1200, (E) shows the solution CDIO for $\alpha = 0.53$, which is the parameter corresponding to the smallest free energy difference with the predicted ensemble (Section 2.2.3). For T1200, (F) compares back-calculated and experimental NMR RDC data (Section 2.2.2). (G) shows the percentile color legend for (C), (D), and (E).

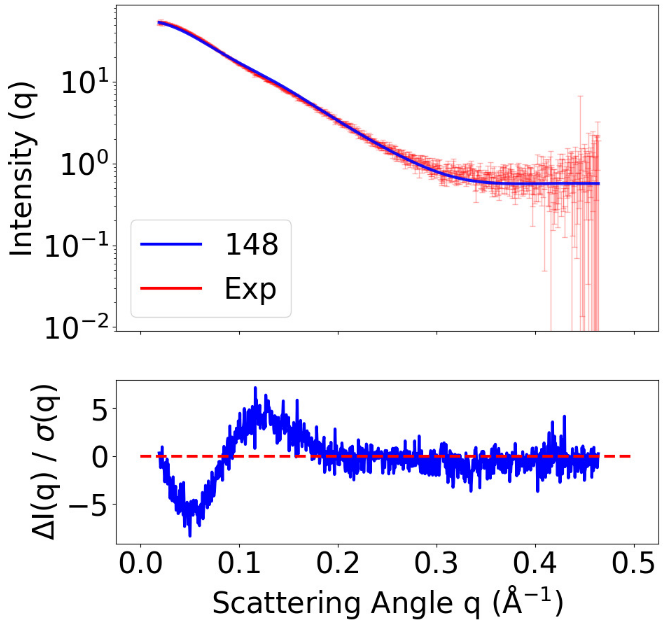


**(A)** SAXS curves, T1300

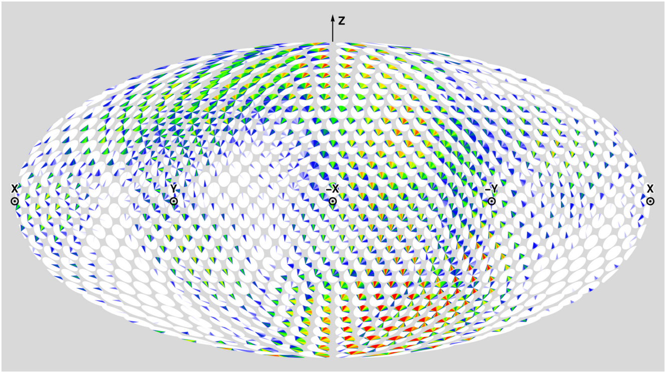


**(B)** Prediction DoS, T1300

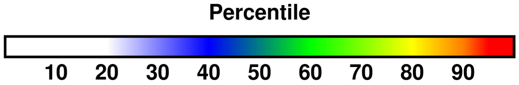


**(C)** Color percentiles for DoS

**Figure S17: Summary of analysis for predictor GuijunLab-Complex (148)**. For predictor 148, only the T1300 submission was assessed (Section 3.1). For T1300, (A) compares back-calculated and experimental SAXS data in the form of an $I(q)$ curve and residuals, with bars showing experimental standard deviation (Section 3.2). For T1300, (B) shows a disk-on-sphere (DoS) visualization of the kernel density estimate of the predicted ensemble (Section 2.2.3). (C) shows the percentile color legend for (B).

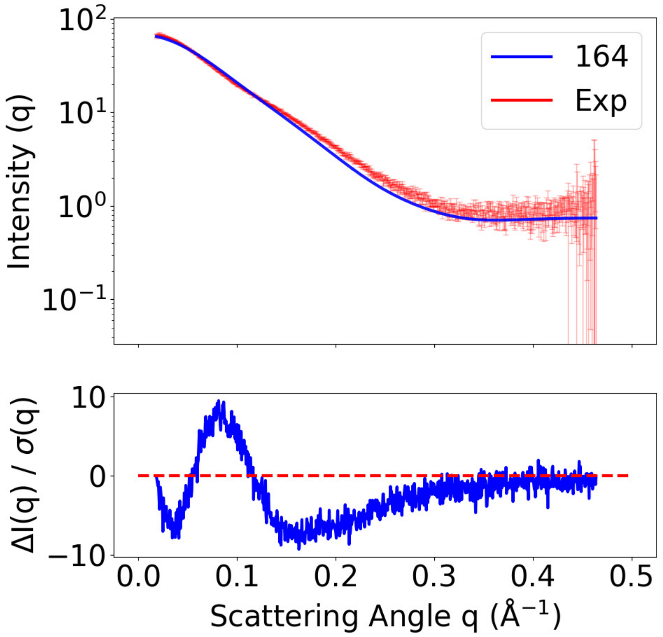


**(A)** SAXS curves, T1200

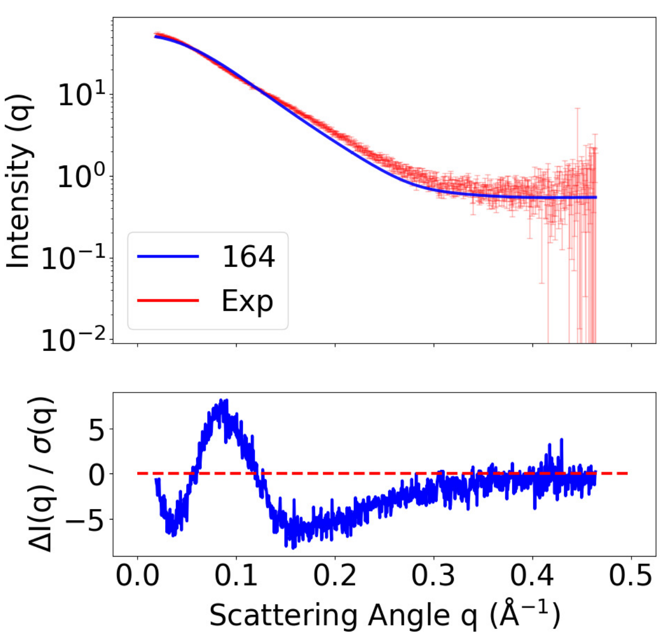


**(B)** SAXS curves, T1300

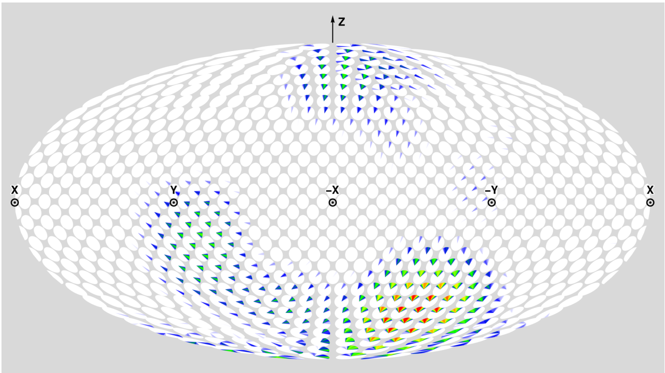

**(C)** Prediction DoS, T1200

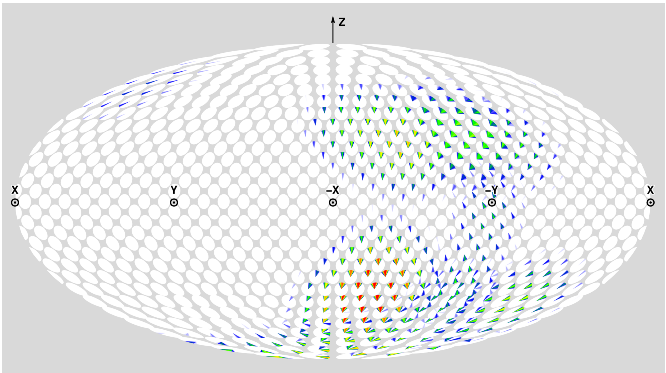

**(D)** Prediction DoS, T1300

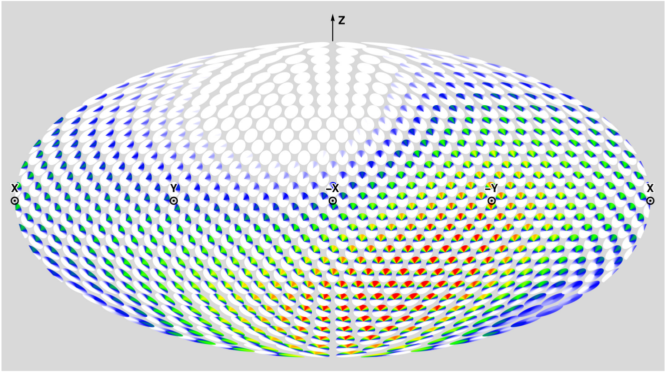

**(E)** Solution CDIO DoS ($\alpha = 1.00$)

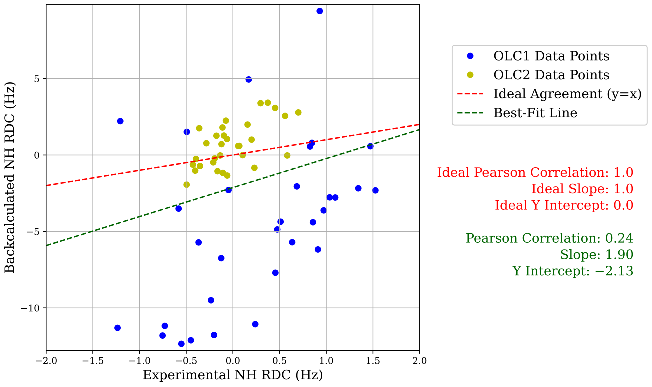


**(F)** Prediction RMB Plot, T1200

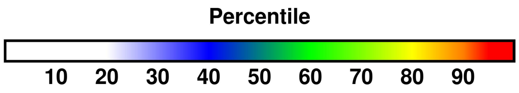


**(G)** Color percentiles for DoS

**Figure S18: Summary of analysis for predictor McGuffin (164).** For T1200 and T1300 respectively, (A) and (B) compare back-calculated and experimental SAXS data in the form of $I(q)$ curves and residuals, with bars showing experimental standard deviation (Section 3.2). For T1200 and T1300 respectively, (C) and (D) show disk-on-sphere (DoS) visualizations of kernel density estimates of predicted ensembles (Section 2.2.3). For T1200, (E) shows the solution CDIO for $\alpha = 1.00$, which is the parameter corresponding to the smallest free energy difference with the predicted ensemble (Section 2.2.3). For T1200, (F) compares back-calculated and experimental NMR RDC data (Section 2.2.2). (G) shows the percentile color legend for (C), (D), and (E).

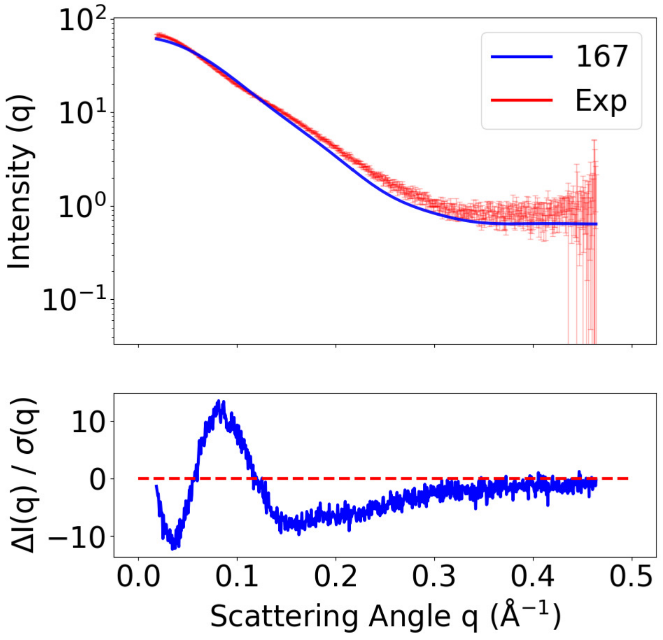


**(A)** SAXS curves, T1200

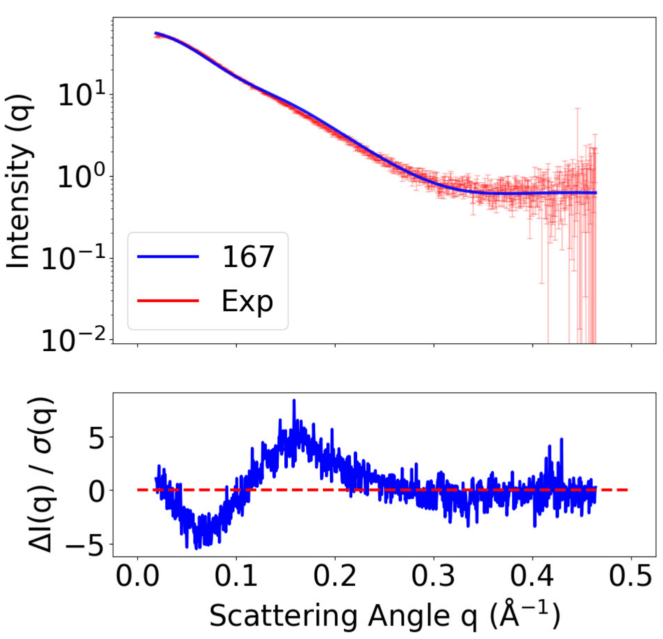


**(B)** SAXS curves, T1300

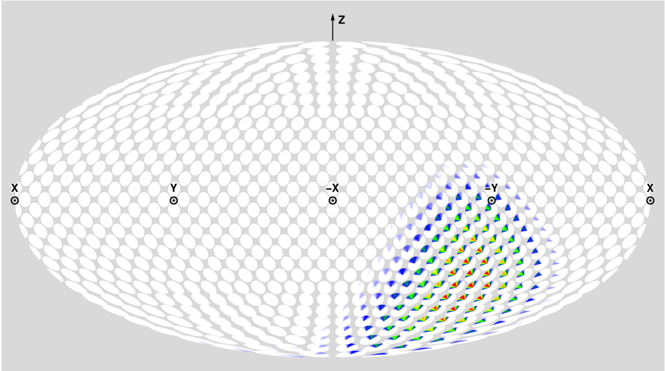

**(C)** Prediction DoS, T1200

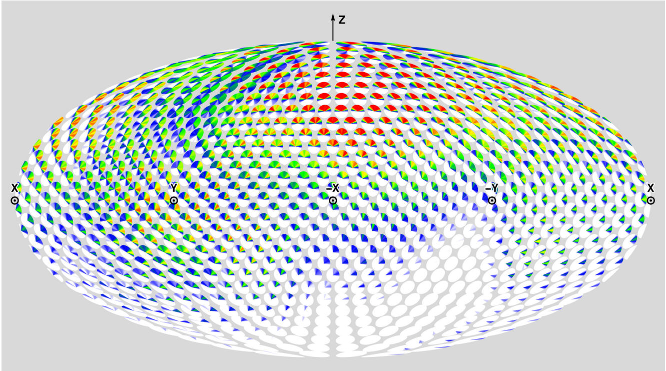

**(D)** Prediction DoS, T1300

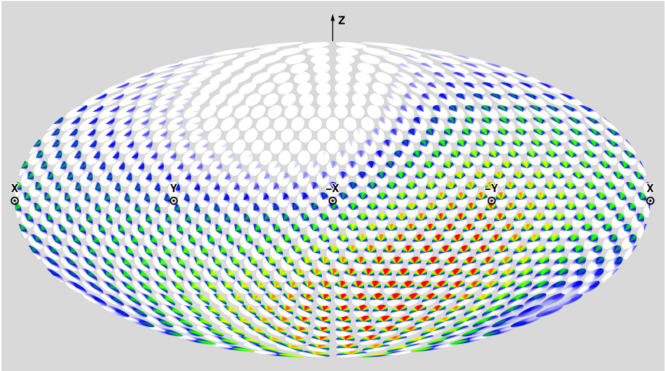

**(E)** Solution CDIO DoS ($\alpha = 1.00$)

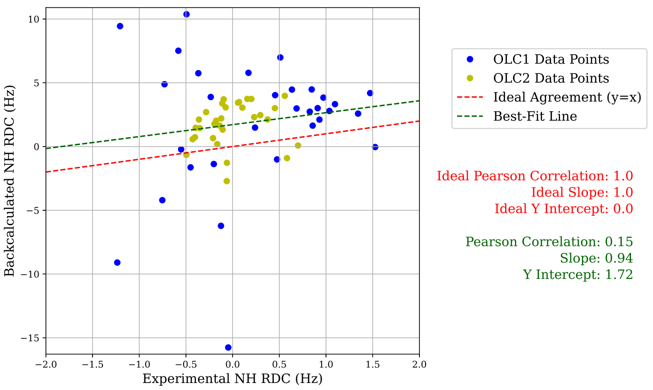


**(F)** Prediction RMB Plot, T1200

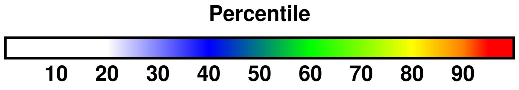


**(G)** Color percentiles for DoS

**Figure S19: Summary of analysis for predictor OpenComplex (167).** For T1200 and T1300 respectively, (A) and (B) compare back-calculated and experimental SAXS data in the form of $I(q)$ curves and residuals, with bars showing experimental standard deviation (Section 3.2). For T1200 and T1300 respectively, (C) and (D) show disk-on-sphere (DoS) visualizations of kernel density estimates of predicted ensembles (Section 2.2.3). For T1200, (E) shows the solution CDIO for $\alpha = 1.00$, which is the parameter corresponding to the smallest free energy difference with the predicted ensemble (Section 2.2.3). For T1200, (F) compares back-calculated and experimental NMR RDC data (Section 2.2.2). (G) shows the percentile color legend for (C), (D), and (E).

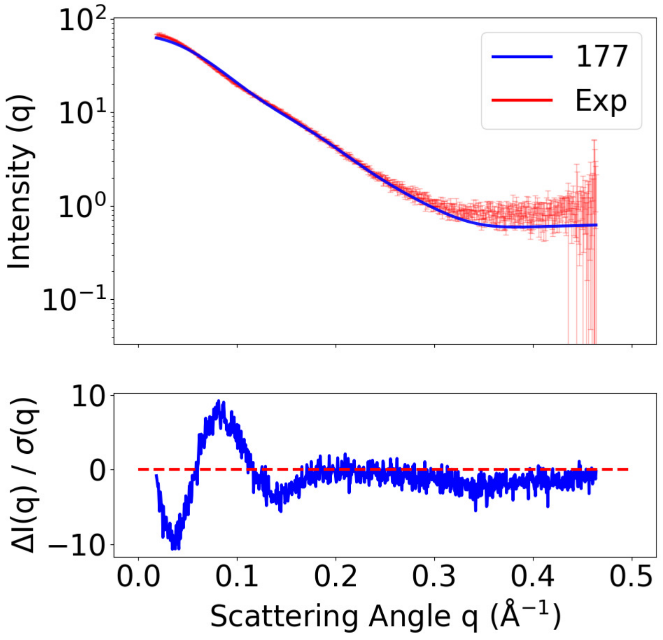


**(A)** SAXS curves, T1200

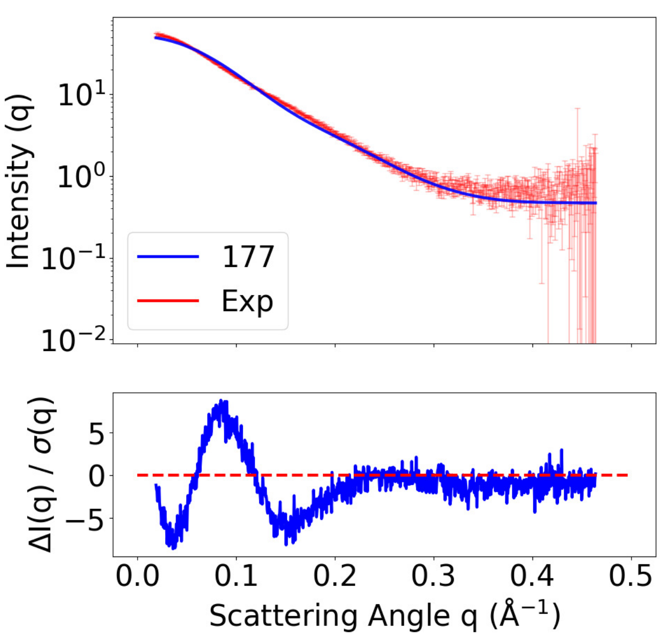


**(B)** SAXS curves, T1300

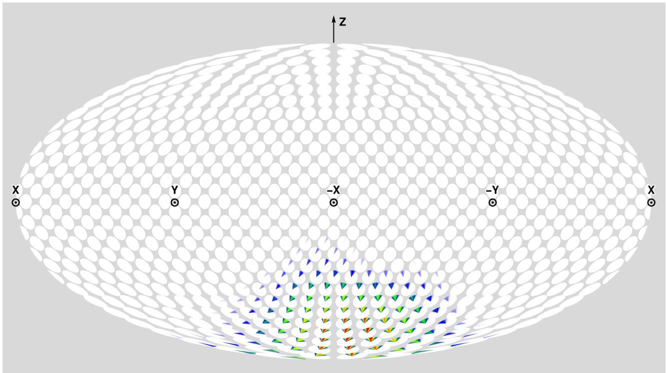


**(C)** Prediction DoS, T1200

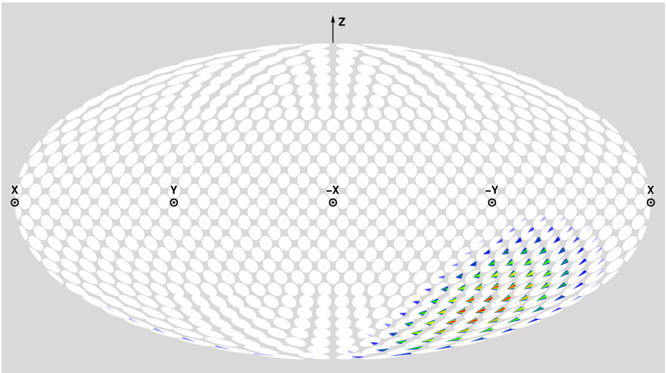


**(D)** Prediction DoS, T1300

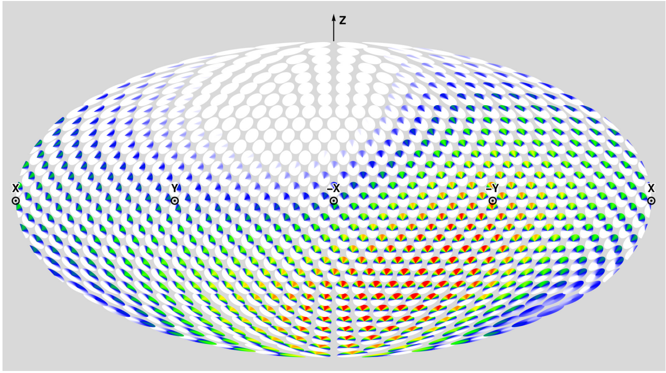


**(E)** Solution CDIO DoS ($\alpha = 1.00$)

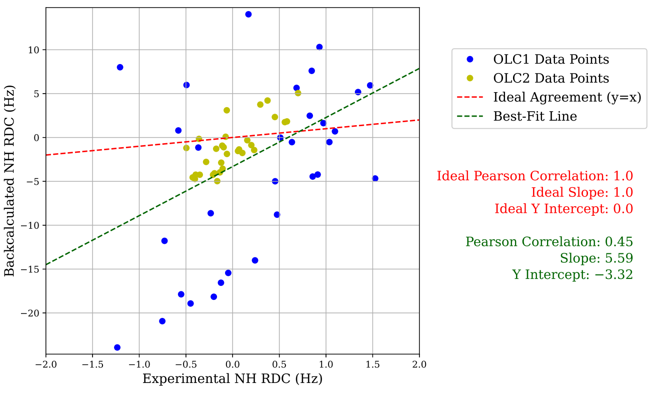


**(F)** Prediction RMB Plot, T1200

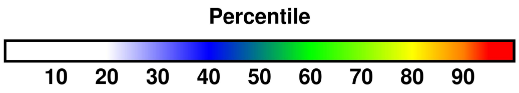


**(G)** Color percentiles for DoS

**Figure S20: Summary of analysis for predictor aicb (177)**. For T1200 and T1300 respectively, (A) and (B) compare back-calculated and experimental SAXS data in the form of $I(q)$ curves and residuals, with bars showing experimental standard deviation (Section 3.2). For T1200 and T1300 respectively, (C) and (D) show disk-on-sphere (DoS) visualizations of kernel density estimates of predicted ensembles (Section 2.2.3). For T1200, (E) shows the solution CDIO for $\alpha = 1.00$, which is the parameter corresponding to the smallest free energy difference with the predicted ensemble (Section 2.2.3). For T1200, (F) compares back-calculated and experimental NMR RDC data (Section 2.2.2). (G) shows the percentile color legend for (C), (D), and (E).

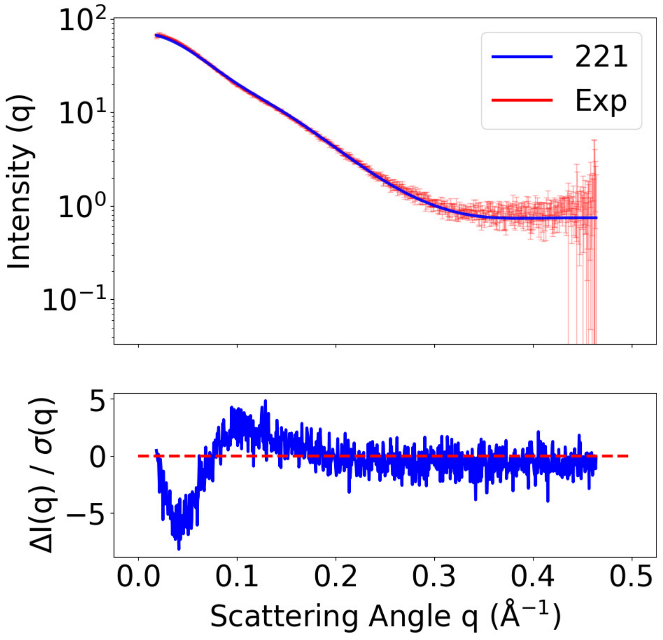


**(A)** SAXS curves, T1200

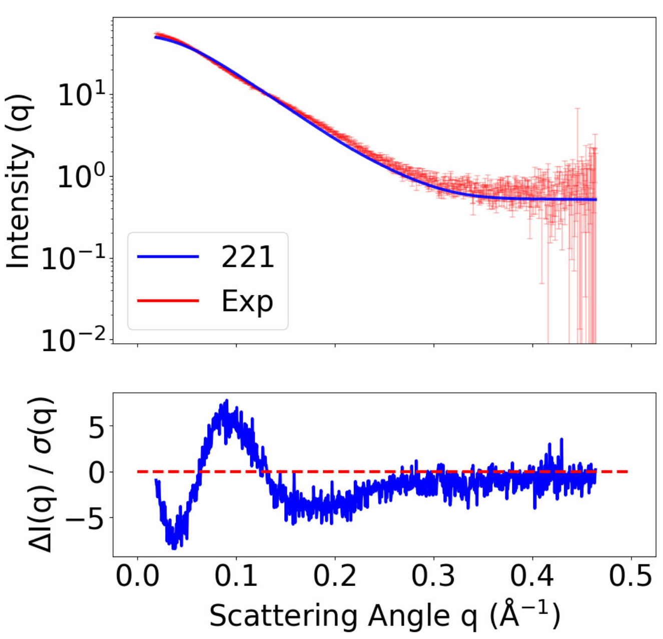


**(B)** SAXS curves, T1300

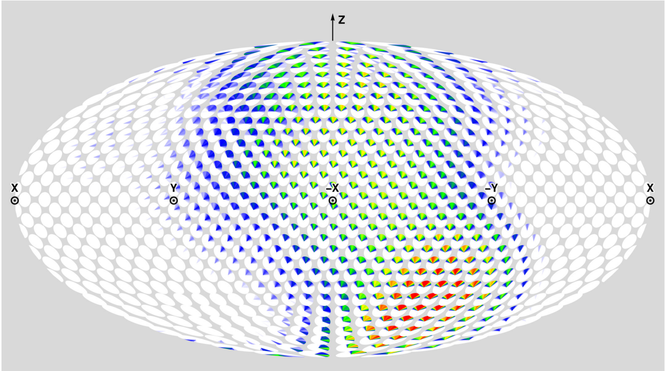

**(C)** Prediction DoS, T1200

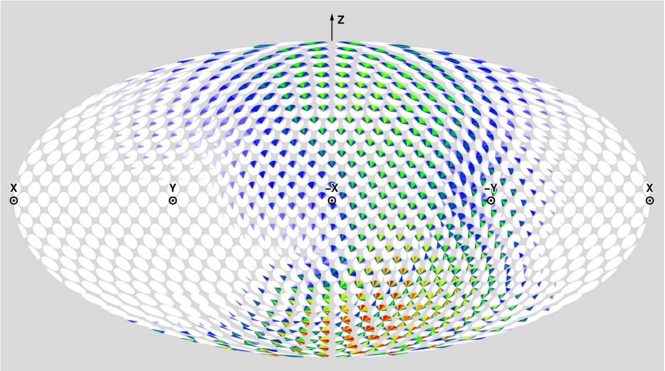

**(D)** Prediction DoS, T1300

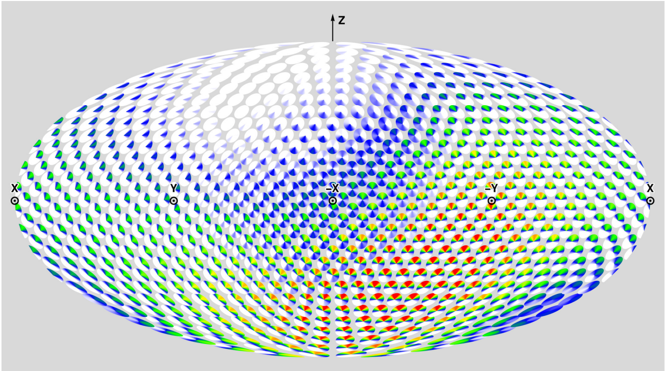

**(E)** Solution CDIO DoS ($\alpha = 0.83$)

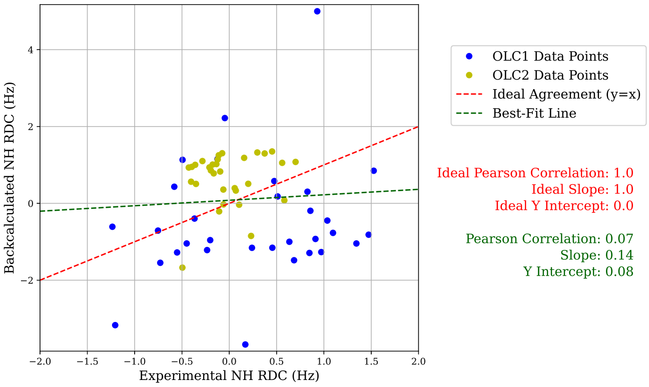


**(F)** Prediction RMB Plot, T1200

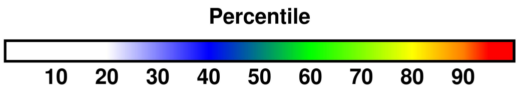


**(G)** Color percentiles for DoS

**Figure S21: Summary of analysis for predictor CSSB_FAKER/CSSB_experimental/CSSB_Human (221).** For T1200 and T1300 respectively, (A) and (B) compare back-calculated and experimental SAXS data in the form of $I(q)$ curves and residuals, with bars showing experimental standard deviation (Section 3.2). For T1200 and T1300 respectively, (C) and (D) show disk-on-sphere (DoS) visualizations of kernel density estimates of predicted ensembles (Section 2.2.3). For T1200, (E) shows the solution CDIO for $\alpha = 0.83$, which is the parameter corresponding to the smallest free energy difference with the predicted ensemble (Section 2.2.3). For T1200, (F) compares back-calculated and experimental NMR RDC data (Section 2.2.2). (G) shows the percentile color legend for (C), (D), and (E).

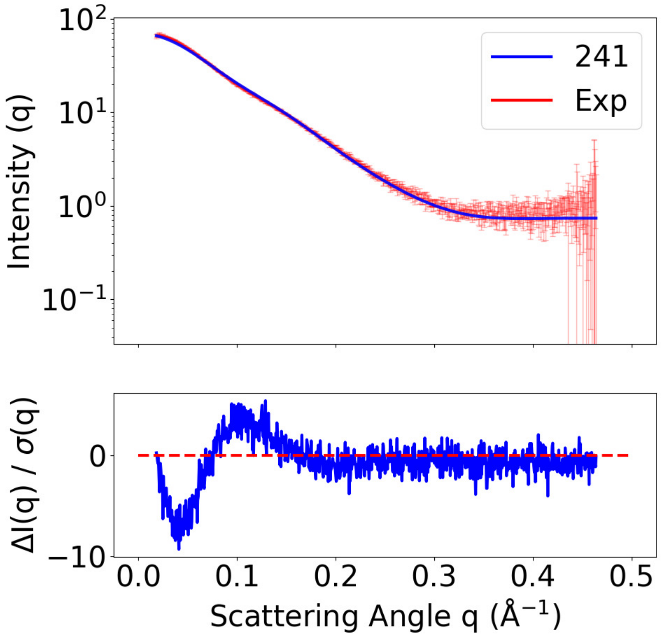


**(A)** SAXS curves, T1200

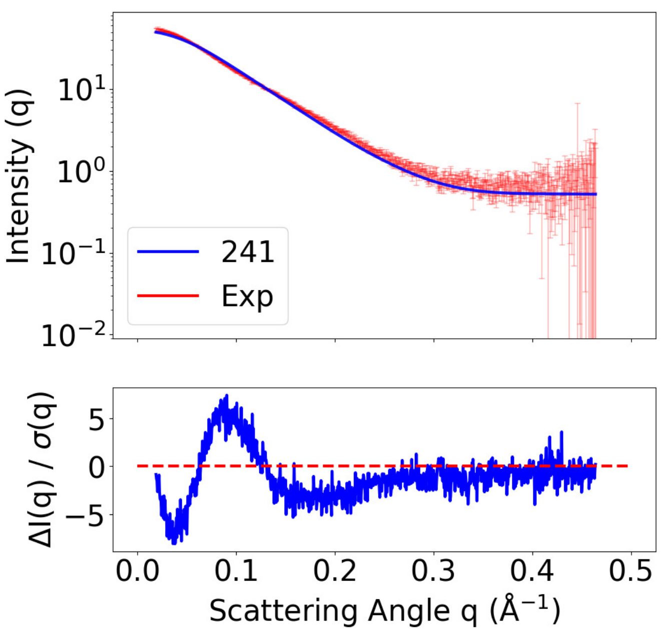


**(B)** SAXS curves, T1300

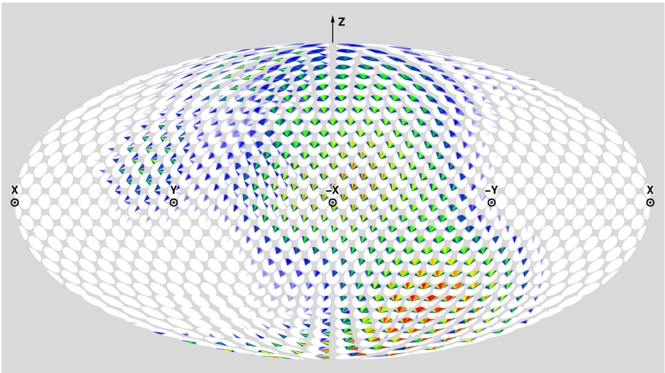

**(C)** Prediction DoS, T1200

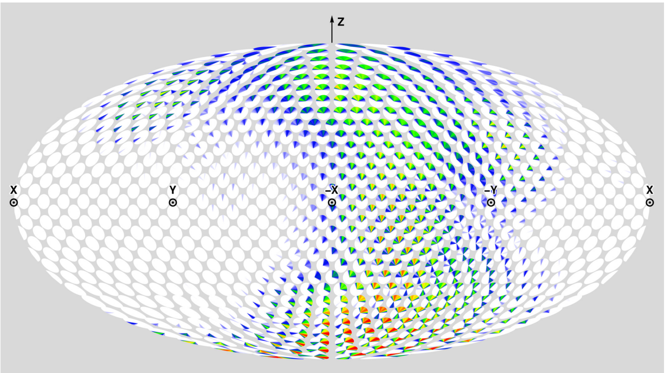

**(D)** Prediction DoS, T1300

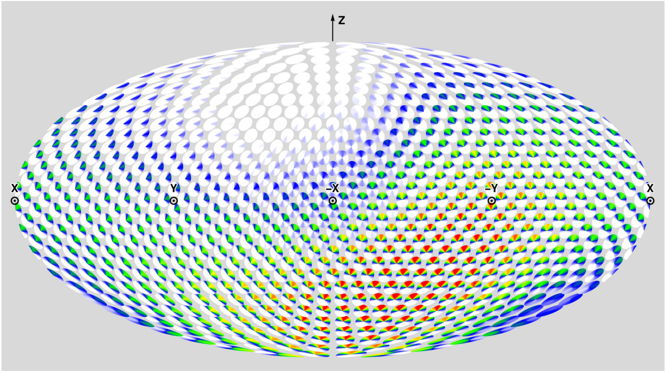

**(E)** Solution CDIO DoS ($\alpha = 0.89$)

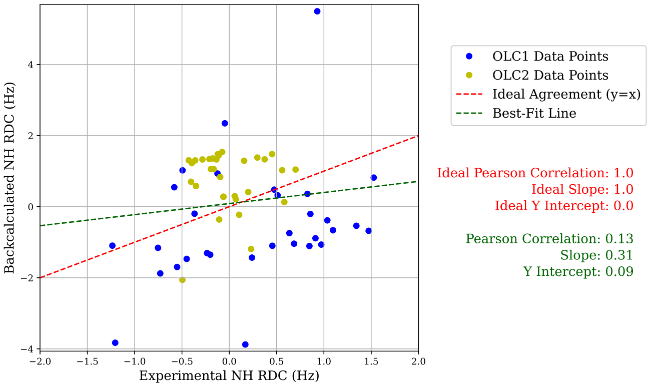


**(F)** Prediction RMB Plot, T1200

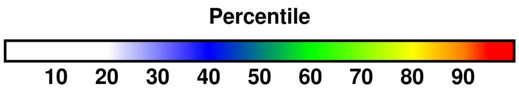


**(G)** Color percentiles for DoS

**Figure S22: Summary of analysis for predictor elofsson (241).** For T1200 and T1300 respectively, (A) and (B) compare back-calculated and experimental SAXS data in the form of $I(q)$ curves and residuals, with bars showing experimental standard deviation (Section 3.2). For T1200 and T1300 respectively, (C) and (D) show disk-on-sphere (DoS) visualizations of kernel density estimates of predicted ensembles (Section 2.2.3). For T1200, (E) shows the solution CDIO for $\alpha = 0.89$, which is the parameter corresponding to the smallest free energy difference with the predicted ensemble (Section 2.2.3). For T1200, (F) compares back-calculated and experimental NMR RDC data (Section 2.2.2). (G) shows the percentile color legend for (C), (D), and (E).

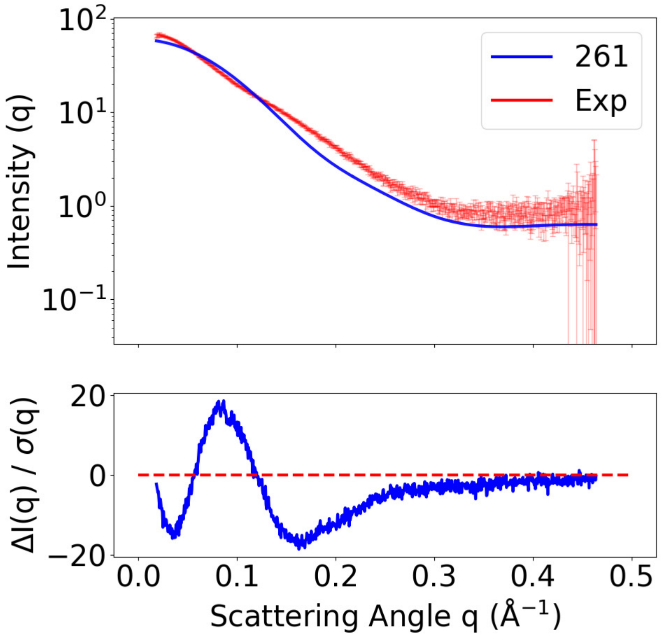


**(A)** SAXS curves, T1200

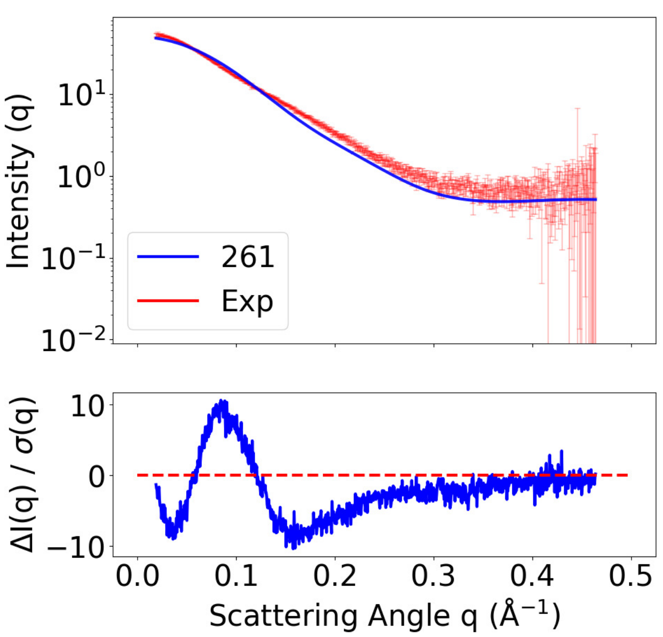


**(B)** SAXS curves, T1300

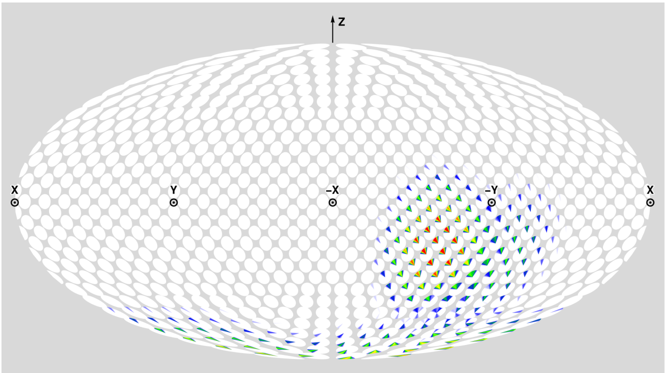


**(C)** Prediction DoS, T1200

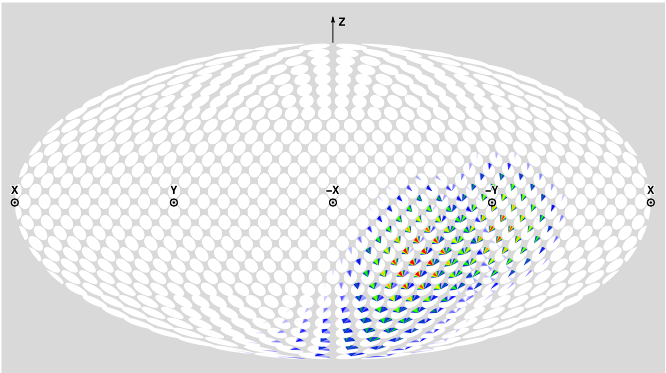


**(D)** Prediction DoS, T1300

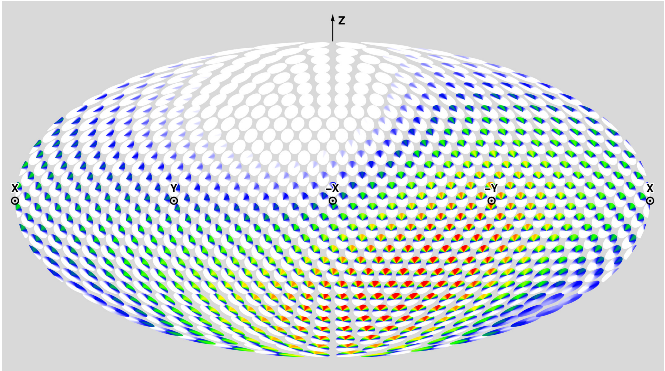


**(E)** Solution CDIO DoS ($\alpha = 1.00$)

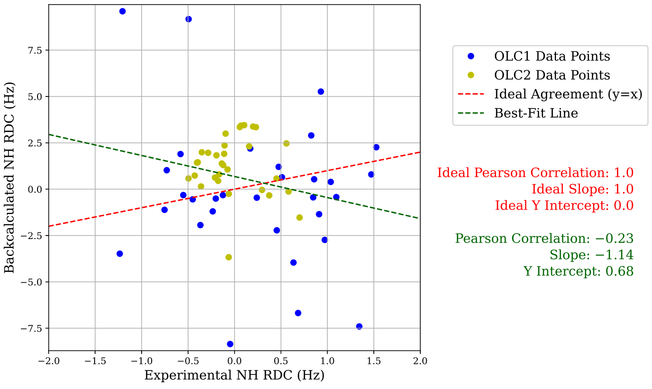


**(F)** Prediction RMB Plot, T1200

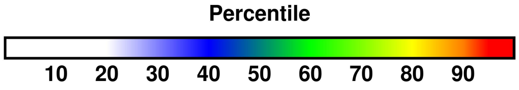


**(G)** Color percentiles for DoS

**Figure S23: Summary of analysis for predictor UNRES (261).** For T1200 and T1300 respectively, (A) and (B) compare back-calculated and experimental SAXS data in the form of $I(q)$ curves and residuals, with bars showing experimental standard deviation (Section 3.2). For T1200 and T1300 respectively, (C) and (D) show disk-on-sphere (DoS) visualizations of kernel density estimates of predicted ensembles (Section 2.2.3). For T1200, (E) shows the solution CDIO for $\alpha = 1.00$, which is the parameter corresponding to the smallest free energy difference with the predicted ensemble (Section 2.2.3). For T1200, (F) compares back-calculated and experimental NMR RDC data (Section 2.2.2). (G) shows the percentile color legend for (C), (D), and (E).

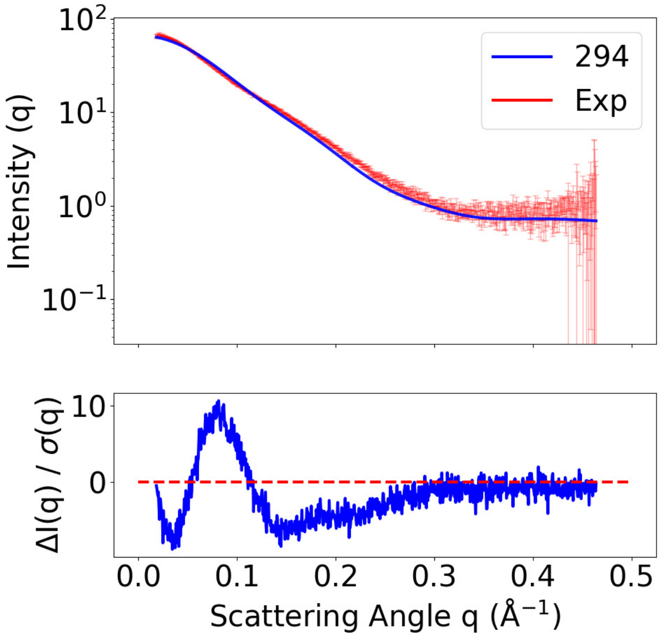


**(A)** SAXS curves, T1200

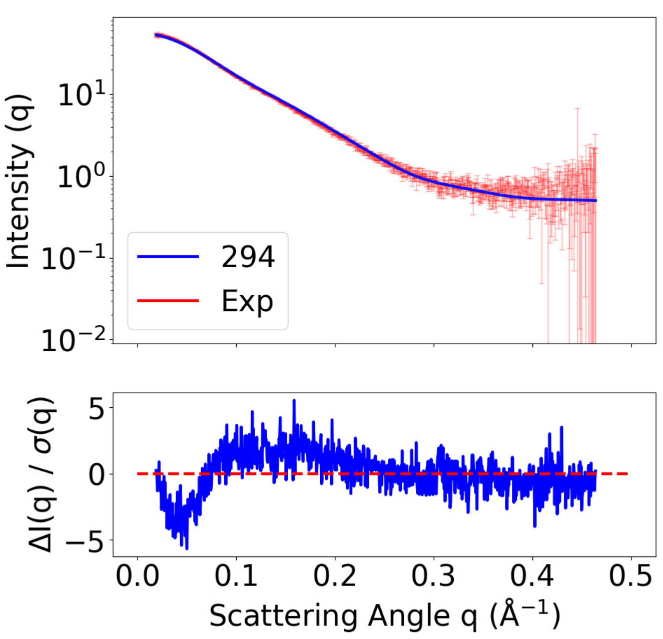


**(B)** SAXS curves, T1300

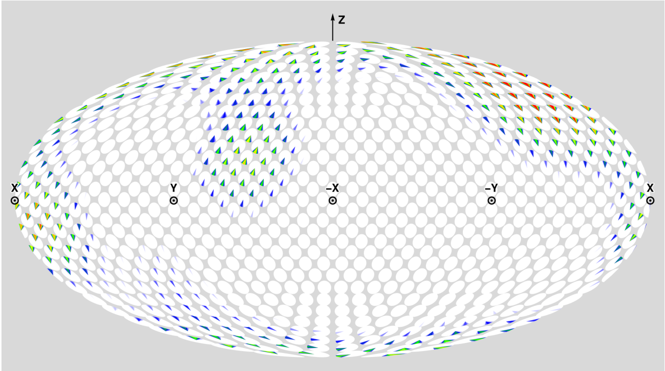


**(C)** Prediction DoS, T1200

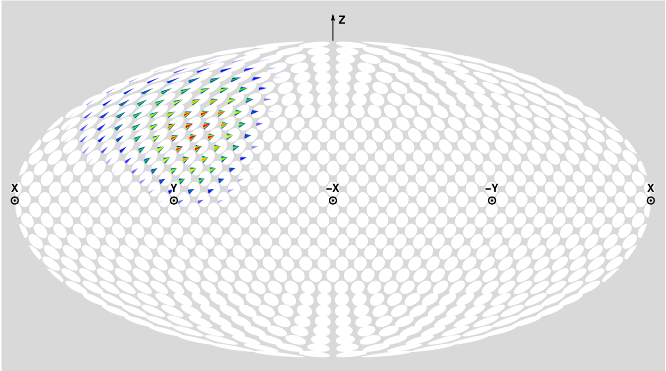


**(D)** Prediction DoS, T1300

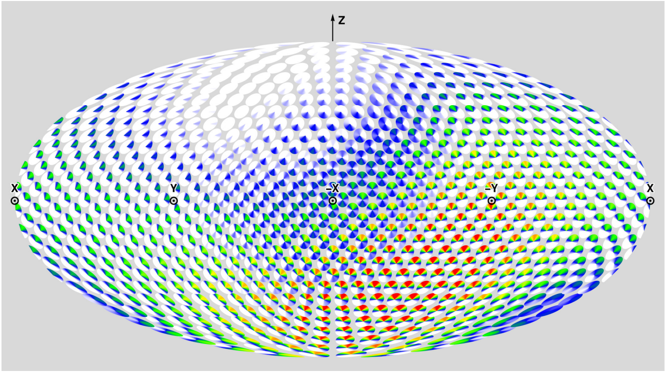


**(E)** Solution CDIO DoS ($\alpha = 0.82$)

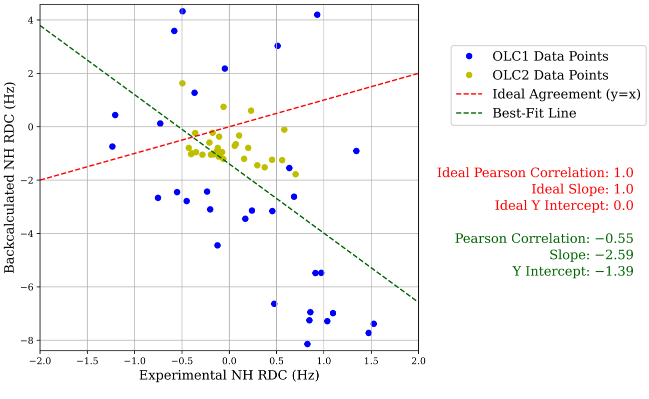


**(F)** Prediction RMB Plot, T1200

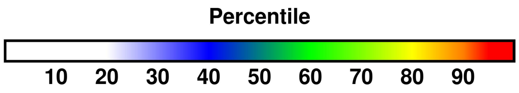


**(G)** Color percentiles for DoS

**Figure S24: Summary of analysis for predictor KiharaLab (294)**. For T1200 and T1300 respectively, (A) and (B) compare back-calculated and experimental SAXS data in the form of $I(q)$ curves and residuals, with bars showing experimental standard deviation (Section 3.2). For T1200 and T1300 respectively, (C) and (D) show disk-on-sphere (DoS) visualizations of kernel density estimates of predicted ensembles (Section 2.2.3). For T1200, (E) shows the solution CDIO for $\alpha = 0.82$, which is the parameter corresponding to the smallest free energy difference with the predicted ensemble (Section 2.2.3). For T1200, (F) compares back-calculated and experimental NMR RDC data (Section 2.2.2). (G) shows the percentile color legend for (C), (D), and (E).

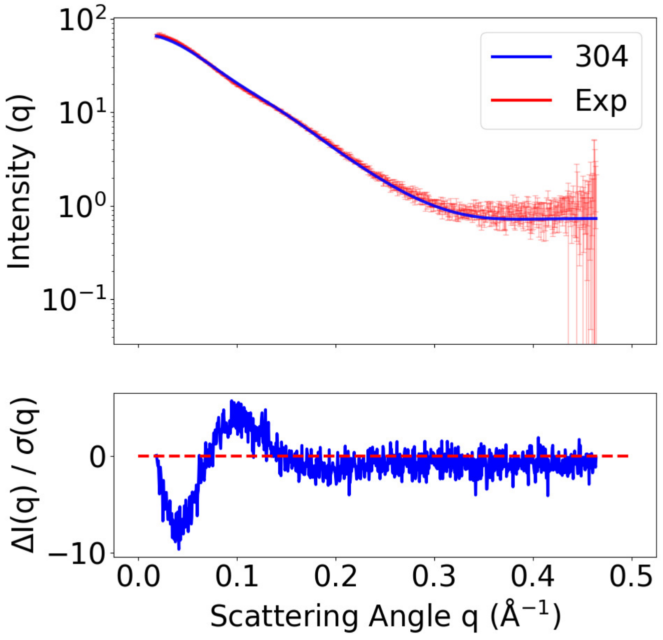


**(A)** SAXS curves, T1200

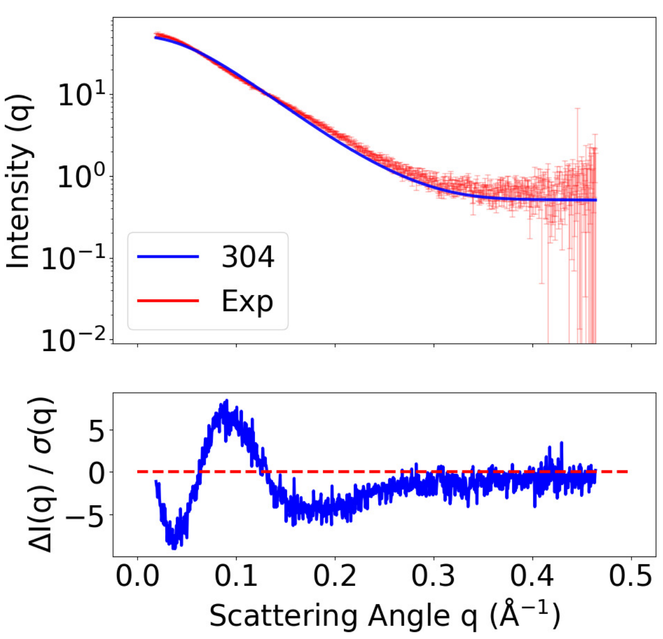


**(B)** SAXS curves, T1300

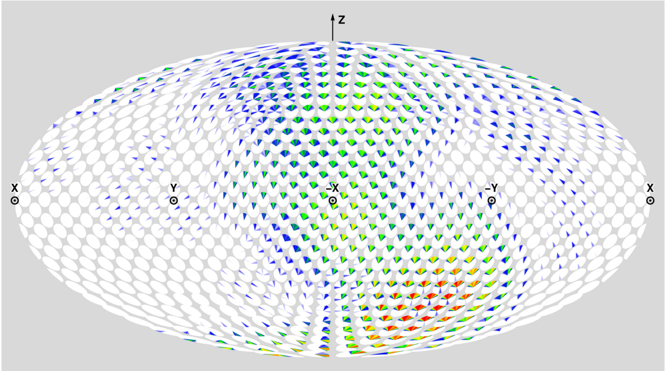


**(C)** Prediction DoS, T1200

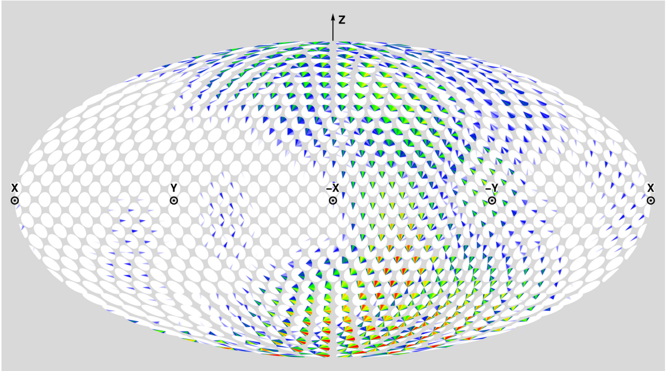


**(D)** Prediction DoS, T1300

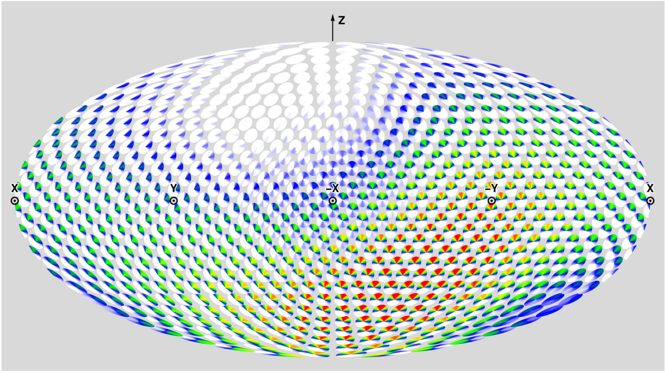


**(E)** Solution CDIO DoS ($\alpha = 0.88$)

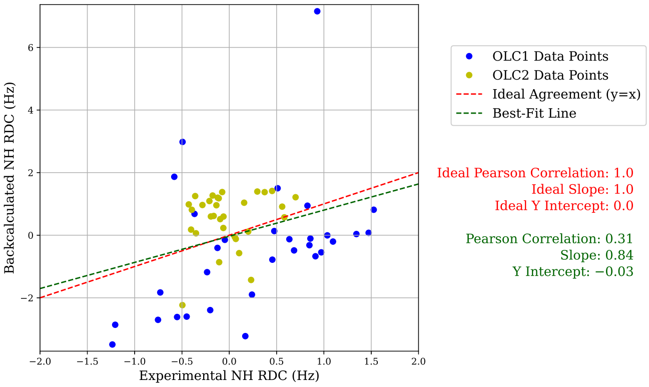


**(F)** Prediction RMB Plot, T1200

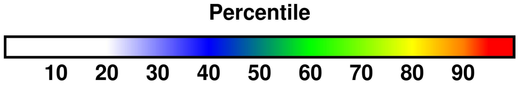


**(G)** Color percentiles for DoS

**Figure S25: Summary of analysis for predictor AF3-server (304).** For T1200 and T1300 respectively, (A) and (B) compare back-calculated and experimental SAXS data in the form of $I(q)$ curves and residuals, with bars showing experimental standard deviation (Section 3.2). For T1200 and T1300 respectively, (C) and (D) show disk-on-sphere (DoS) visualizations of kernel density estimates of predicted ensembles (Section 2.2.3). For T1200, (E) shows the solution CDIO for $\alpha = 0.88$, which is the parameter corresponding to the smallest free energy difference with the predicted ensemble (Section 2.2.3). For T1200, (F) compares back-calculated and experimental NMR RDC data (Section 2.2.2). (G) shows the percentile color legend for (C), (D), and (E).

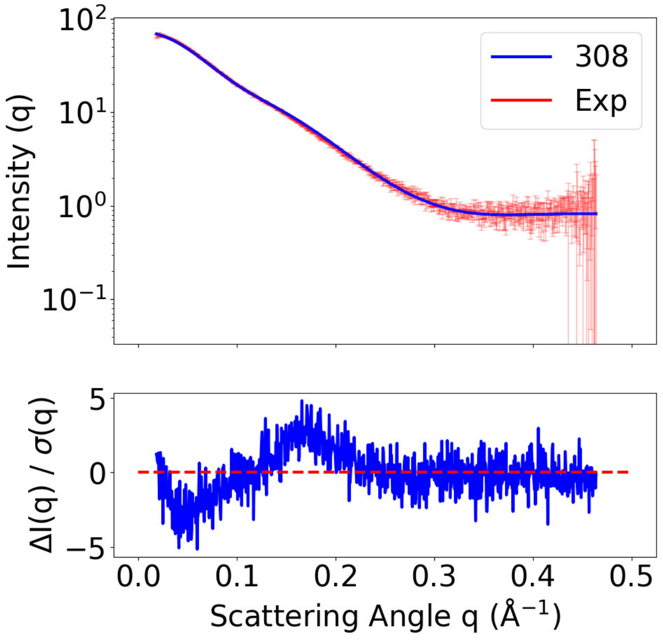


**(A)** SAXS curves, T1200

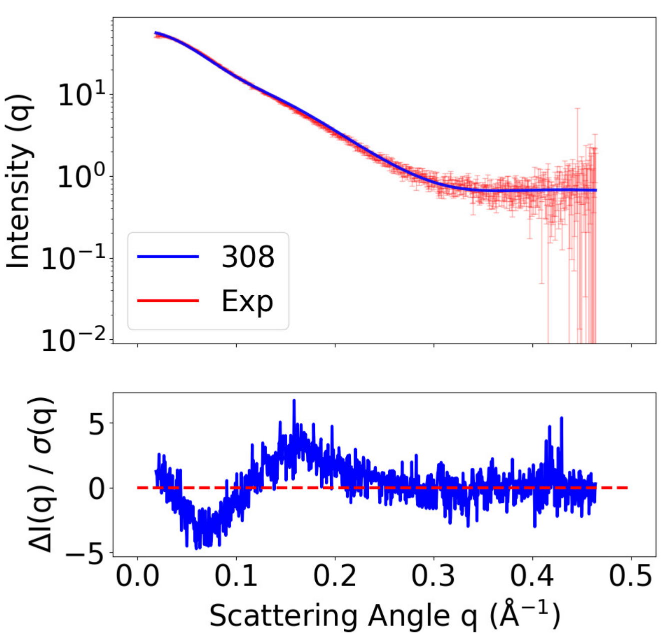


**(B)** SAXS curves, T1300

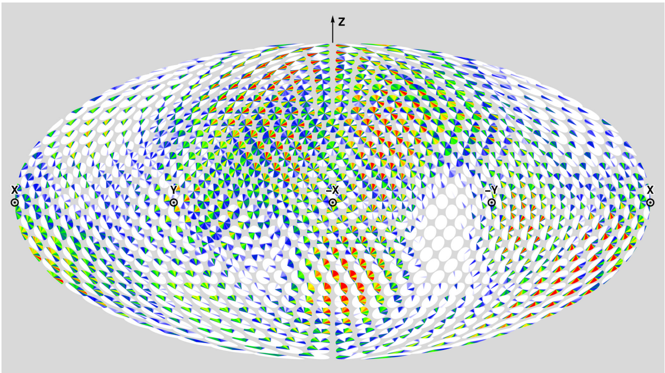

**(C)** Prediction DoS, T1200

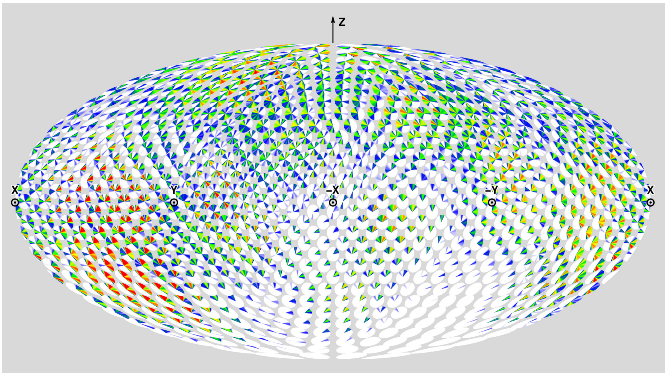

**(D)** Prediction DoS, T1300

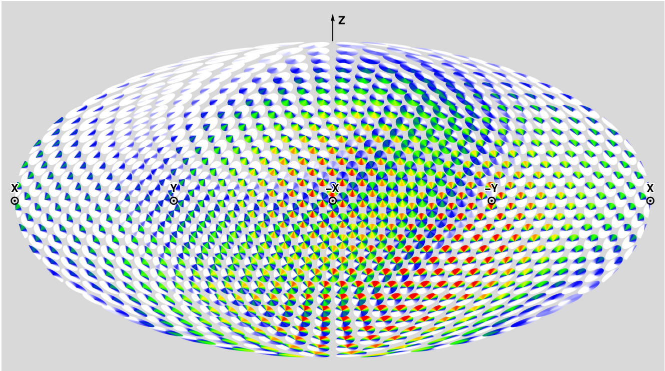

**(E)** Solution CDIO DoS ($\alpha = 0.60$)

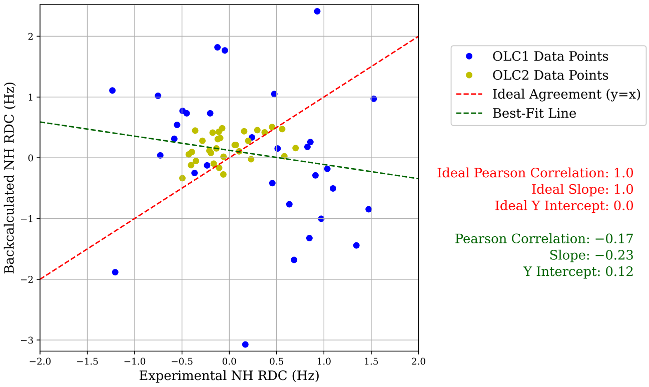


**(F)** Prediction RMB Plot, T1200

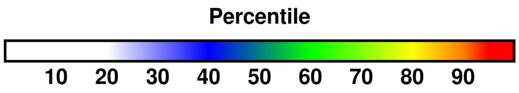


**(G)** Color percentiles for DoS

**Figure S26: Summary of analysis for predictor MoMAteam1 (308).** For T1200 and T1300 respectively, (A) and (B) compare back-calculated and experimental SAXS data in the form of $I(q)$ curves and residuals, with bars showing experimental standard deviation (Section 3.2). For T1200 and T1300 respectively, (C) and (D) show disk-on-sphere (DoS) visualizations of kernel density estimates of predicted ensembles (Section 2.2.3). For T1200, (E) shows the solution CDIO for $\alpha = 0.60$, which is the parameter corresponding to the smallest free energy difference with the predicted ensemble (Section 2.2.3). For T1200, (F) compares back-calculated and experimental NMR RDC data (Section 2.2.2). (G) shows the percentile color legend for (C), (D), and (E).

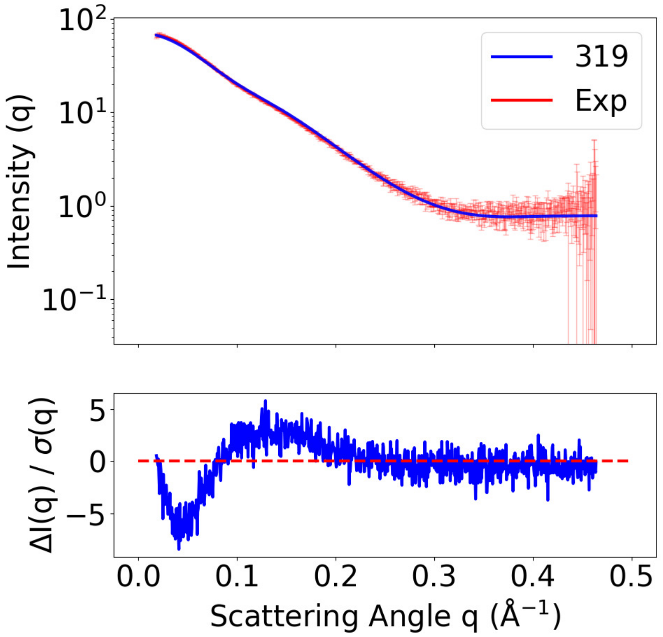


**(A)** SAXS curves, T1200

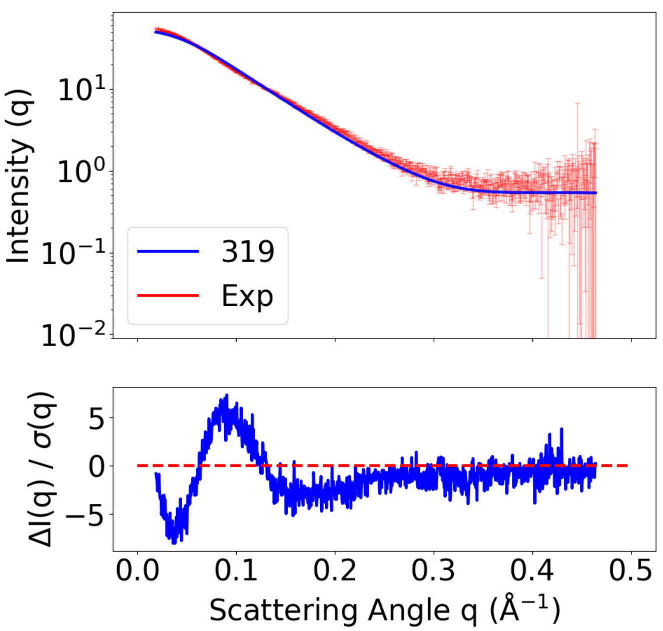


**(B)** SAXS curves, T1300

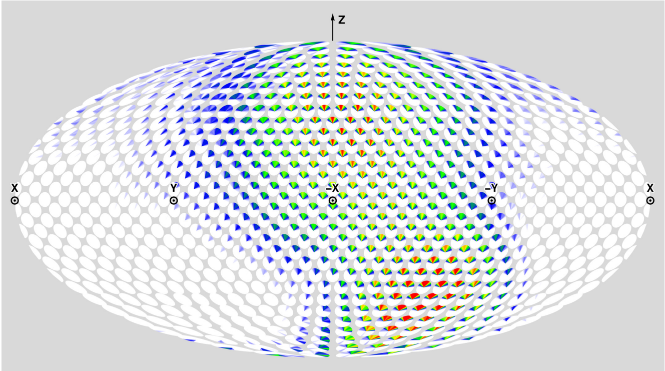


**(C)** Prediction DoS, T1200

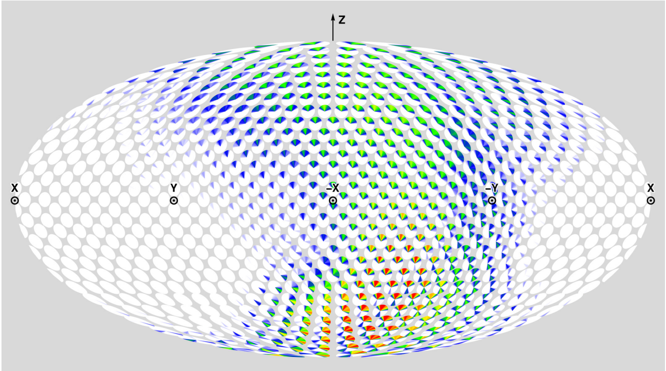


**(D)** Prediction DoS, T1300

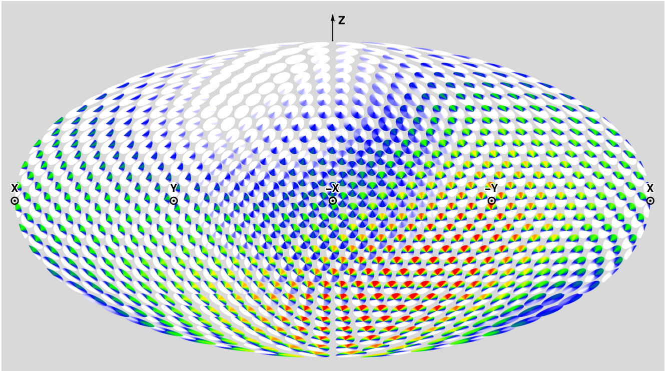


**(E)** Solution CDIO DoS ($\alpha = 0.82$)

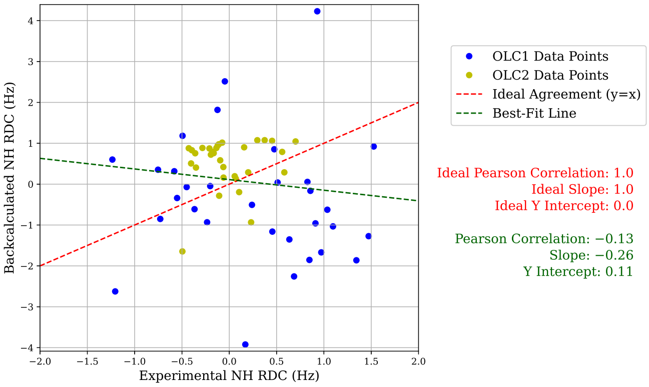


**(F)** Prediction RMB Plot, T1200

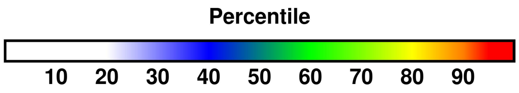


**(G)** Color percentiles for DoS

**Figure S27: Summary of analysis for predictor MULTICOM_LLM (319).** For T1200 and T1300 respectively, (A) and (B) compare back-calculated and experimental SAXS data in the form of $I(q)$ curves and residuals, with bars showing experimental standard deviation (Section 3.2). For T1200 and T1300 respectively, (C) and (D) show disk-on-sphere (DoS) visualizations of kernel density estimates of predicted ensembles (Section 2.2.3). For T1200, (E) shows the solution CDIO for $\alpha = 0.82$, which is the parameter corresponding to the smallest free energy difference with the predicted ensemble (Section 2.2.3). For T1200, (F) compares back-calculated and experimental NMR RDC data (Section 2.2.2). (G) shows the percentile color legend for (C), (D), and (E).

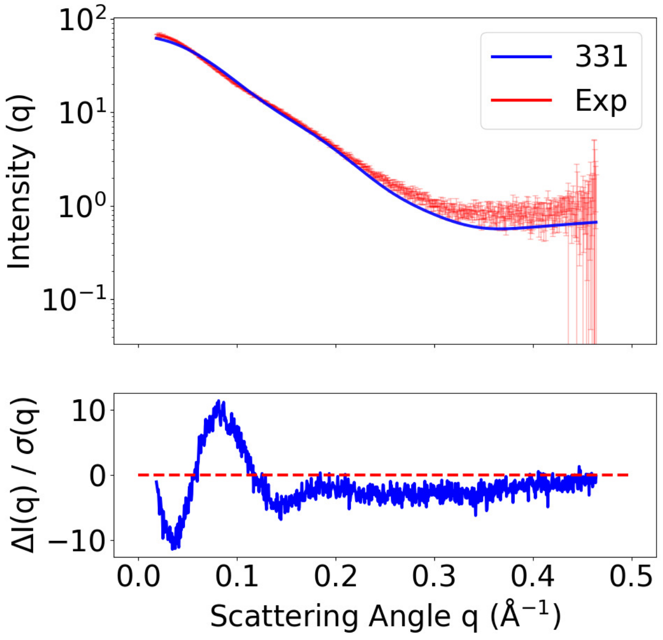


**(A)** SAXS curves, T1200

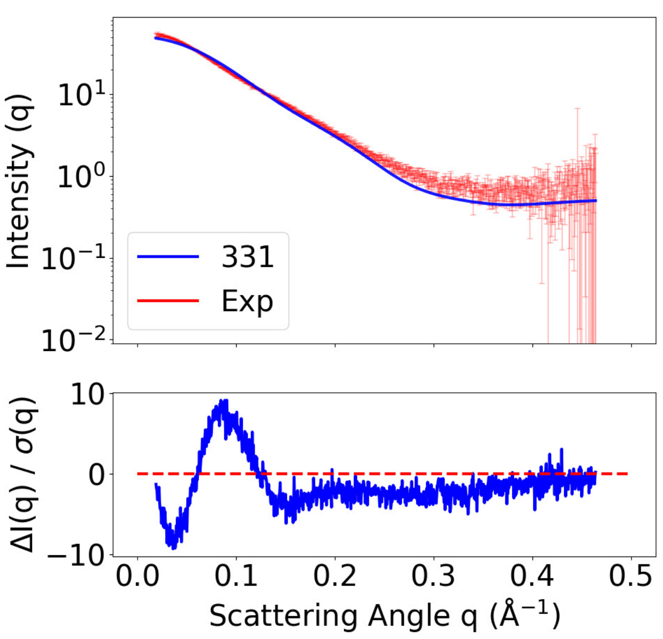


**(B)** SAXS curves, T1300

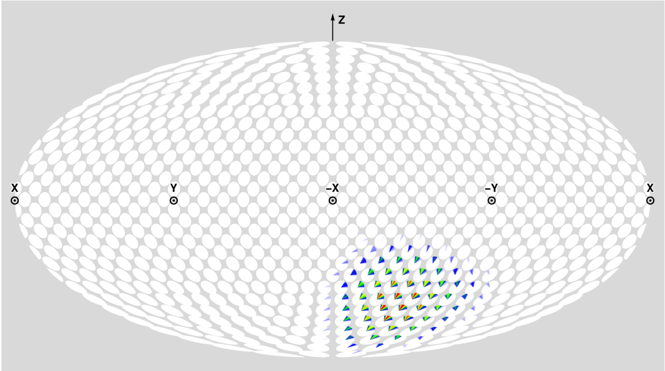


**(C)** Prediction DoS, T1200

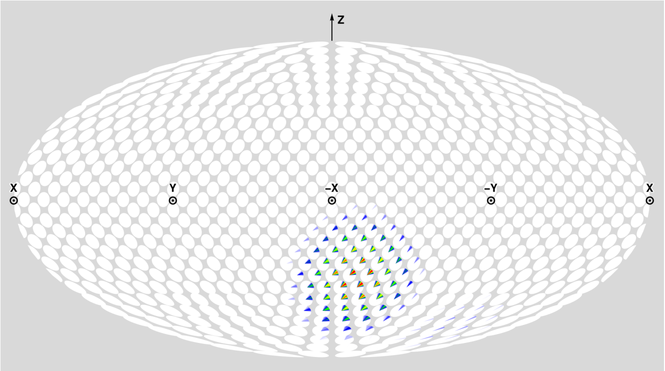


**(D)** Prediction DoS, T1300

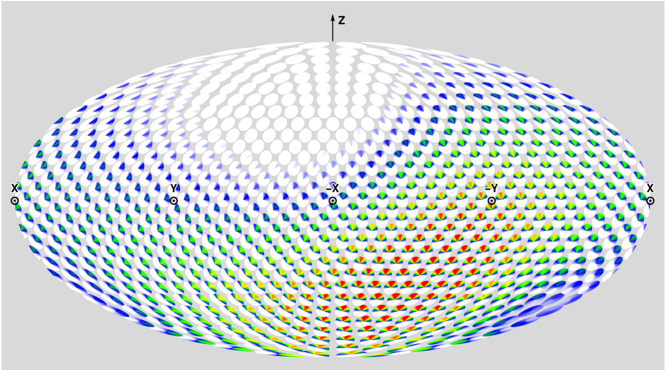


**(E)** Solution CDIO DoS ($\alpha = 1.00$)

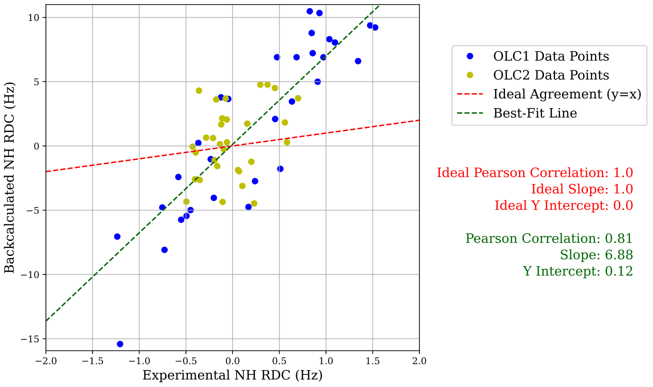


**(F)** Prediction RMB Plot, T1200

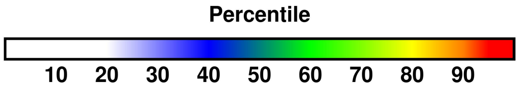


**(G)** Color percentiles for DoS

**Figure S28: Summary of analysis for predictor MULTICOM_AI (331).** For T1200 and T1300 respectively, (A) and (B) compare back-calculated and experimental SAXS data in the form of $I(q)$ curves and residuals, with bars showing experimental standard deviation (Section 3.2). For T1200 and T1300 respectively, (C) and (D) show disk-on-sphere (DoS) visualizations of kernel density estimates of predicted ensembles (Section 2.2.3). For T1200, (E) shows the solution CDIO for $\alpha = 1.00$, which is the parameter corresponding to the smallest free energy difference with the predicted ensemble (Section 2.2.3). For T1200, (F) compares back-calculated and experimental NMR RDC data (Section 2.2.2). (G) shows the percentile color legend for (C), (D), and (E).

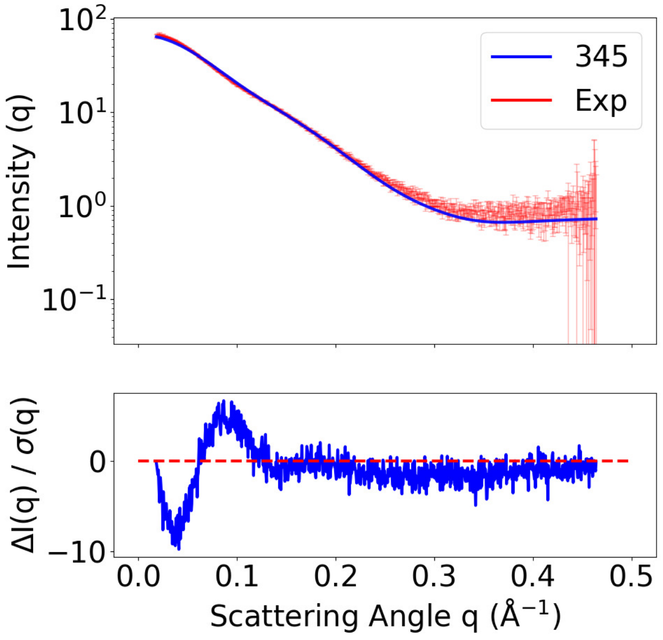


**(A)** SAXS curves, T1200

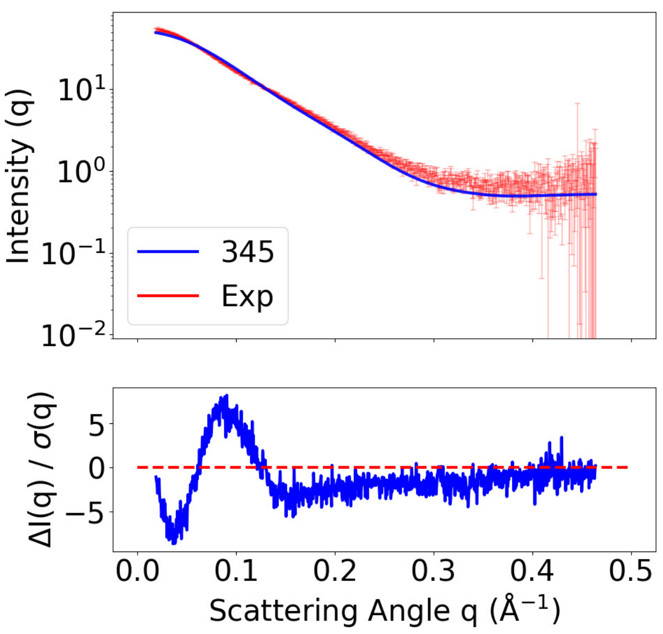


**(B)** SAXS curves, T1300

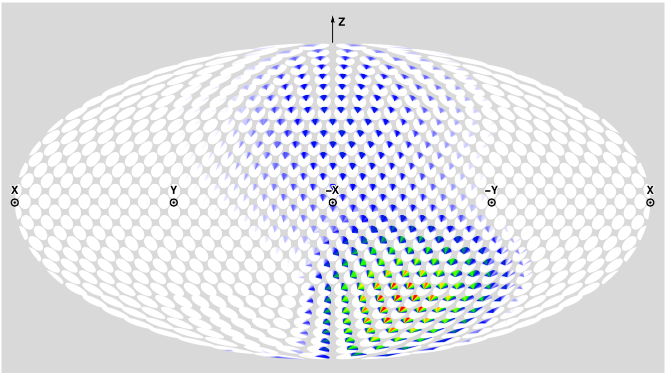

**(C)** Prediction DoS, T1200

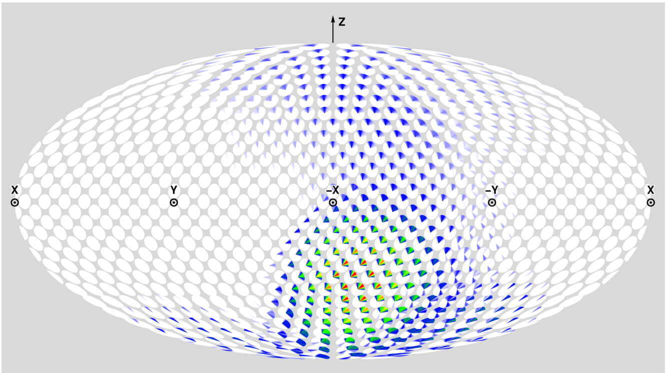

**(D)** Prediction DoS, T1300

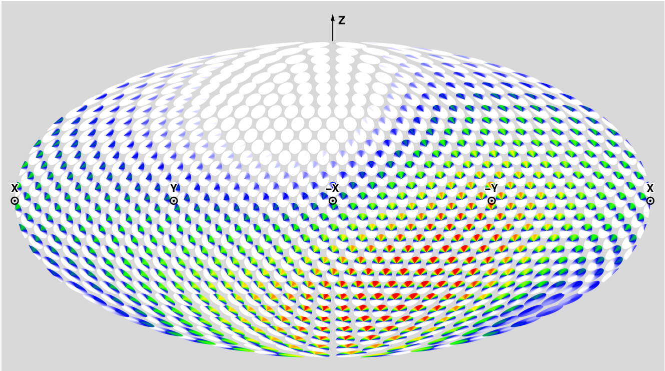

**(E)** Solution CDIO DoS ($\alpha = 0.98$)

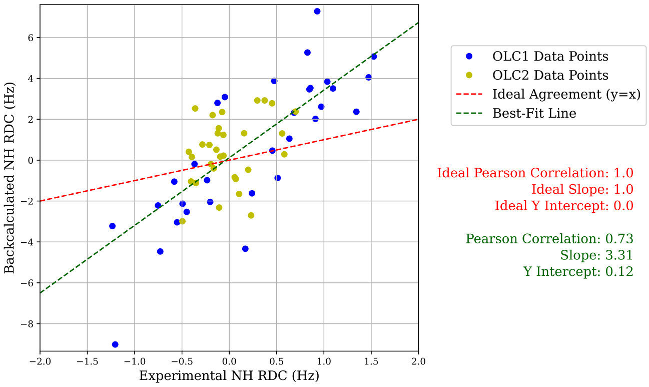


**(F)** Prediction RMB Plot, T1200

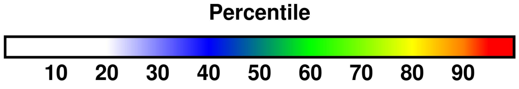


**(G)** Color percentiles for DoS

**Figure S29: Summary of analysis for predictor MULTICOM_human (345)**. For T1200 and T1300 respectively, (A) and (B) compare back-calculated and experimental SAXS data in the form of $I(q)$ curves and residuals, with bars showing experimental standard deviation (Section 3.2). For T1200 and T1300 respectively, (C) and (D) show disk-on-sphere (DoS) visualizations of kernel density estimates of predicted ensembles (Section 2.2.3). For T1200, (E) shows the solution CDIO for $\alpha = 0.98$, which is the parameter corresponding to the smallest free energy difference with the predicted ensemble (Section 2.2.3). For T1200, (F) compares back-calculated and experimental NMR RDC data (Section 2.2.2). (G) shows the percentile color legend for (C), (D), and (E).

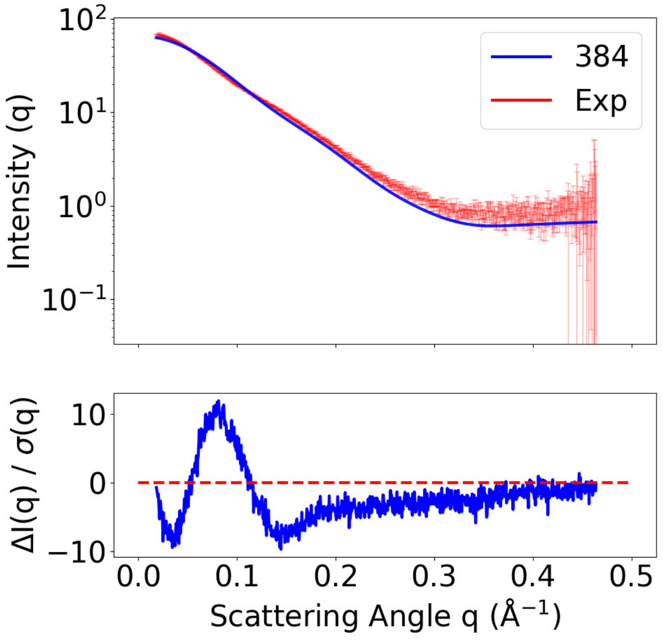


**(A)** SAXS curves, T1200

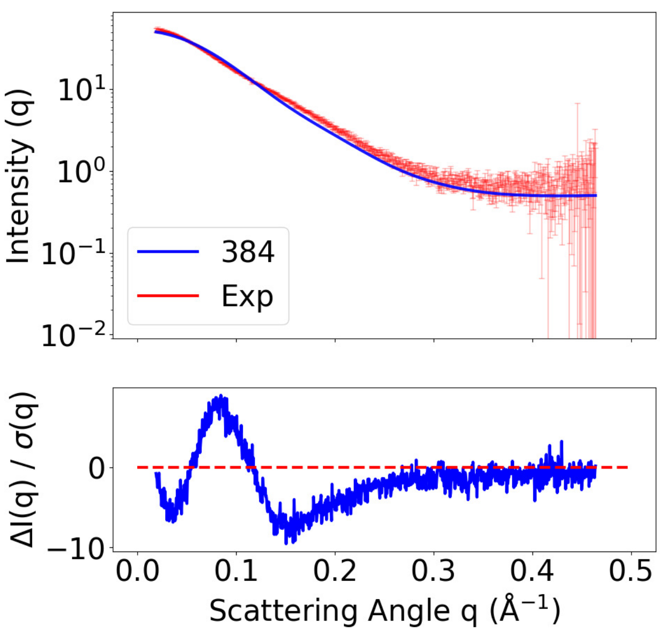


**(B)** SAXS curves, T1300

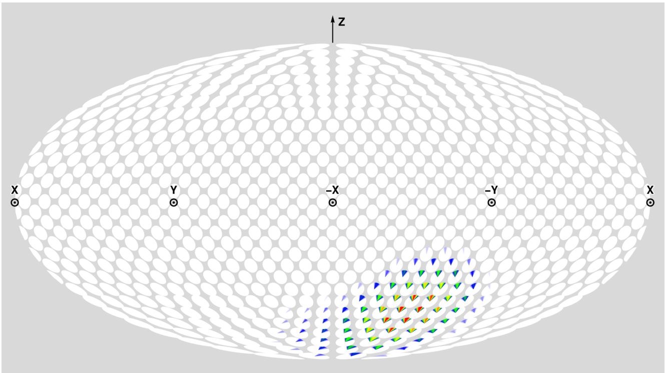


**(C)** Prediction DoS, T1200

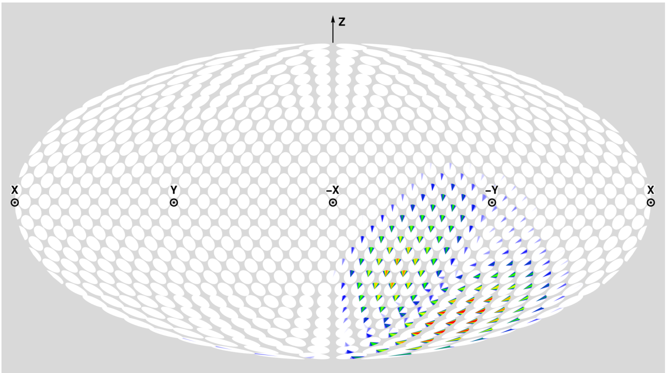


**(D)** Prediction DoS, T1300

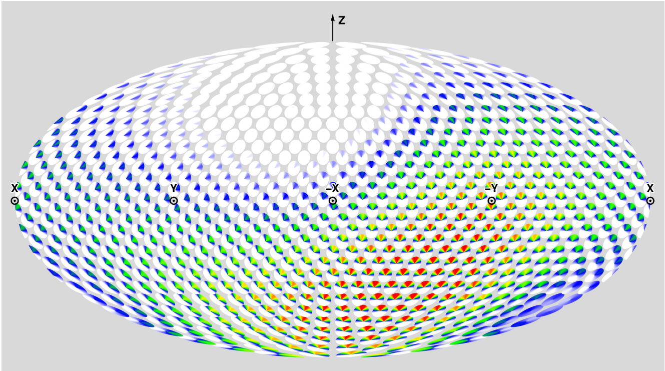


**(E)** Solution CDIO DoS ($\alpha = 1.00$)

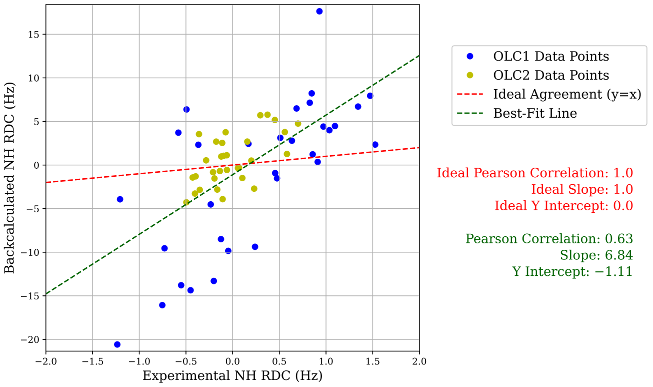


**(F)** Prediction RMB Plot, T1200

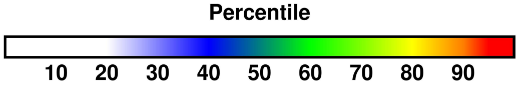


**(G)** Color percentiles for DoS

**Figure S30: Summary of analysis for predictor pert-plddt (384).** For T1200 and T1300 respectively, (A) and (B) compare back-calculated and experimental SAXS data in the form of $I(q)$ curves and residuals, with bars showing experimental standard deviation (Section 3.2). For T1200 and T1300 respectively, (C) and (D) show disk-on-sphere (DoS) visualizations of kernel density estimates of predicted ensembles (Section 2.2.3). For T1200, (E) shows the solution CDIO for $\alpha = 1.00$, which is the parameter corresponding to the smallest free energy difference with the predicted ensemble (Section 2.2.3). For T1200, (F) compares back-calculated and experimental NMR RDC data (Section 2.2.2). (G) shows the percentile color legend for (C), (D), and (E).

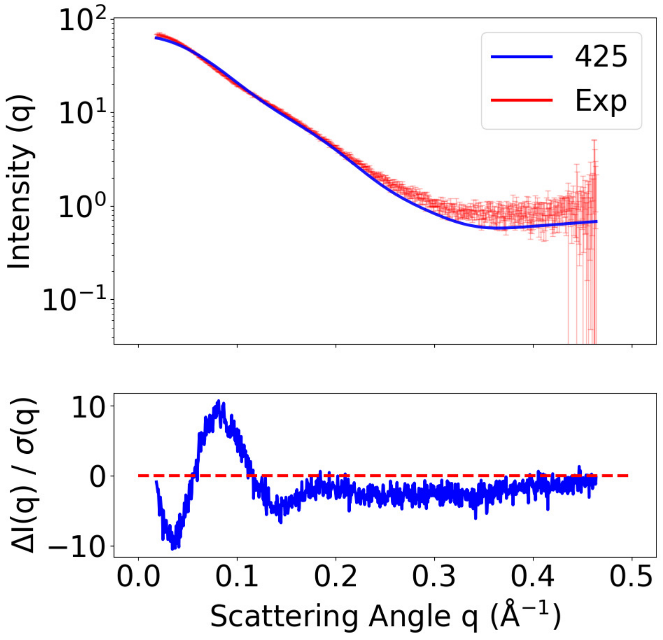


**(A)** SAXS curves, T1200

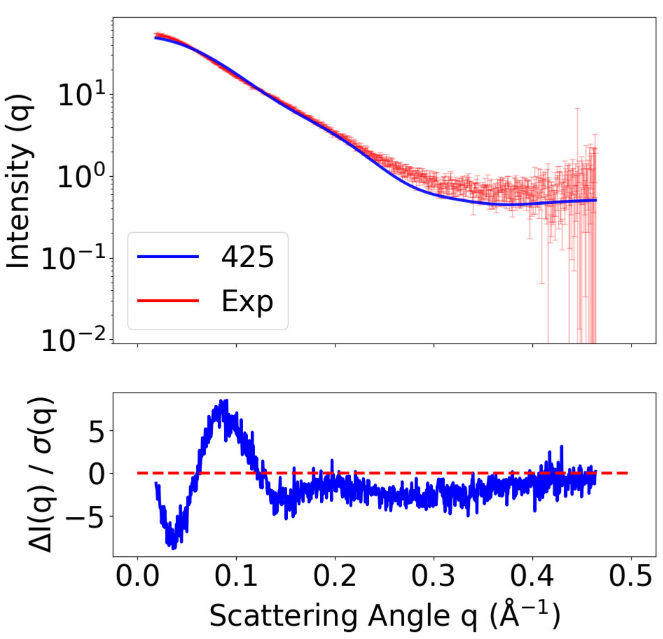


**(B)** SAXS curves, T1300

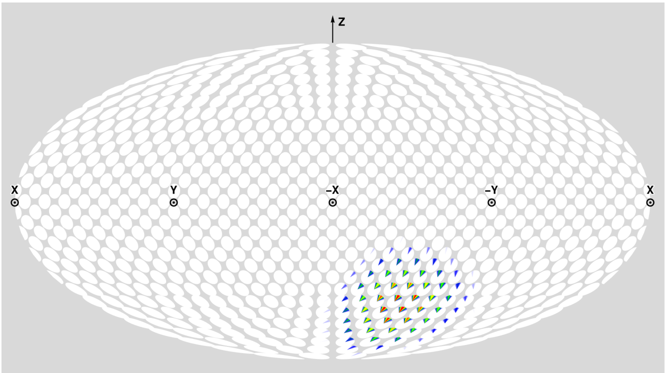


**(C)** Prediction DoS, T1200

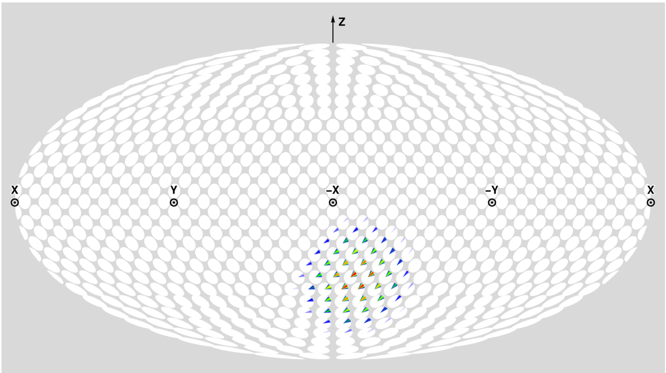


**(D)** Prediction DoS, T1300

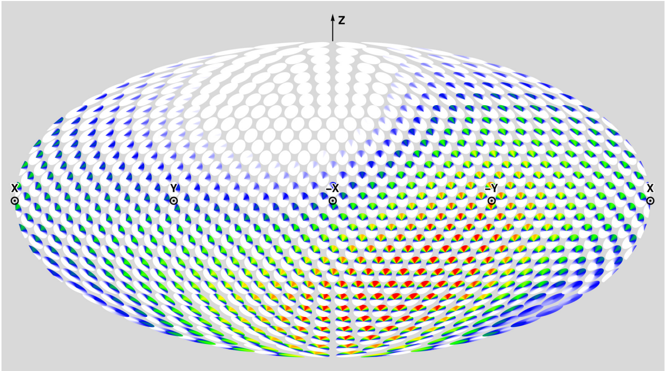


**(E)** Solution CDIO DoS ($\alpha = 1.00$)

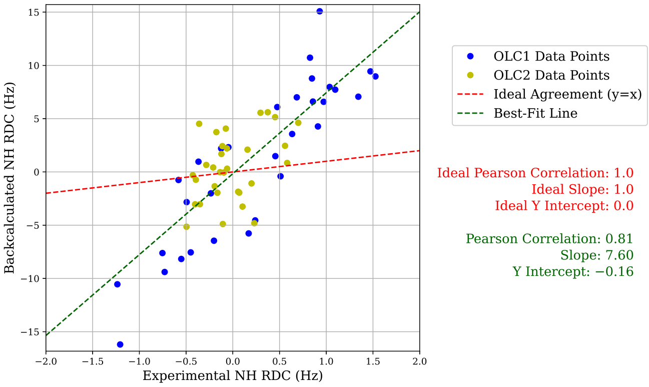


**(F)** Prediction RMB Plot, T1200

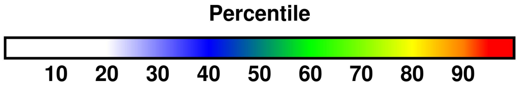


**(G)** Color percentiles for DoS

**Figure S31: Summary of analysis for predictor MULTICOM_GATE (425).** For T1200 and T1300 respectively, (A) and (B) compare back-calculated and experimental SAXS data in the form of $I(q)$ curves and residuals, with bars showing experimental standard deviation (Section 3.2). For T1200 and T1300 respectively, (C) and (D) show disk-on-sphere (DoS) visualizations of kernel density estimates of predicted ensembles (Section 2.2.3). For T1200, (E) shows the solution CDIO for $\alpha = 1.00$, which is the parameter corresponding to the smallest free energy difference with the predicted ensemble (Section 2.2.3). For T1200, (F) compares back-calculated and experimental NMR RDC data (Section 2.2.2). (G) shows the percentile color legend for (C), (D), and (E).

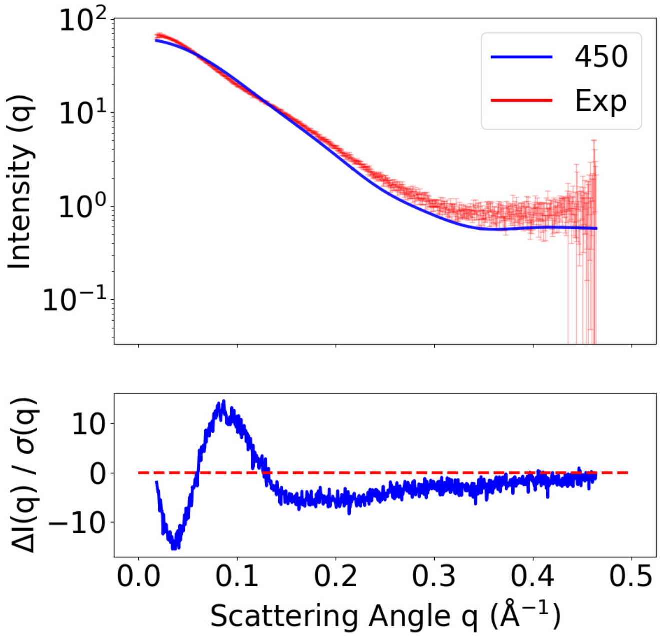


**(A)** SAXS curves, T1200

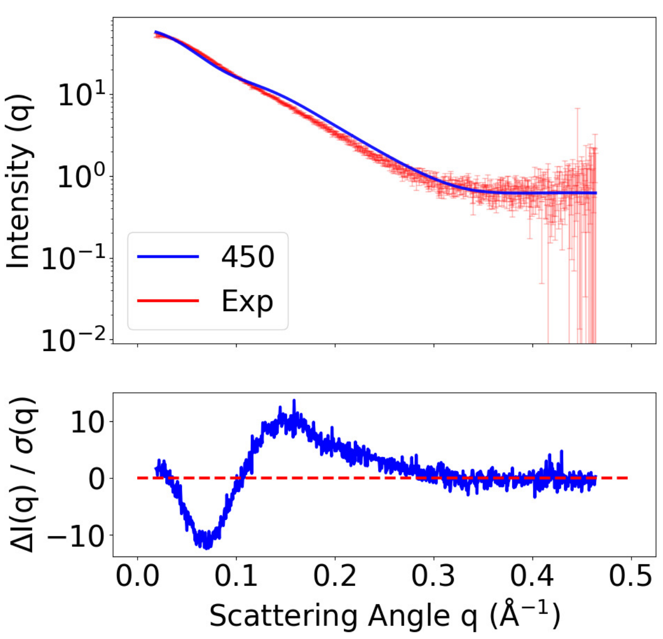


**(B)** SAXS curves, T1300

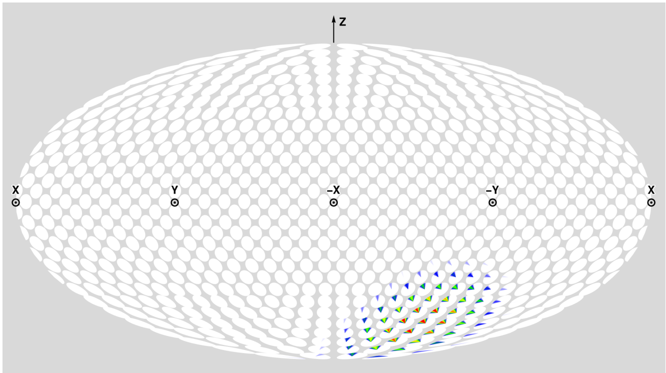


**(C)** Prediction DoS, T1200

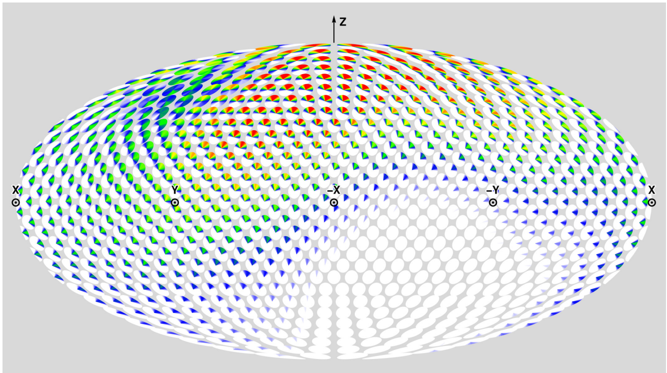


**(D)** Prediction DoS, T1300

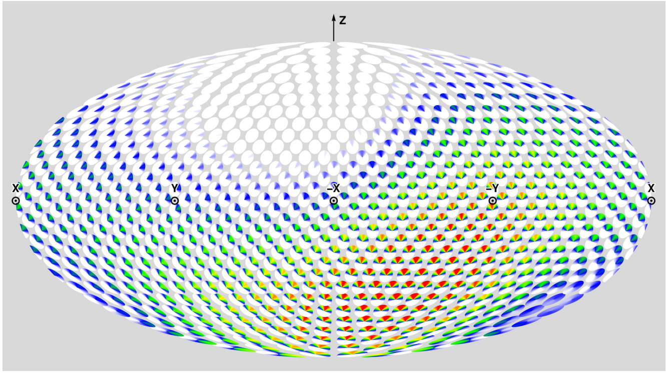


**(E)** Solution CDIO DoS ($\alpha = 1.00$)

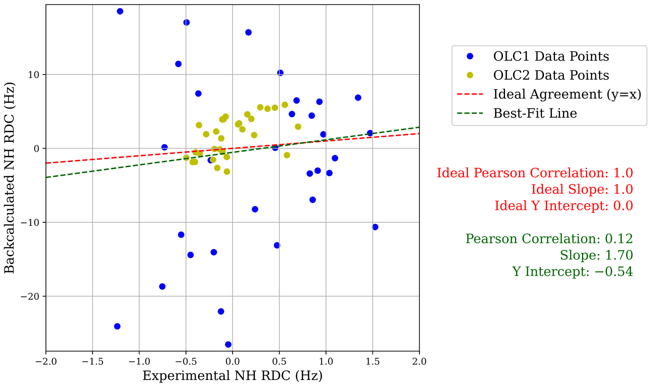


**(F)** Prediction RMB Plot, T1200

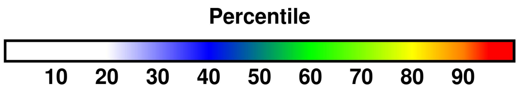


**(G)** Color percentiles for DoS

**Figure S32: Summary of analysis for predictor OpenComplex_Server (450).** For T1200 and T1300 respectively, (A) and (B) compare back-calculated and experimental SAXS data in the form of $I(q)$ curves and residuals, with bars showing experimental standard deviation (Section 3.2). For T1200 and T1300 respectively, (C) and (D) show disk-on-sphere (DoS) visualizations of kernel density estimates of predicted ensembles (Section 2.2.3). For T1200, (E) shows the solution CDIO for $\alpha = 1.00$, which is the parameter corresponding to the smallest free energy difference with the predicted ensemble (Section 2.2.3). For T1200, (F) compares back-calculated and experimental NMR RDC data (Section 2.2.2). (G) shows the percentile color legend for (C), (D), and (E).

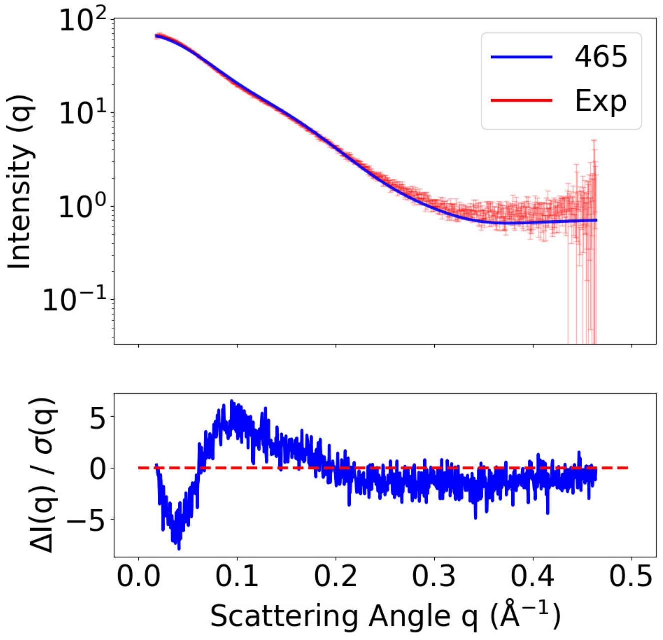


**(A)** SAXS curves, T1200

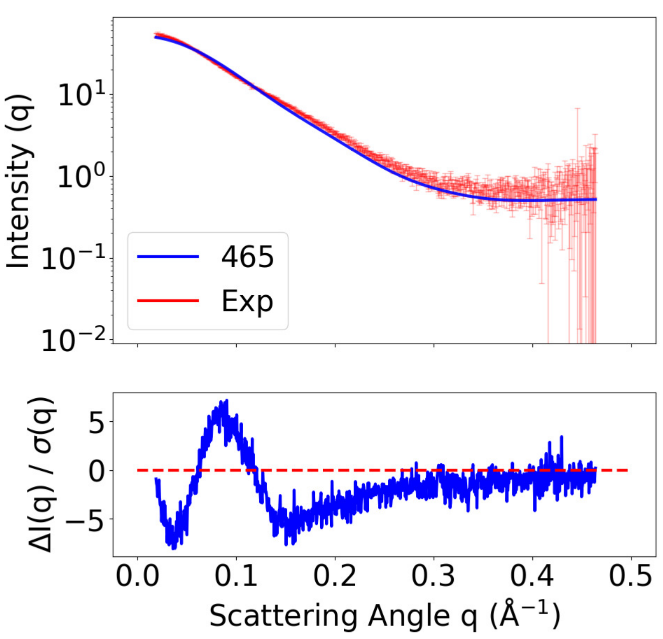


**(B)** SAXS curves, T1300

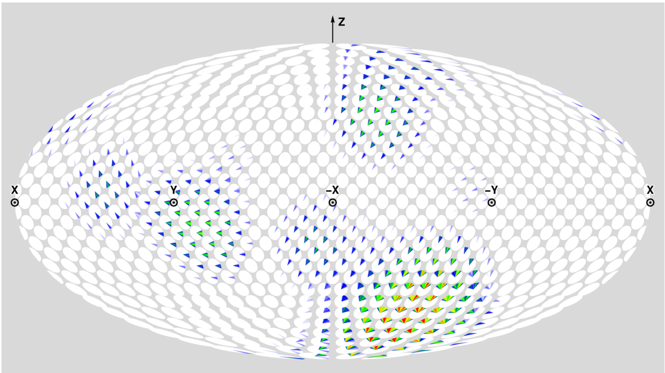


**(C)** Prediction DoS, T1200

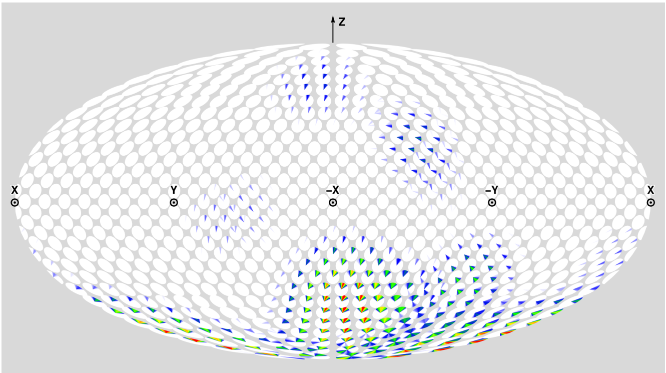


**(D)** Prediction DoS, T1300

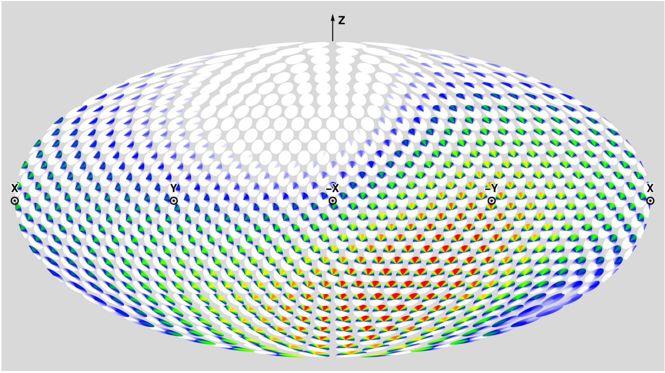


**(E)** Solution CDIO DoS ($\alpha = 1.00$)

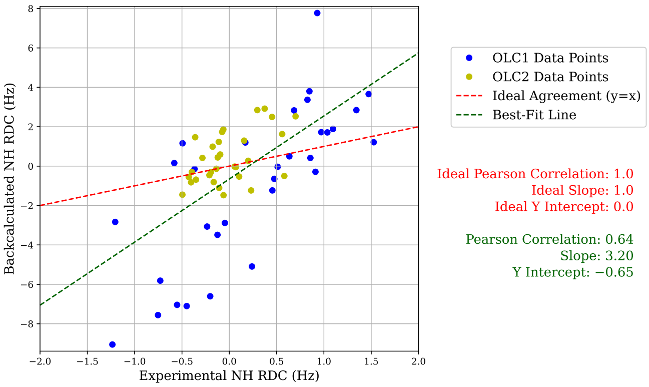


**(F)** Prediction RMB Plot, T1200

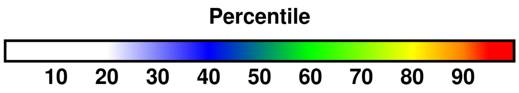


**(G)** Color percentiles for DoS

**Figure S33: Summary of analysis for predictor Wallner (465).** For T1200 and T1300 respectively, (A) and (B) compare back-calculated and experimental SAXS data in the form of $I(q)$ curves and residuals, with bars showing experimental standard deviation (Section 3.2). For T1200 and T1300 respectively, (C) and (D) show disk-on-sphere (DoS) visualizations of kernel density estimates of predicted ensembles (Section 2.2.3). For T1200, (E) shows the solution CDIO for $\alpha = 1.00$, which is the parameter corresponding to the smallest free energy difference with the predicted ensemble (Section 2.2.3). For T1200, (F) compares back-calculated and experimental NMR RDC data (Section 2.2.2). (G) shows the percentile color legend for (C), (D), and (E).

### Specifications for predicting the structure of the two-domain protein ZLBT-C

Terry Oas*[,1,2], Aulane Mpouli[2], Edward Cheng[2], and Bruce Donald*[,1,2,3]

[1]Department of Biochemistry, Duke University
[2]Department of Chemistry, Duke University
[3]Department of Computer Science, Duke University
*Contacts:

May, 2024

ZLBT-C is a mimic of two of the five nearly identical three-helix bundle domains from the N-terminal region of staphylococcal protein A (Figure 1)[1]. ZLBT is a biotech variant of the B domain of protein A with a lanthanide binding tag inserted between helices 2 and 3. The C domain is linked to ZLBT via the 6-residue wild-type B-C linker, KADNKF. The structures of the helical cores of both ZLBT[2] and C domains[3] have been determined, so the challenge outlined below is not to predict these structures, but rather to predict the range of structures that position the two domains relative to each other. The published description of this structure, based on experimental NMR residual dipolar coupling (RDC) data, is a continuous distribution of interdomain orientation (CDIO)(see Figure 2)[1]. Predictions could be made in the form of a 3D (Bingham) probability distribution over the space of the relative orientations of the two domains (SO(3)) or as an ensemble of population-weighted structures. In both cases, to compare predictions with experimental data, it is necessary to define domain-fixed Cartesian coordinate systems based on atomic coordinates of the core domain structures. For this reason, the predicted helical core structures must match the deposited structures within a minimal RMSD.

The predictions will be compared with NMR RDC data and small-angle X-ray scattering (SAXS) profiles. In this way, both the CDIO and the interdomain distance distribution of a prediction can be compared with experimental data. The requirements for such predictions are the following.

1. The backbone RMSD of residues 36-50, 70-85 (ZLBT helix 2/3 core, 2LR2, Model 1[2]) and 112-125, 129-143 (C helix 2/3 core, 4NPD, Alternates A[3]) of ZLBT-C should fall within 0.5 Å of each deposited structure (see Figure 1).
2. If the prediction takes the form of an ensemble, the population of each ensemble member must be given as a positive rational number and must sum to 1.0. The uncertainty should be provided for the population of each ensemble member. Coordinate files should be in PDB format.
3. If the prediction takes the form of a continuous distribution[1], the quaternion mean, variances, and covariances representing each Bingham distribution mode (if more than one) must be given, along with the relative probability of each mode. The Cartesian coordinate system of each domain used to define these quaternions should match the ones defined in the attached algorithm.
4. A graphical representation of the ZLBT-C CDIO has been published(Figure 2)[1], but not the quantitative properties described in 3) nor the interdomain distance distribution, so this remains a predictive challenge.
5. Unpublished RDC and SAXS data will be available for a ZLBT-C construct with a $Gly_6$ linker that replaces the wild-type linker. Predictions should be made for both the wild-type sequence and this one.
6. Predictions will be evaluated based on a comparison between the experimental and predicted CDIOs using the free energy function described in Qi, et al.[1] and the $\chi^2$ values of the predicted vs. observed SAXS profiles.

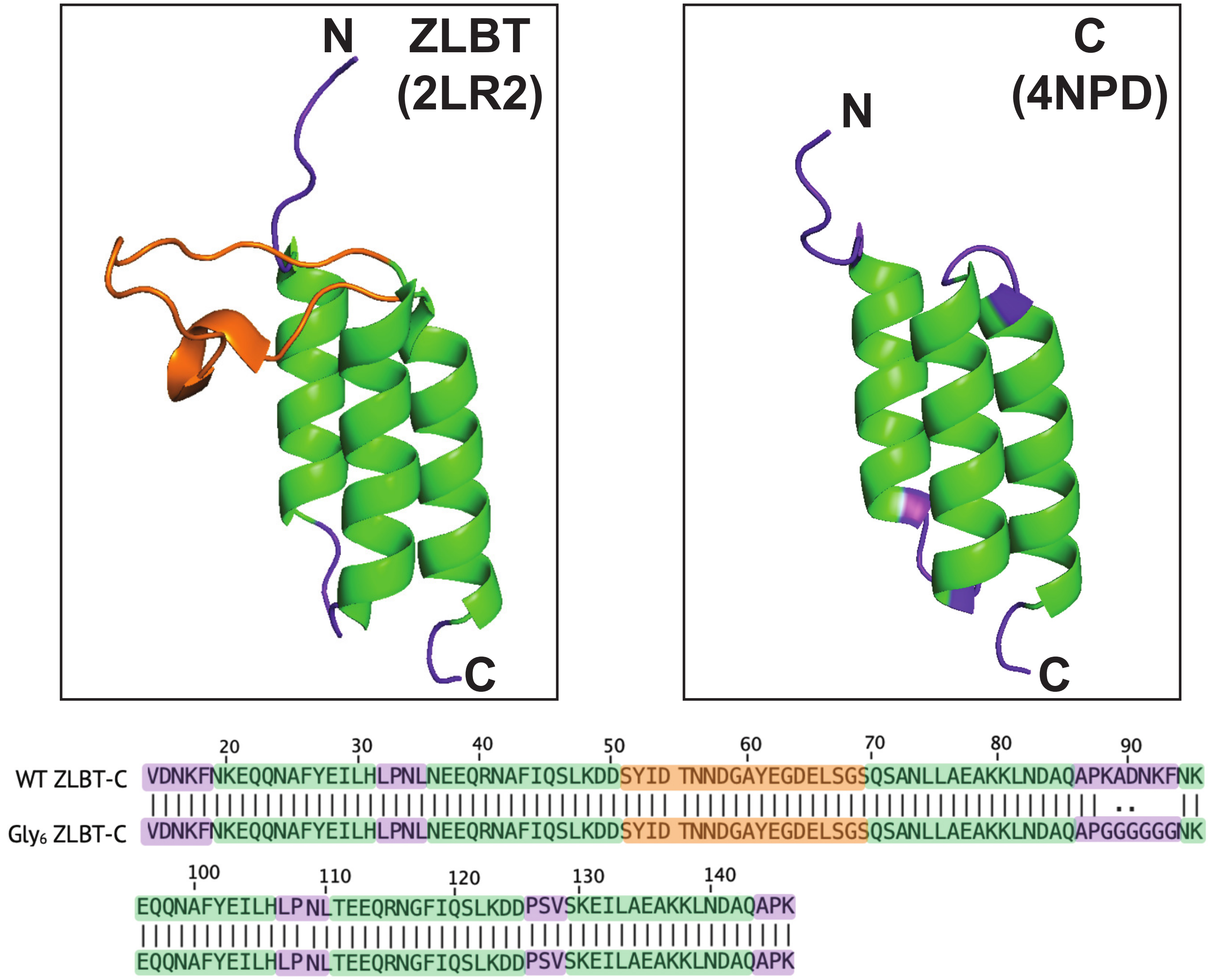


Figure 1: Structures of the ZLBT (2LR2, Model 1[2]) and C (4NPD, Alternates A[3]) domains of ZLBT-C, a two-domain protein from staphylococcal protein A[1]. In ZLBT-C, the C-terminus of the ZLBT domain is connected to the N-terminus of the C domain via a linker comprising one C-terminal residue of ZLBT ($K_{88}$) and five N-terminal residues of C ($A_{89}$-$F_{93}$). Color coding is as follows: Green = helical cores; Purple = termini, linker and inter-helical loops; Salmon = lanthanide binding tag (LBT). The sequences of the wild-type construct and a second with GGGGGG substituted for the KADNKF linker are shown below the structures. $A_{86}$, $P_{87}$, and $N_{94}$ are not helical but are also not considered part of the flexible linker because they form helix caps. NOTE: The residue numbers correspond to those given in 2LR2, followed by the residues in 4NPD. To convert from the residue numbers shown in this figure to those in 4NPD, subtract 88.

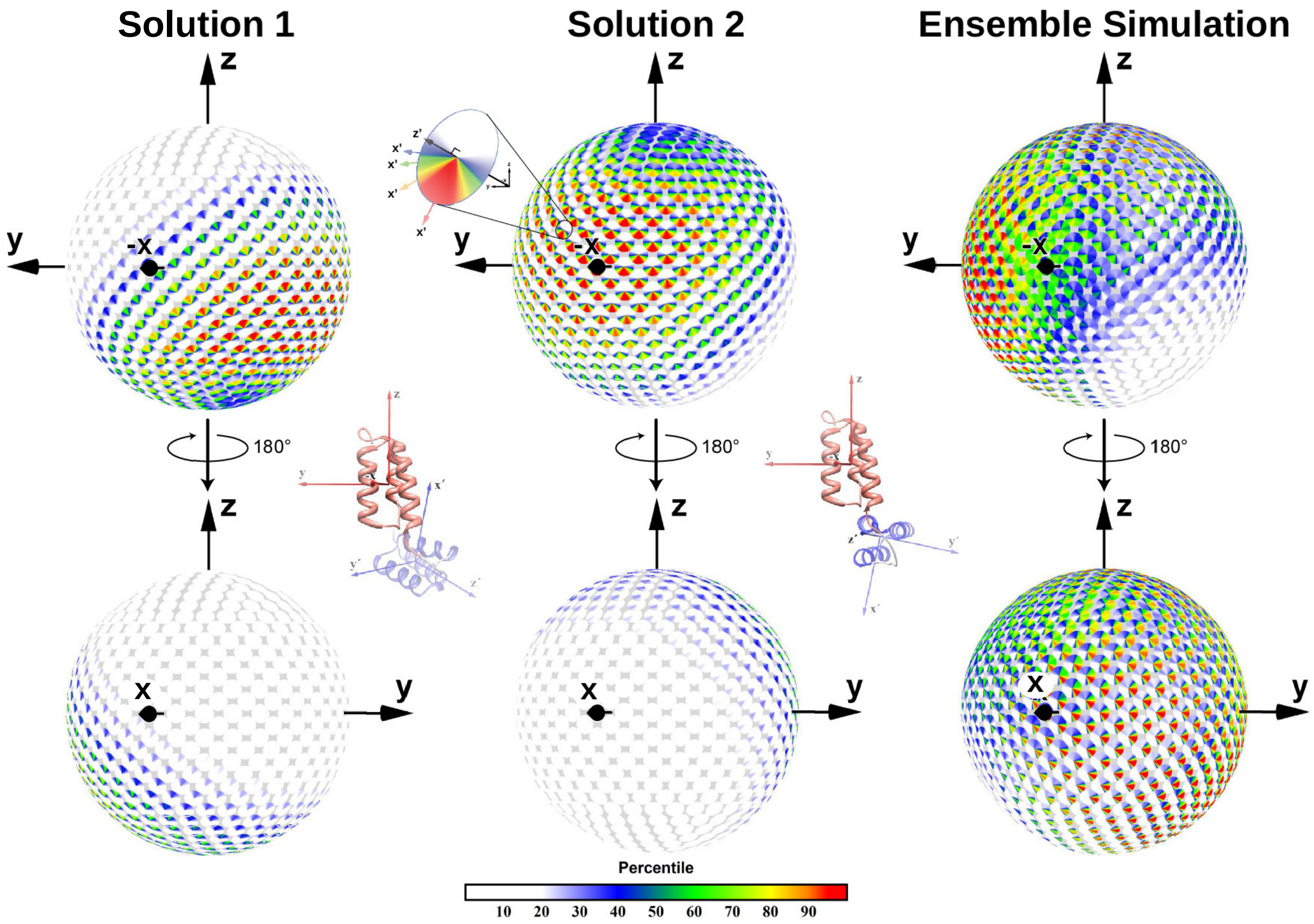


Figure 2: Disk-on-Sphere representations of two degenerate solutions to the observed NMR RDC values for ZLBT-C. The two solutions are equally probable because there is a bipolar degeneracy of the RDC observations[1]. Color-coded probabilities of the C domain's x'-axis orientation are depicted as disks whose position on the sphere corresponding to the ZLBT coordinate frame (x,y,z) represents the z'-axis of the C domain. The ribbon drawing structures represent the most probable interdomain orientation for each solution. The Ensemble Simulation is a kernalized representation of a ZLBT-C ensemble using the RanCh program[4]. The RanCh simulation suggests that Solution 2 is the correct solution because the high-probability orientations of Solution 1 are predicted to be infeasible in the RanCh simulation, likely due to steric clashes. However, predictions will be compared with both solutions, using the free energy function described by Qi, et al.[1].

### Algorithm to define the coordinate frames of the ZLBT and C domains of ZLBT-C


Terry Oas[*,1,2,], Aulane Mpouli[2], Edward Cheng[2], and Bruce Donald[*,1,2,3]

[1]Department of Biochemistry, Duke University
[2]Department of Chemistry, Duke University
[3]Department of Computer Science, Duke University
[*]Contacts:


May, 2024

1. Define the residues corresponding to the cores of Helix 2 and Helix 3 in each domain as listed below:

| Domain | Helix 2 Start | Helix 2 End | Helix 3 Start | Helix 3 End |
|---|---|---|---|---|
| ZLBT | 36 | 50 | 70 | 85 |
| C | 112 | 125 | 129 | 143 |

2. Perform the following for each domain.

3. Let $H_i$ denote helix $i$ ($i = 2, 3$).

4. Make a list the coordinates of the N, $C_\alpha$ and $C'$ backbone atoms of the Helices 2 & 3 ($H_2 \cup H_3$ list.)

5. Zero-mean the coordinates in $H_2 \cup H_3$.

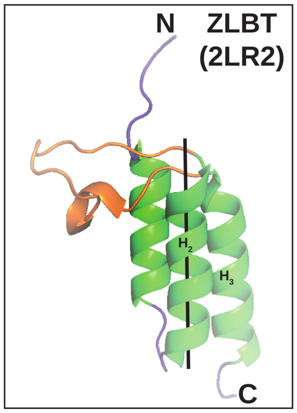


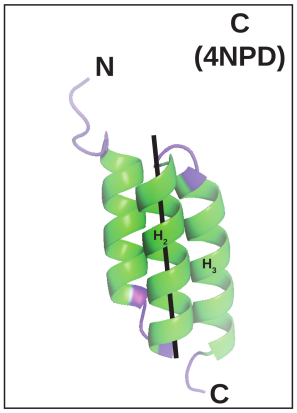


6. We will use Singular Value Decomposition (SVD) to compute the optimal (least-squares) fit of orthonormal vectors (representing a coordinate frame) to the helical core. The SVD algorithm decomposes a list of $n$ coordinates as an $n \times 3$ matrix ($\mathbf{M}$) into $\mathbf{U}\Sigma\mathbf{V^T}$, where $\mathbf{U}$ is a $n \times$N orthogonal matrix, $\Sigma$ is an $n \times 3$ diagonal matrix, and V is a $3 \times 3$ orthogonal matrix whose columns represent orthogonal axes based on the helical core coordinates. The first column of $\mathbf{V^T}$ is the unit vector parallel to the major axis of the best-fit ellipsoid (in the least-squares sense) to the helical core. The third column is the unit vector parallel to the smallest ellipsoid minor axis.

7. Perform SVD on the centered $H_2 \cup H_3$ list and take the third column of $\mathbf{V^T}$ as the $3 \times 1$ unit vector, $\overrightarrow{V_{23}}$. This vector is perpendicular to the central axes of Helices 2 & 3, points towards and bisects the two helices.

8. Make a list the coordinates of the N, $C_\alpha$ and C' backbone atoms of the Helix 2 ($H_2$ list.)

9. Zero-mean the coordinates in $H_2$.

10. Perform SVD on the centered $H_2$ list and take the first column of $\mathbf{V^T}$ as the unit vector, $\overrightarrow{V_2}$. This vector is parallel to the central axis of Helix 2 and points towards the N-terminus.

11. The axes of the helical-core-based coordinate frame are defined as shown below:

| Axis | Definition |
|---|---|
| $x'$ | $-\overrightarrow{V_2} \times \overrightarrow{V_{23}} \times \overrightarrow{V_2}$ |
| $y'$ | $\overrightarrow{V_2} \times \overrightarrow{V_{23}}$ |
| $z'$ | $-\overrightarrow{V_2}$ |
